# A sperm–oocyte protein partnership required for egg activation in *Caenorhabditis* elegans

**DOI:** 10.1101/2025.01.23.634539

**Authors:** Tatsuya Tsukamoto, Ji Kent Kwah, Mark E. Zweifel, Naomi Courtemanche, Micah D. Gearhart, Katherine M. Walstrom, Aimee Jaramillo-Lambert, David Greenstein

## Abstract

Fertilization triggers the completion of female meiosis and launches the oocyte-to-embryo transition. *C. elegans spe-11* is one of the few known paternal-effect embryonic lethal genes. We report that the sperm protein, SPE-11, forms a complex with an oocyte protein, OOPS-1 (<u>Oo</u>cyte <u>P</u>artner of <u>S</u>PE-11), and that the protein complex is required for the completion of meiosis, the block to polyspermy, and eggshell formation. Consistent with the molecular interaction of their encoded proteins, *oops-1* and *spe-11* exhibit identical null phenotypes, displaying defects in meiotic progression and cytokinesis. We show that the complex binds F-actin in the absence of other proteins and inhibits the nucleation of actin filaments *in vitro*. Thus, the OOPS-1–SPE-11 complex may function to promote F-actin-mediated meiotic cytokinesis. Both OOPS-1 and SPE-11 are intrinsically disordered proteins that are highly phosphorylated. Biochemical and genetic experiments define interactions with the protein phosphatase 1 homologs GSP-3/4, which appear to promote OOPS-1–SPE-11 function. Genetic results support a model in which the OOPS-1–SPE-11 complex interacts with the cortical EGG complex to promote meiotic cytokinesis and to activate synthesis of the eggshell.

## INTRODUCTION

In most sexually reproducing animals, oocytes arrest in meiotic prophase for a prolonged period— up to 50 years in humans. Meiosis resumes in response to hormonal signaling in the process of meiotic maturation. During oocyte meiotic maturation, the nuclear envelope of the oocyte breaks down (NEBD) in response to the activation of CDK1/Cyclin B, the maturation-promoting factor. At NEBD, microtubules gain access to the bivalents and the acentriolar meiotic spindle assembles. Fertilization triggers the process of egg activation, which results in the completion of oocyte meiosis, though the molecular mechanisms by which sperm activate embryonic development are not fully understood. Defects in oocyte meiosis and early post-fertilization development are a major cause of infertility, miscarriage, and human birth defects, and basic studies of early post- fertilization development in multiple model systems have proved informative (Gruhn and Hoffman, 2022).

The timing of the meiotic divisions with respect to fertilization varies with the species. In humans, the first meiotic division is completed before fertilization, with the second division occurring after fertilization. By contrast, in the nematode *Caenorhabditis elegans*, both meiotic divisions happen after fertilization (Albertson and Thomson, 1993; McNally and McNally, 2005). In *C. elegans*, sperm promote the completion of oocyte meiosis at several levels. The major sperm protein (MSP), which functions as the chief cytoskeletal element underlying amoeboid locomotion of nematode sperm (Italiano et al., 1996), functions as a hormone that triggers oocyte meiotic maturation and ovulation (Miller et al., 2001; Kosinski et al., 2005). During the first meiotic division, half of the homologous chromosomes are extruded in the first polar body, and half of the remaining sister chromatids are deposited in the second polar body during meiosis II. The sperm- supplied GSP-3/4 protein phosphatase 1 homologs and the GSKL-1/2 glycogen synthase kinase homologs function with the oocyte MEMI-1-3 proteins to promote meiosis II after fertilization (Ataeian et al., 2016; Banerjee and Srayko, 2022). The role of sperm-supplied factors in promoting meiosis I has been less clear. A key aspect of oocyte meiosis is that the asymmetric cell divisions that form small polar bodies and a large embryo depend on the assembly of the meiotic contractile actin ring at the cell cortex immediately adjacent to one pole of the meiotic spindle.

Many studies have focused on the role of microtubules, chromosome cohesion, and cell-cycle regulation in the control of meiotic chromosome segregation. Recent results also highlight the importance of a dynamic actin network for meiotic spindle positioning, meiotic spindle organization, chromosome clustering, proper chromosome segregation, and polar body extrusion in diverse species (Duan and Sun, 2019; Santella et al., 2020). In humans, the meiotic functions of actin have been observed to prevent aneuploidy, and maternal age-dependent decline in actin function and regulation has gained attention as a contributor to human infertility (Dunkley et al., 2022). In mice, meiotic spindle positioning to the cortex and meiotic cytokinesis require Formin2 and Spire actin nucleators to generate a dynamic cytoplasmic actin meshwork (Azoury et al., 2008; Li et al., 2008; Schuh and Ellenberg, 2008; Pfender et al., 2011; Montaville et al., 2014). In starfish, transport of chromosomes to the meiotic spindle depends on actin-based processes (Bun et al, 2018; Burdyniuk et al., 2018). Local disassembly of the actin cytoskeleton has been proposed to provide the contractile force that drives chromosome transport to the meiotic spindle. In *C. elegans*, actomyosin dynamics in the oocyte are triggered upon the onset of meiotic maturation (Yan et al., 2022). Following fertilization, actomyosin contractility and cortical destabilization have been observed to drive membrane ingressions at the cortex during the meiotic divisions (Willis et al., 2006; Fabritius et al., 2011; Schlientz and Bowerman, 2020; Quiogue et al., 2023). This global actomyosin contractility at the cortex has been proposed to provide the force that pushes the spindle through the actomyosin-free center of the constricting and ingressing contractile ring (Fabritius et al., 2011). How fertilization controls these fundamental cell and developmental events remains to be determined.

In *C. elegans,* strict paternal-effect embryonic lethal mutations in *spe-11* interfere with meiotic cytokinesis, polar body formation, synthesis of the eggshell, and the block to polyspermy (L’Hernault et al., 1988; Hill et al., 1989; McNally and McNally, 2005; Johnston et al., 2010). SPE-11 encodes a hydrophilic protein that appears to lack homologs outside of Caenorhabditid nematodes (Browning and Strome, 1996). In sperm, SPE-11 localizes to the perinuclear RNA halo (Browning and Strome, 1996), which is abnormal in *spe-11* mutants (Ward et al., 1981). RNA induces SPE-11 to undergo phase separation *in vitro*, and it has been proposed that SPE-11 associates with sperm RNAs that are delivered at fertilization (Li et al., 2023). Yet, the finding that SPE-11 can promote embryonic development when provided to the embryo through the maternal germline (Browning and Strome, 1996) suggests that its perinuclear localization in sperm may not be required. Despite SPE-11’s status as one of the few known paternal-effect mutations, its molecular interactions and activities have been inscrutable.

Here, we identify the <u>Oo</u>cyte <u>P</u>artner of <u>S</u>PE-11 (OOPS-1), which forms a complex with SPE-11 at fertilization. We show that the complex is required for multiple aspects of the egg activation process, including the completion of meiosis, eggshell formation, and the block to polyspermy. We present biochemical evidence that the OOPS-1–SPE-11 complex can function as an actin regulator. The biochemical and genetic results presented here identify molecular associations and activities of the OOPS-1–SPE-11 complex that begin to explain its essential roles.

## RESULTS

### Identification of the oocyte partner of *spe-11*

The RNA-binding proteins OMA-1 and LIN-41 control the translation of genes that play important roles during oocyte development or the oocyte-to-embryo transition (Spike et al., 2014a,b; Tsukamoto et al., 2017). The set of mRNAs that associate with OMA-1 and/or LIN-41 (n=2668, 4-fold enrichment, P<0.05) also contains many uncharacterized genes (∼50%). We screened gene knockout databases for OMA-1 and LIN-41 target genes with deletion alleles annotated as being sterile or lethal (n=90; *C. elegans* Deletion Mutant Consortium, 2012). We found one gene, *C31H1.8*, defined by the *tm6141* deletion allele, that is required in the female germline for early post-fertilization development—the completion of meiosis, eggshell formation, and the block to polyspermy—hereafter referred to as *oops-1* (<u>oo</u>cyte <u>p</u>artner of *<u>s</u>pe-11*), as described below. To confirm our initial observations with *oops-1(tm6141)*, we used genome editing to delete the entire open reading frame to generate the *oops-1(tn1898)* null allele (Fig. S1). *oops-1(tn1898)* mutant hermaphrodites or females produce all inviable embryos even when mated to wild-type males (Fig. 1C). Since *oops-1* mutant males efficiently sire progeny, the defect is on the oocyte side.

**Fig. 1.**
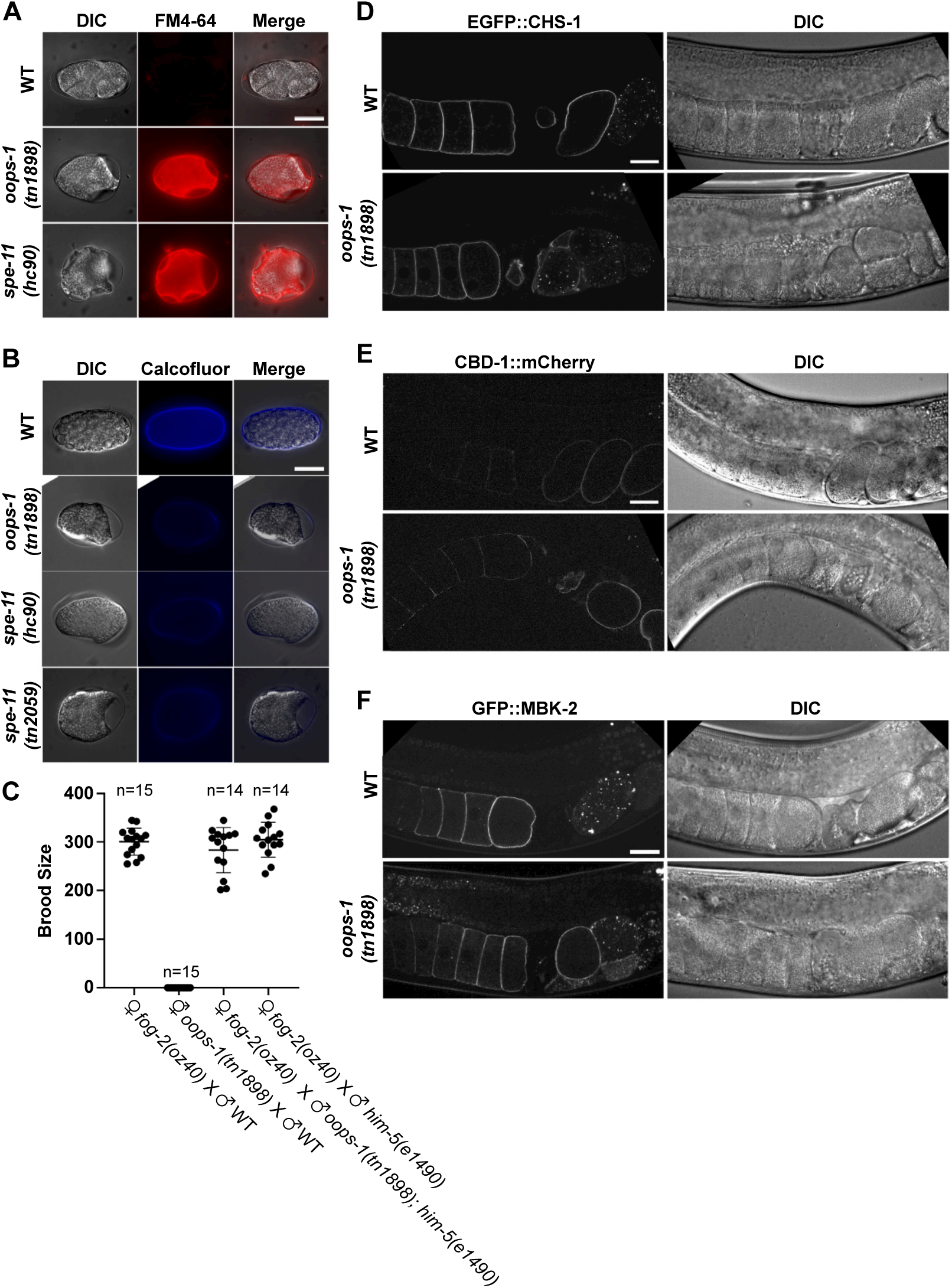
*oops-1* and *spe-11* mutant embryos exhibit a defect in chitin deposition. (A) FM4-64 staining showing the permeability layer is disrupted in mutant embryos. (B) Calcofluor staining to detect the chitin layer of the eggshell. (C) Brood size of mating assays between *oops-1(tn1898)* mutants and the wild type. (D-F) Expression of EGFP::CHS-1, CBD-1::mCherry, and GFP::MBK- 2 in wild-type and *oops-1(tn1898)* mutants. All three egg activation proteins exhibit cortical staining in oocytes and newly fertilized embryos and are subsequently internalized and degraded. Bars, 20µm.

We observed that oocytes in *oops-1(tn1898)* hermaphrodites appear to stack up in the gonad arm on the second day of adulthood (Fig. S1)—a phenotype caused by the depletion of the sperm- provided meiotic maturation signal. Since *oops-1(tn1898)* null mutants produce apparently normal numbers of sperm, we reasoned that sperm depletion might be caused by polyspermy. To assess polyspermy, *fog-2(oz40); his-72(uge30[gfp::his-72])* females, with or without the *oops-1(tn1898)* null mutation, were first treated with *mat-1(RNAi*) prior to mating. *mat-1* encodes a subunit of the anaphase-promoting complex. This treatment blocks the embryos at metaphase of meiosis I so that the sperm chromatin can be scored after mating. GFP::HIS-72 marks the oocyte-contributed chromatin (Delaney et al., 2018), enabling scoring of the unmarked paternal chromatin, which remains highly condensed, typically in the periphery distinct from the maternally contributed chromatin. We observed that one third of the embryos of *oops-1(tn1898)* mated females exhibited polyspermy, which was not observed in wild-type controls (Table 1). Consistent with the observation that the synthesis of the chitinous layer of the eggshell is required to prevent polyspermy (Johnston et al., 2010), *oops-1(tn1898)* mutant embryos are osmotically sensitive, fail to establish a permeability barrier, and exhibit a defect in chitin deposition (Fig. 1, A and B). The chitin layer of the eggshell is synthesized by CHS-1 chitin synthase, which is a component of the EGG complex that contains proteins needed for fertilization and egg activation (Maruyama et al., 2007; Parry et al., 2009). We observed that CHS-1, and the EGG complex components EGG-1-3, and MBK-2, as well as the PERM complex components, CBD-1, PERM-2, and PERM-4 (González et al., 2018), localize normally *in oops-1(tn1898)* null mutants (Figs. 1, D-F and S2).

**Table 1.** *oops-1* mutants exhibit polyspermy.

| Relevant oocyte<br>genotype<sup>b</sup> | Number of sperm observed in the embryo<sup>a</sup> |  |  |  |
| --- | --- | --- | --- | --- |
|  | 0 | 1 | 2 | 3 |
| WT (n=203) | 3% (n=6) | 97% (n=197) | 0% (n=0) | 0% (n=0) |
| *oops-1(tn1898)* (n=141) | 13.5% (n=19) | 53.2% (n=75) | 31.2% (n=44) | 2.1% (n=3) |

<sup>a</sup>Mating was to *plg-1(e2001); him-5(e1490)* males. The *plg-1* mutation was used to confirm mating via the deposition of a mating plug.
<sup>b</sup>The genotype of the females also included *fog-2(oz40); his-72(uge30[gfp::his-72])*. The females were first treated with *mat-1(RNAi)* to arrest the cell cycle post-fertilization, otherwise the maternal and paternal chromatin will replicate, which would confound the scoring of polyspermy.

### *oops-1* is required for the completion of oocyte meiosis

Because *oops-1* mutant embryos arrest at the 1-cell stage without the formation of polar bodies, we examined oocyte meiosis and early embryogenesis at high resolution using *in utero* time-lapse recordings. In these experiments, the meiotic spindles and chromatin were visualized using β- tubulin::GFP and mCherry::histone, respectively (Table 2, and Movies 1-3). To ensure phototoxicity is not an issue, we analyzed wild-type embryos, all of which completed meiosis I and II and formed polar bodies (Table 2 and Fig. 2). We observed that approximately half of the *oops-1(tn1898)* mutant embryos exhibit a phenotype in which oocyte meiotic chromosomes segregate at anaphase I and II but fail to form polar bodies (Table 2, Fig. 2, and Movie 2, referred to as the “completion phenotype”). By contrast, the other half display meiotic arrest, with the spindle drifting away from the cortex (Movie 3). The meiotic arrest phenotypes are striking. In these embryos, we observed that the meiotic spindle fails to properly shorten and maintain its association with the cortex. Instead, the spindle drifts away from the cortex and fails to maintain its structure, suggesting that the cytoskeletal forces that maintain its cortical association and integrity are deficient.

**Fig. 2.**
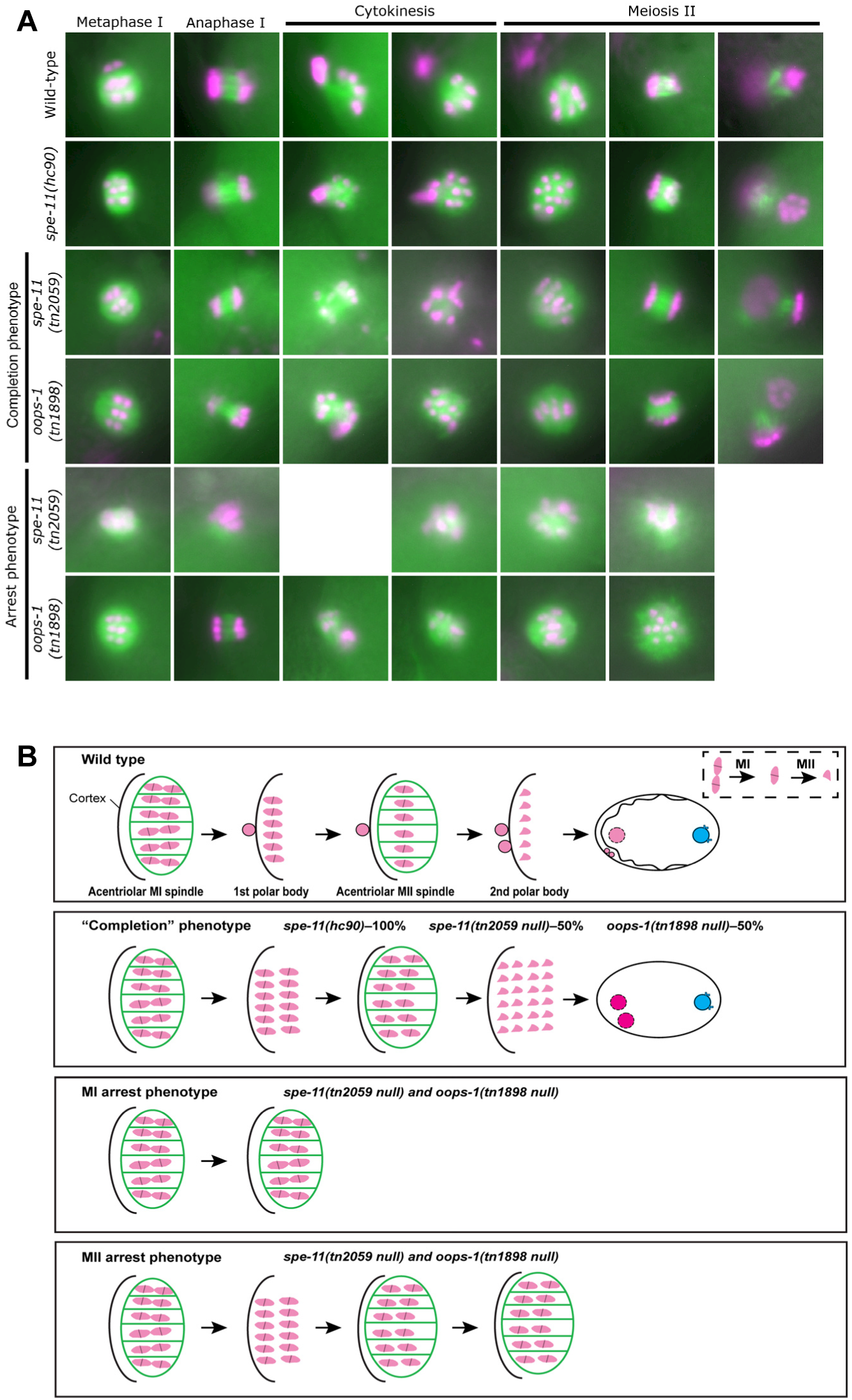
*oops-1* and *spe-11* null mutant embryos exhibit defects in the completion of oocyte meiosis. (A) Analysis of *oops-1* and *spe-11* mutants by live imaging. In the wild type, the MI and MII spindles form sequentially, chromosomes segregate and two polar bodies form. In the *spe- 11(hc90)* reduction-of-function mutant, the MI and MII spindles form and chromosomes segregate but polar bodies do not form—referred to this as the “completion phenotype.” The completion phenotype is observed in approximately 50% of *oops-1(tn1898)* and *spe-11(tn2059)* null mutants. The null mutants also exhibit arrest in MI [as in the *spe-11(tn2059)* example shown] or MII [as in the *oops-1(tn1898)* example shown]. (B) Summary of oocyte meiosis in the wild type and mutants. The inset in the top panel shows that homologs segregate in MI and sister chromatids segregate in MII. In the “completion phenotype” the internalized sets of chromosomes often form multiple nuclei. Meiotic cytokinesis (polar body formation) does not occur.

**Table 2.** Quantification of meiotic defects in *oops-1* and *spe-11* mutants.

| Genotype | Pre-anaphase I arrest | Anaphase I arrest | Meiosis II arrest | Meiotic exit<sup>a</sup> | Polar body extrusion failure<sup>b</sup> |
| --- | --- | --- | --- | --- | --- |
| WT | 0%<br>(0/20) | 0%<br>(0/20) | 0%<br>(0/20) | 100%<br>(20/20) | 0%<br>(0/20) |
| *oops-1(tn1898)* | 31.7%<br>(13/41) | 21.9%<br>(9/41) | 7.4%<br>(3/41) | 39.0%<br>(16/41) | 46.4%<br>(19/41) |
| *spe-11(hc90)* | 0%<br>(0/22) | 0%<br>(0/22) | 0%<br>(0/22) | 100%<br>(22/22) | 100%<br>(22/22) |
| *spe-11(tn2059)* | 31.6%<br>(6/19) | 10.5%<br>(2/19) | 10.5%<br>(2/19) | 47.4%<br>(9/19) | 57.9%<br>(11/19) |
| *spe-11(tn2059);*<br>*oops-1(tn1898)* | 21.1%<br>(4/19) | 36.8%<br>(7/19) | 0%<br>(0/19) | 42.1%<br>(8/19) | 42.1%<br>(8/19) |

<sup>a</sup>The mutant embryos that exit meiosis without producing polar bodies correspond to those exhibiting the “completion” phenotype (Fig. 2).
<sup>b</sup>The embryos exhibiting a failure to extrude polar bodies correspond to those exhibiting the “completion” phenotype (Meiotic exit) plus those embryos that arrest in meiosis II.

### OOPS-1 interacts with SPE-11

OOPS-1 is predicted to be an intrinsically disordered protein (IDP) using IUPRED2 (Mészáros et al., 2018) and AlphaFold (Jumper et al., 2021) with apparent homologs restricted to Caenorhabditid nematodes. To understand how OOPS-1 functions, we conducted OOPS-1 tandem-affinity purification (TAP) and mass spectrometry. Three biological replicates identified SPE-11 (Table 3 and File S1). In contrast to *oops-1*, *spe-11* mutant males produce inviable embryos even when mated to wild-type hermaphrodites or females; whereas oocytes from *spe-11* mutants generate viable and fertile progeny provided the sperm comes from wild-type males, defining *spe-11* as one of the few known strict paternal-effect lethal mutations (L’Hernault et al., 1988). We also conducted SPE-11 TAP, which recovered OOPS-1 with high efficiency (Table 3 and File S2). In six of seven of these TAPs, we isolated the nearly identical sperm-specific GSP- 3/4 serine/threonine protein phosphatases, which are homologous to human protein phosphatase 1 catalytic subunits (Chu et al., 2006; Wu et al., 2012).

**Table 3.** Tandem affinity purification of OOPS-1 and SPE-11.

| OOPS-1 TAP<sup>a</sup> |  |  |  | SPE-11 TAP<sup>b</sup> |  |  |  |
| --- | --- | --- | --- | --- | --- | --- | --- |
| Protein Coverage<sup>c</sup> (%) |  |  |  | Protein Coverage<sup>c</sup> (%) |  |  |  |
| Protein<sup>d</sup> | Exp. I | Exp. II | Exp. III | Exp. IV | Exp. V | Exp. VI | Exp. VII |
| OOPS-1 | 30.8 | 45.5 | 33.1 | 65.5 | 59.9 | 62.7 | 72.7 |
| SPE-11 | 32.4 | 54.8 | 37.1 | 78.6 | 68.9 | 85.3 | 87.6 |
| GSP-3 | 14.8 | 11.8 | 0.0 | 48.5 | 38.7 | 37.4 | 63.6 |
| GSP-4 | 14.8 | 11.8 | 0.0 | 48.5 | 38.7 | 37.4 | 63.6 |

<sup>a</sup>Experiments I–III were conducted using DG4800 *oops-1(tn1908[gfp::tev::3xflag::oops-1])* adult hermaphrodites (Fig. S1).
<sup>b</sup>Experiments IV–VII were conducted using DG5430 *spe-11(tn2094[gfp::tev::3xflag::spe-11])* animals (Fig. S4), which were adult hermaphrodites (Exps. IV and V) or a mixture of adult males and adult hermaphrodites using the DG5462 *spe-11(tn2094[gfp::tev::3xflag::spe-11]); him-5(e1490)* strain (Exps. VI and VII).
<sup>c</sup>Instrumentation was upgraded between experiments III and IV and thus the OOPS-1 and SPE-11 TAPs are not directly comparable in terms of the absolute values of the percentage protein coverage.
<sup>d</sup>Proteins shared between OOPS-1 and SPE-11 TAPs. A full list of all proteins detected is found in Files S1 and S2.

To determine whether OOPS-1 and SPE-11 can interact in the absence of other *C. elegans* proteins, we co-expressed tagged proteins in *Escherichia coli*. Affinity purification established that OOPS-1 and SPE-11 form a stable complex (Figs 3 and S3).

**Fig. 3.**
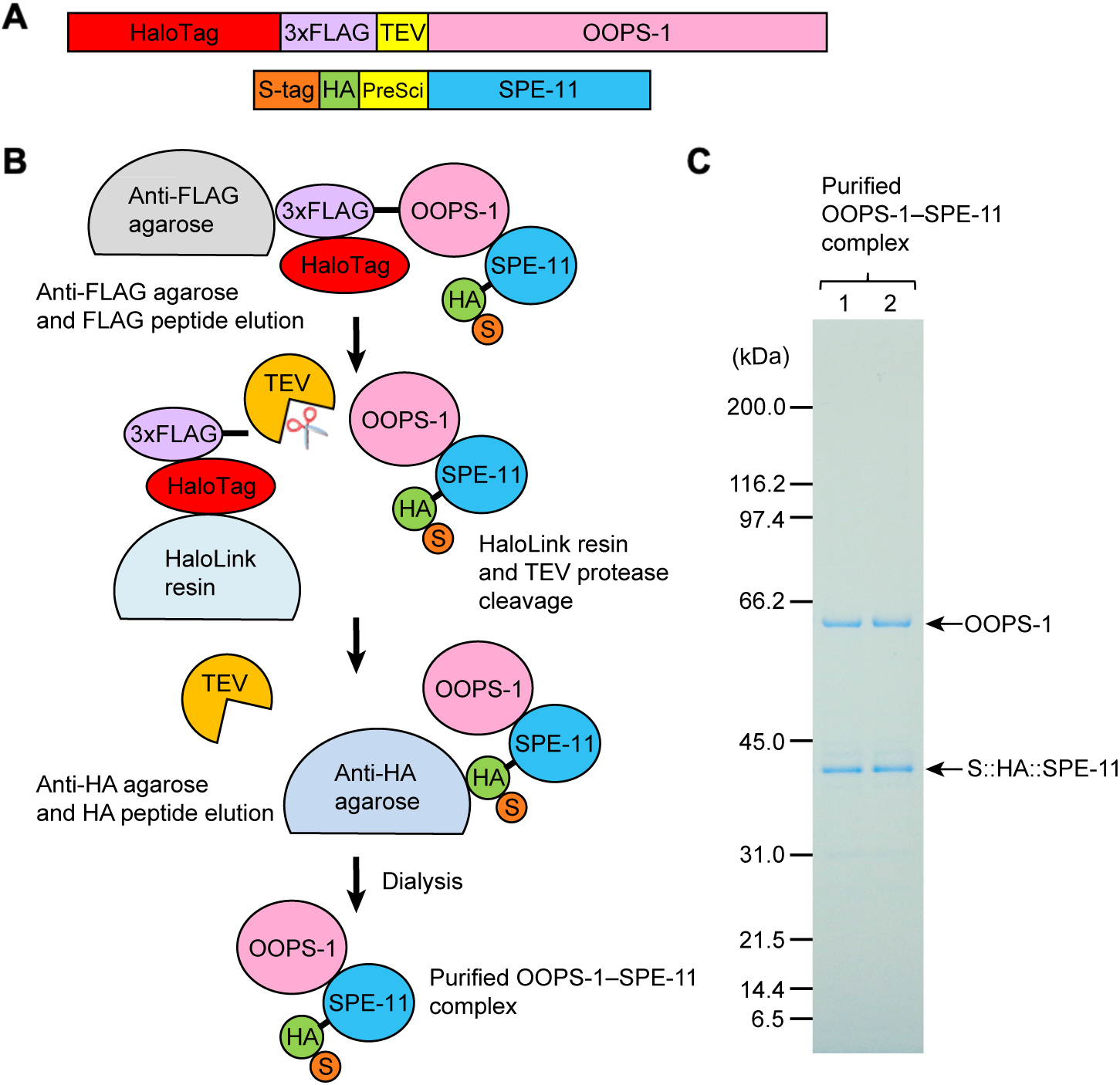
Purification of the OOPS-1–SPE-11 complex. (A) Affinity-tagged versions of OOPS-1 and SPE-11 were co-expressed in *E. coli.* (B) The scheme used to purify the complex using affinity chromatography. (C) A colloidal Coomassie-stained protein gel with two lanes of the purified complex (see Fig. S3 gel analysis of intermediate steps of the purification).

### Null mutations in *spe-11* phenocopy null mutations in *oops-1*

The meiotic phenotypes we observed for *oops-1(tn1898)* null mutants are slightly different from what was reported previously for the *spe-11(hc90)* mutation (McNally and McNally, 2005), which contains a premature termination codon at position W191 (Browning and Strome, 1996; SPE-11 has 299 amino acids). Specifically, all *spe-11(hc90)* mutant embryos observed by McNally and McNally (2005; n=5) exhibited the phenotype in which oocyte meiotic chromosomes segregate at anaphase I and II but fail to form polar bodies—the completion phenotype described above that we observed in half of the *oops-1(tn1898)* null mutants. To validate this finding, we repeated the published results for *spe-11(hc90)* and found they were correct (n=22; Table 2, Fig. 2, and Movie 4). However, prior genetic results could not exclude the possibility that *spe-11(hc90)* reduces but does not eliminate *spe-11* function (L’Hernault et al., 1988). Thus, we used genome editing to delete the entire *spe-11* open reading frame to generate the *spe-11(tn2059)* null mutation (Fig. S4). Imaging revealed that *spe-11(tn2059)* null mutants phenocopy the *oops-1(tn1898)* mutant phenotype in displaying either the completion phenotype or meiotic arrest (Table 2, Fig. 2, and Movies 5 and 6). In addition, *spe-11(tn2059); oops-1(tn1898)* double null mutants behave similarly to the individual single mutants (Table 2), suggesting that OOPS-1 and SPE-11 function together during early post-fertilization development.

### OOPS-1 and SPE-11 exhibit complementary expression patterns

Because OOPS-1 and SPE-11 are required by gametes of opposite sex, we compared their expression patterns. We generated viable and fertile N- and C-terminal fusions of OOPS-1 to GFP (Fig. S1). GFP::OOPS-1 expression is observed throughout the adult hermaphrodite germline and is enriched at the cell cortex of distal germ cells and developing oocytes; however, cortical localization becomes less apparent and expression levels decline in the most fully-grown oocytes (Figs 4 and S5). WormBase (Sternberg et al., 2024) predicts four potential isoforms (A-D), which fall into two classes: isoforms OOPS-1A/C include exons 1–3; and OOPS-1B/D start within exon 3 (Fig. S1; isoforms C and D are 2 amino acid shorter versions of isoforms A and B, respectively). Because the N-terminally tagged allele yields an expression pattern like that produced by tagging at the C-terminus (Figs 4 and S5), which would label all isoforms, OOPS-1A/C are likely the functional forms. Consistent with this, the *oops-1(gk503838)* allele, which alters the initiator methionine of OOPS-1B/D to an isoleucine, is viable and fertile (Fig. S1). To examine further whether OOPS-1B/D might be functional if made, we generated a 407-bp deletion (corresponding to 103 amino acids) in the N-terminally tagged *oops-1(tn1908)* strain to generate a new allele, *oops-1(tn1908 tn2000)*, which cannot produce isoforms OOPS-1A/C, and which tags OOPS-1B/D (Fig. S1). GFP::OOPS-1B/D localizes to the cytoplasm and is abundant in oocytes but lacks the cortical enrichment observed in the germline in OOPS-1A/C (Fig. S5). Thus, an N-terminal domain within OOPS-1A/C is responsible for cortical tethering in the germline and potentially for limiting OOPS-1 levels. This cortical tethering domain appears to be dispensable for post- fertilization development, with the caveat that GFP::OOPS-1B/D is expressed at high levels. Similarly, we generated viable and fertile alleles of SPE-11 tagged at the N-terminus with mScarlet or GFP (Fig. S4). In the adult hermaphrodite gonad, the only cells observed to express mScarlet::SPE-11 are sperm (Fig. 4). Thus, the OOPS-1–SPE-11 complex likely forms and functions upon fertilization.

**Fig. 4.**
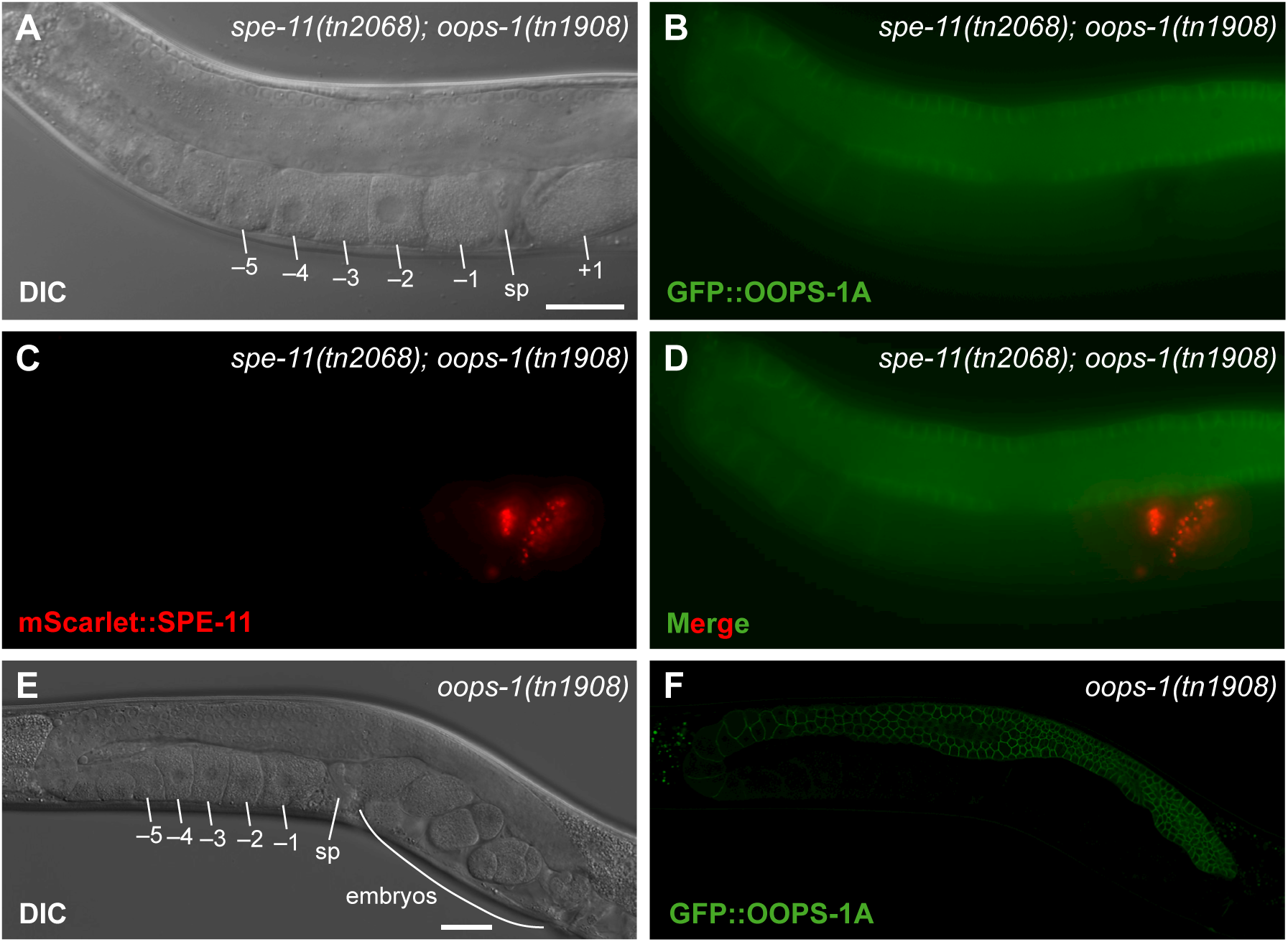
**OOPS-1 and SPE-11 exhibit complementary expression patterns**. (A-D) DIC (A) and wide-field fluorescence micrographs (B-D) of a *spe-11(tn2068[mScarlet::tev::3xflag::spe-11]); oops-1(tn1908[gfp::tev::3xflag::oops-1a])* adult hermaphrodite expressing GFP::OOPS-1A (green) and mScarlet::SPE-11 (red). OOPS-1A is expressed throughout the female germline and becomes downregulated in oocytes. SPE-11 expression is restricted to sperm. The proteins encounter each other upon fertilization but are at low levels. DIC (E) and a single optical section of a confocal fluorescence micrograph (F) of an *oops-1(tn1908[gfp::tev::3xflag::oops-1a]* adult hermaphrodite high-lighting the cortical localization of GFP::OOPS-1A in the distal germline and its down-regulation in proximal oocytes. Proximal oocytes (–1 to –5), the spermatheca (sp), and a newly fertilized embryo (+1) and older embyos in the uterus are indicated. Bars, 30 μm.

### SPE-11 can function when expressed in the oocyte, but OOPS-1 cannot function efficiently when expressed in sperm

Browning and Strome (1996) expressed the *spe-11* cDNA during oogenesis using a heat shock promoter and observed transient rescue of fertility in *spe-11(hc90)* mutants. Using the tools available at the time, it was not possible to quantitatively evaluate the efficiency of the rescue. We generated two independent single-copy insertions (*tnSi5* and *tnSi6*) in which mScarlet::SPE-11 is expressed under control of the *mex-5* promoter and the *spe-11 3’UTR* (Fig. 5A). We observed that *tnSi5* and *tnSi6* were able to rescue *spe-11(tn2059)* null mutants to fertility; however, these constructs expressed the mScarlet::SPE-11 protein in both oocytes and sperm (Fig. 5C). To test whether mScarlet::SPE-11 expression exclusively in the female germline rescues the *spe-11* mutant phenotype, we used the *fog-2(oz40)* mutation to feminize the germline and mated the *tnSi5; fog-2(oz40)* and *tnSi6; fog-2(oz40)* females with *spe-11(tn2059)* null mutant males. We observed 96% and 98% embryonic viability, indicating that SPE-11 can efficiently mediate post-fertilization development when delivered through the oocyte (Table 4). By contrast, viable cross progeny were never observed when *fog-2(oz40)* females were crossed with *spe-11(tn2059)* null mutant males (Table 4). We were also able to rescue *spe-11(tn2059)* hermaphrodites to fertility through the expression of GFP::SPE-11 in oocytes using the *oma-1*, *oma-2*, *rme-2*, and *puf-5* promoters and their respective 3’UTRs (*tnSi21*, *tnSi23*, *tnSi25* and *tnSi27*, respectively), further providing evidence that provision of SPE-11 by the oocyte is sufficient.

**Fig. 5.**
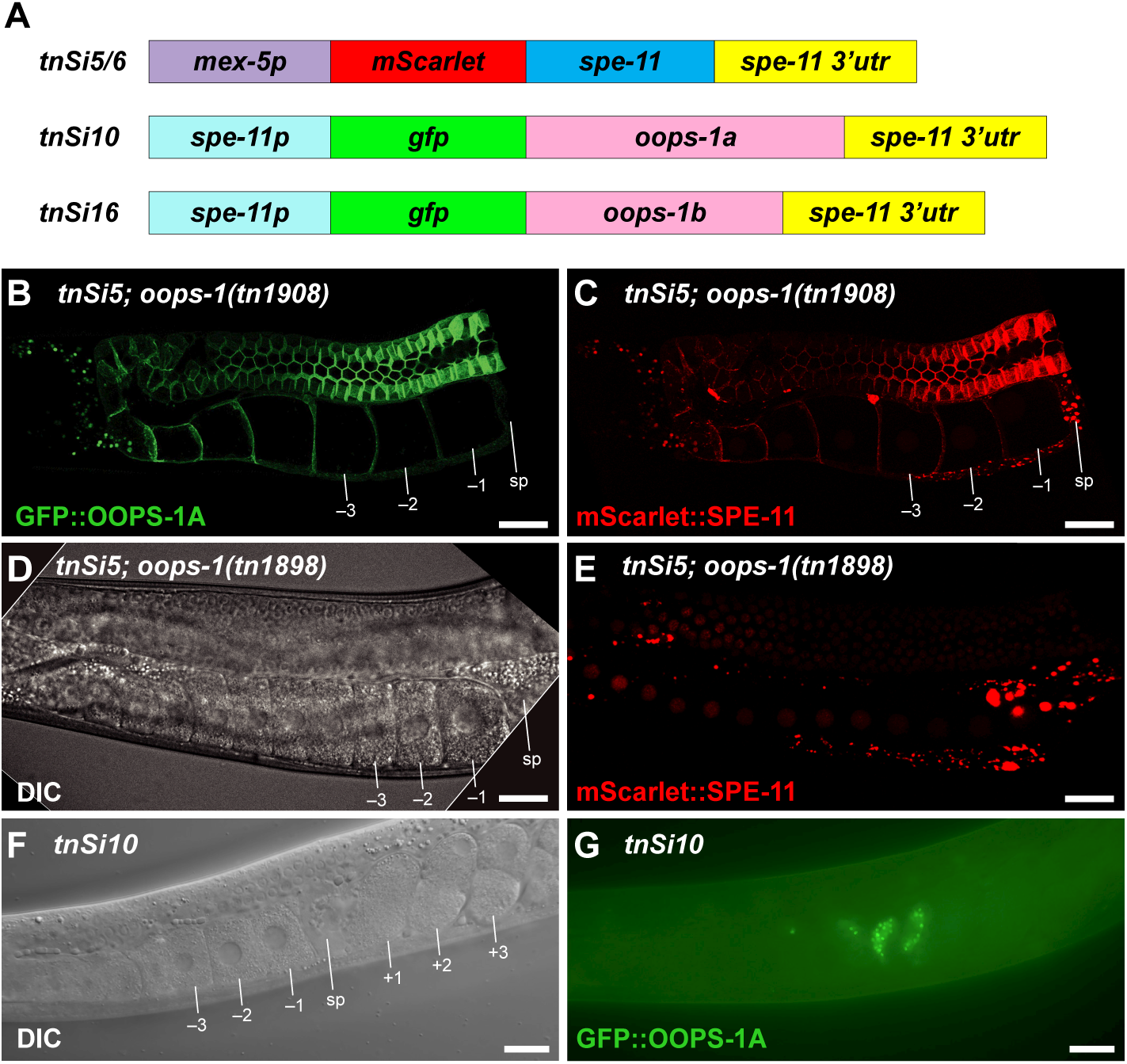
Ectopic expression of SPE-11 in the female germline and OOPS-1A in the male germline. Single-copy insertion constructs to express SPE-11 in the female germline and OOPS- 1A and OOPS-1B in the male germline. (B, C) Fluorescence micrograph of a *tnSi5; oops- 1(tn1908[gfp::tev::3xflag::oops-1a])* adult hermaphrodite expressing GFP::OOPS-1A (B) and mScarlet::SPE-11 (C). Note, mScarlet::SPE-11 and GFP::OOPS-1A colocalize and are prominent at the cortex of germ cells and oocytes. (D, E) DIC (D) and fluorescence micrograph (E) of a *tnSi5; oops-1(tn1898* null) adult hermaphrodite. Note, mScarlet::SPE-11 exhibits nuclear localization in the absence of OOPS-1. Thus, the cortical localization of mScarlet::SPE-11 observed when SPE- 11 is ectopically expressed in the female germline is dependent on OOPS-1. (F, G) DIC (F) and fluorescence micrographs (G) of a *tnSi10* adult hermaphrodite expressing GFP::OOPS-1A in sperm. Bars, 20 µm.

**Table 4.** SPE-11 can function via the oocyte, but provision of OOPS-1 by sperm is unable to fully support development.

| Female | Male | % Viability |
| --- | --- | --- |
| *fog-2(oz40)* | *spe-11(tn2059); him-5(e1490)* | 0 (n=9473) |
| *tnSi5<sup>a</sup>; fog-2(oz40)* | *spe-11(tn2059); him-5(e1490)* | 96.3 (n=28,750) |
| *tnSi6<sup>b</sup>; fog-2(oz40)* | *spe-11(tn2059); him-5(e1490)* | 98.0 (n=31,126) |
| *tnSi5<sup>a</sup>; fog-2(oz40)* | WT | 97.5 (n=28,985) |
| *tnSi6<sup>b</sup>; fog-2(oz40)* | WT | 97.9 (n=27,149) |
| *fog-2(oz40)* | WT | 99.4 (n=32,760) |
| *fog-2(oz40)* | *tnSi10<sup>c</sup>; him-5(e1490)* | 100 (n>5000) |
| *oops-1(tn1898); fog-2(oz40)* | *tnSi10<sup>c</sup>; him-5(e1490)* | 0 (n>5000) |
| *fog-2(oz40)* | *tnSi16<sup>d</sup>; him-5(e1490)* | 100 (n>5000) |
| *oops-1(tn1898); fog-2(oz40)* | *tnSi16<sup>d</sup>; him-5(e1490)* | 0.5 (n~28,180) |

<sup>a</sup>*tnSi5[mex-5p::mScarlet::spe-11::spe-11 3'UTR]*
<sup>b</sup>*tnSi6[mex-5p::mScarlet::spe-11::spe-11 3'UTR]*
<sup>c</sup>*tnSi10[spe-11p::GFP::oops-1a::spe-11 3'UTR]*
<sup>d</sup>*tnSi16[spe-11p::GFP::oops-1b::spe-11 3'UTR]*

When mScarlet::SPE-11 is expressed in the female germline, we observed colocalization with GFP::OOPS-1 at the cortex of germ cells and oocytes (Fig. 5, B and C). This cortical localization is not observed in *oops-1(tn1898)* null mutants, and instead mScarlet::SPE-11 exhibited nuclear localization (Fig. 5, D and E). Thus, OOPS-1 and SPE-11 directly interact *in vivo*.

To determine whether OOPS-1 can function when delivered by the sperm, we generated single- copy insertions to express GFP::OOPS-1A (*tnSi10*) or GFP::OOPS-1B (*tnSi16*) in the male germline under control of the *spe-11* promoter and *3’UTR* (Fig. 5A). We tested the ability of *tnSi10* and *tnSi16* males to rescue *oops-1(tn1898); fog-2(oz40)* females upon mating and observed no rescuing activity for *tnSi10* and only very weak activity for *tnSi16* (∼0.5% embryonic viability; Table 4). In addition to using the *spe-11* promoter (Fig. 5, F and G), we also used the sperm- specific *peel-1* and *trp-3* promoters and their respective *3’UTR* elements to express GFP::OOPS- 1A in sperm (*tnSi12* and *tnSi14*, respectively) and were also unable to rescue *oops-1(tn1898)* mutants to fertility.

### Protein domains required for OOPS-1 and SPE-11 function

Individually, OOPS-1 and SPE-11 are predicted by AlphaFold to be largely unstructured. IDPs have frequently been found to undergo a disorder-to-order transition upon binding their partners (Trivedi and Nagarajaram, 2022). To determine the protein regions essential for OOPS-1 and SPE-11 function, we generated in-frame deletions in the context of the N-terminally tagged GFP::OOPS-1 and GFP::SPE-11 (Fig. 6). This analysis identified a 211 amino acid region from the central portion of OOPS-1 as being required for function (Fig. 6, A and B). We combined 5’ and 3’ deletions to generate a strain expressing GFP::OOPS-1(139-349) Mini (Fig. 6C), which was viable and fertile, indicating that this central region of OOPS-1 is necessary and sufficient for function; however, GFP::OOPS-1 Mini is expressed at high levels in the germline (Fig. S5). For SPE-11, multiple regions are required for function (Fig. 6, D and E). Using AlphaFold Multimer, both proteins are predicted to exhibit structurally ordered regions with high per residue measures of local confidence using the predicted local distance difference test (pLDDT; Fig. S6).

**Fig. 6.**
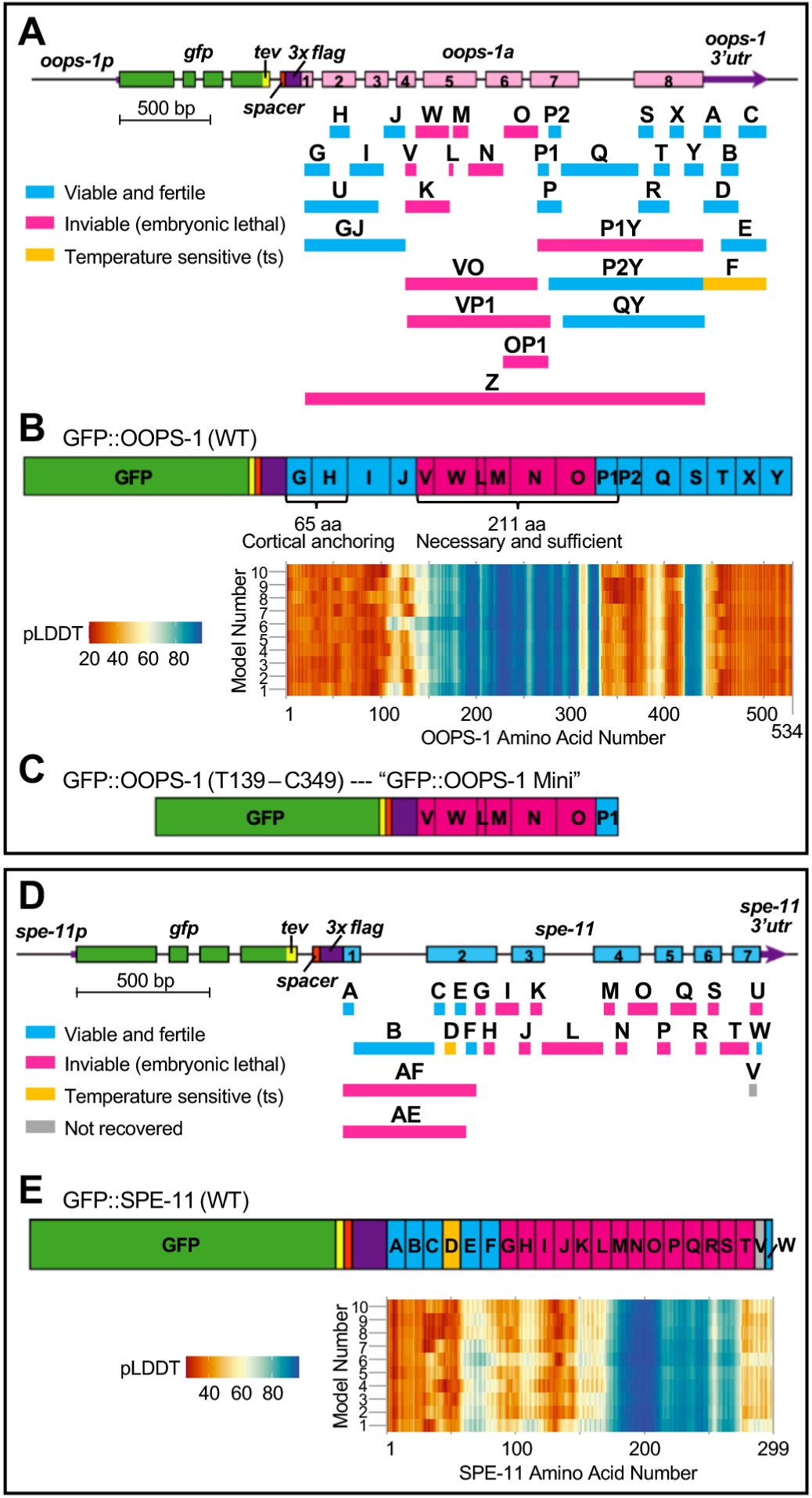
Protein domains required for OOPS-1 and SPE-11 function. (A, D) Deletions made by CRISPR-Cas9 genome editing in the context of *oops-1(tn1908[gfp::tev::3xflag::oops-1a])* (A) and *spe-11(tn2094[gfp::tev::3xflag::spe-11])* (D). (B, E) Essential and non-essential regions of OOPS-1 (B) and SPE-11 (E) mapped onto a diagram of each protein. Structures for the OOPS-1 and SPE-11 complex were modeled using the AlphaFold 3 server (Fig. S6). The pLDDT values for the Ca for each amino acid were plotted in a heatmap. (C) GFP::OOPS-1 Mini made by combining N-terminal and C-terminal deletions confers fertility and viability.

Remarkably, the regions of high pLDDT values correspond well to the essential regions of OOPS- 1 and SPE-11 (Fig. 6, B and E), suggesting these regions might mediate their interaction. For SPE- 11, we observed that a more N-terminal 14 amino acid deletion (deletion D in Fig. 6D) confers temperature sensitivity with an average brood size of 2 ± 3 (n=43) at 25°C. This mutation, hereafter referred to as *spe-11(*ts), might disrupt interactions with other proteins needed for post-fertilization development.

### SPE-11 genetically interacts with components of the EGG complex

We conducted a genetic selection for dominant suppressors of *spe-11(*ts) and isolated mutations that substantially increase fertility (Table 5 and Fig. S7). Since the *spe-11(*ts) allele is marked by *gfp*, we were able to determine that the suppressor mutations do not alter GFP::SPE-11(ts) expression levels. Seven of these rare dominant suppressors were backcrossed to remove unlinked mutations to facilitate mutation identification using whole-genome sequencing (WGS; File S3). These mutations define at least three loci (*gsp-3*, *chs-1*, and *egg-3*; Table 5). Five additional mutations were identified by genetic mapping and Sanger sequencing the three candidate suppressor loci. The isolation of an allele of *gsp-3* as a *spe-11(*ts) suppressor is consistent with the identification of GSP-3/4 as a protein found in OOPS-1 and SPE-11 TAPs (Table 3). The P195S mutation in *gsp-3(tn2202)* affects an amino acid that is conserved in both GSP-4 and the human PP1B homolog and is predicted to be in a surface-exposed loop (Fig. S8). CHS-1, chitin synthase, is activated upon fertilization to generate the chitin layer of the eggshell (Zhang et al., 2005; Olson et al., 2012; Stein and Golden, 2018), which fails to form in *oops-1* and *spe-11* mutants. CHS-1 and EGG-3 are required for the formation of polar bodies and share mutant phenotypes with *spe- 11* and *oops-1* (Maruyama et al., 2007; Johnston et al., 2006, 2010; González et al., 2018; Kawasaki et al., 2024). The suppressor mutations in CHS-1 are predicted to be intracellular (Fig. S9). The strong suppression of *spe-11(*ts) by the *chs-1* mutations suggests they might restore interactions with a defective SPE-11 (Fig. S7). The two *egg-3* suppressor mutations affect adjacent amino acids, predicted to localize to a surface-exposed loop (Fig. S10).

**Table 5.** Dominant *spe-11(ts)* suppressor mutations.

| Dominant suppressor loci<sup>a</sup> | Function | Amino acid substitutions (*alleles*)<sup>b</sup> |
| --- | --- | --- |
| *chs-1* | Chitin Synthase, required for eggshell formation, the block to polyspermy, and the completion of meiosis | S347F(*tn2248*), R645K(*tn2195*), **L669F(*tn2201*)**, A684V(*tn2242*), **A827T(*tn2191*)**, **A1114V(*tn2189*, *tn2198*, *tn2210*)**, R1308C(*tn2193*) |
| *gsp-3* | Sperm-specific PP1 phosphatase | **P195S(*tn2202*)** |
| *egg-3* | EGG-3 required for eggshell formation, the block to polyspermy, and the completion of meiosis | **P197S(*tn2205*)**, G198R(*tn2190*) |

<sup>a</sup>26 additional suppressor mutations have not been assigned.
<sup>b</sup>Mutations shown in bold font were ascertained by WGS and Sanger sequencing (File S3).
Mutations shown in regular font were ascertained by Sanger sequencing.

### OOPS-1 and SPE-11 are phosphoproteins

The data presented suggest that the GSP-3/4 phosphatases might promote the function of the OOPS-1–SPE-11 complex. Although there are many potential forms such a regulatory mechanism might take, we considered the possibility that the OOPS-1–SPE-11 complex might be controlled by protein phosphorylation. Thus, we examined the mass spectrometry spectra from the TAP experiments for instances of protein phosphorylation. We found multiple examples in which phosphorylation was supported at high-confidence levels by multiple diagnostic b- and y-type fragment ions in the mass spectra (Fig. 7 and File S4). These phosphorylation sites are in non- essential regions of both proteins (Fig. 6), suggesting they might define activating and/or inhibiting regulatory modifications.

**Fig. 7.**
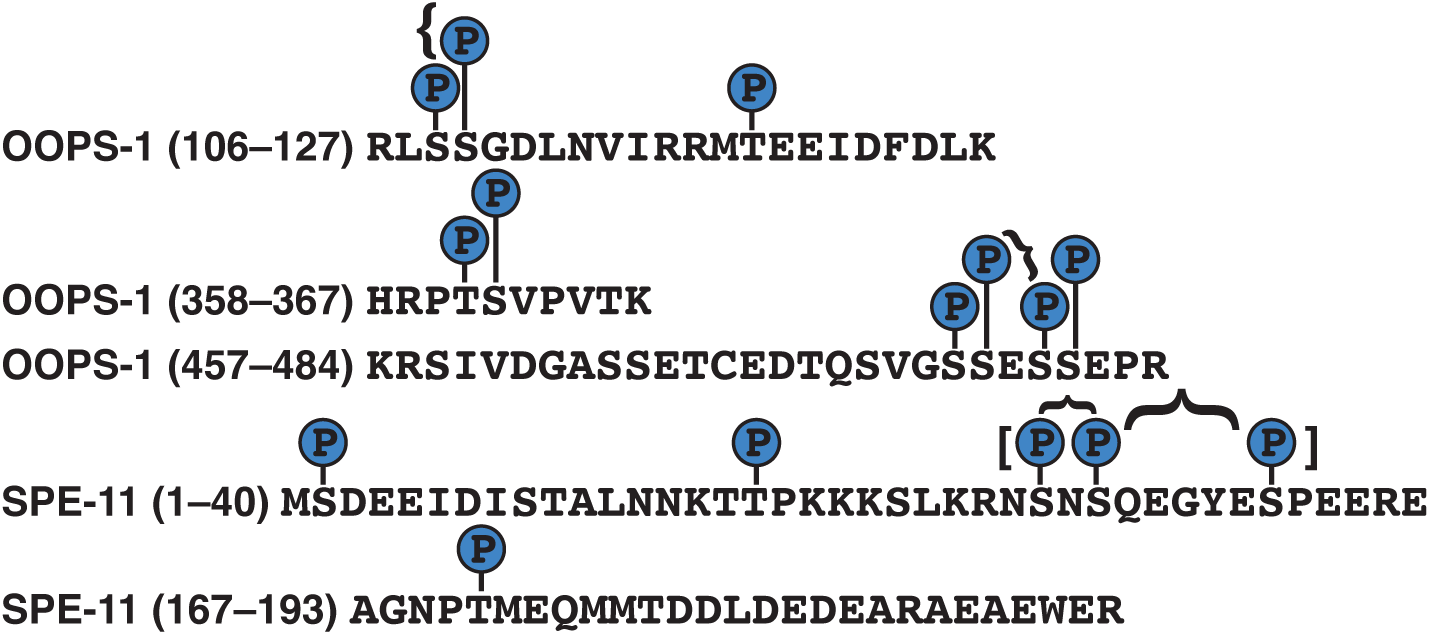
OOPS-1 and SPE-11 are phosphoproteins. High-confidence phosphorylation sites identified by mass spectrometry in OOPS-1 and SPE-11, supported by multiple diagnostic b- and y-type fragment ions (see File S4). Doubly (braces) and triply (brackets) phosphorylated sites are indicated.

### The OOPS-1–SPE-11 complex functions as an actin regulator *in vitro*

*spe-11* and *oops-1* mutant embryos are defective in meiotic cytokinesis and exhibit similarities to embryos treated with the actin inhibitor Latrunculin A (Yang et al., 2003; McNally and McNally, 2005). Thus, we investigated whether the purified complex interacts with filamentous actin (F- actin; Fig. 8). We initiated actin polymerization in the presence and absence of the OOPS-1–SPE- 11 complex and separated F-actin from G-actin by centrifugation at 100,000 x g (Fig. 8, A and B). Ordinarily, the OOPS-1–SPE-11 complex localizes to the supernatant fraction after centrifugation [Fig. 8B, duplicate reactions (Rxns) 2 and 3]; however, when F-actin is present, the complex co- sediments with F-actin (Fig. 8B, Rxns 4 and 5). This result suggests that the OOPS-1–SPE-11 complex binds F-actin in the absence of other proteins. We also observed a reduction in the concentration of pelleted actin in the presence of the complex (Fig. 8B, cf Rxn1 with Rxns 4 and 5), consistent with the possibility that the complex decreases the rate of spontaneous actin polymerization, potentially by sequestering actin monomers. To examine the rate of spontaneous actin polymerization, we monitored the time course of actin assembly in the presence of a range of concentrations of the OOPS-1–SPE-11 complex *in vitro* (Fig. 8C). In these assays, filament nucleation is the primary determinant of the polymerization rate. We observed a concentration- dependent decrease in the rate of actin assembly in the presence of the OOPS-1–SPE-11 complex. Reactions containing OOPS-1–SPE-11 also attained lower final fluorescence values than did the reaction containing actin alone. Collectively, these results suggest that the OOPS-1–SPE-11 complex either slows filament nucleation, decreases the total concentration of F-actin that can be obtained through polymerization, or both. To dissect the mechanism by which the complex influences polymerization, we examined the effects of the OOPS-1–SPE-11 complex on F-actin assembly in the presence of a constitutively active formin. Formins stimulate polymerization by speeding filament nucleation and regulating filament elongation. In bulk assembly assays, the nucleation activity of formin gives rise to a dramatic increase in the rate of actin polymerization (Pruyne et al., 2002; Pring et al., 2003). Consistent with this activity, formin-mediated actin assembly attains equilibrium within 5 min in the absence of the OOPS-1–SPE-11 complex (Fig. 8D). Inclusion of the complex in the reactions slows actin assembly in a dose-dependent fashion. In contrast, the fluorescence signal measured at the end of polymerization is unaffected by inclusion of OOPS-1–SPE-11. This supports a mechanism in which the complex inhibits filament nucleation rather than decreasing the total concentration of polymerized actin generated in the reaction. Application of a linear fit to the time courses at the point where half of the actin is polymerized reveals a ∼60% decrease in the rate of filament nucleation at the highest concentration of complex we sampled. Remarkably, inhibition of actin polymerization occurs at sub- stoichiometric concentrations of the OOPS-1–SPE-11 complex: ∼45–180-fold below the concentration of actin monomers. Thus, inhibition cannot involve “monomer sequestration,” but might occur through direct binding of actin nuclei, as these species are present at low concentrations throughout the polymerization reaction. These observations indicate that the complex can function as a potent actin regulator, even when present at limiting concentrations.

**Fig. 8.**
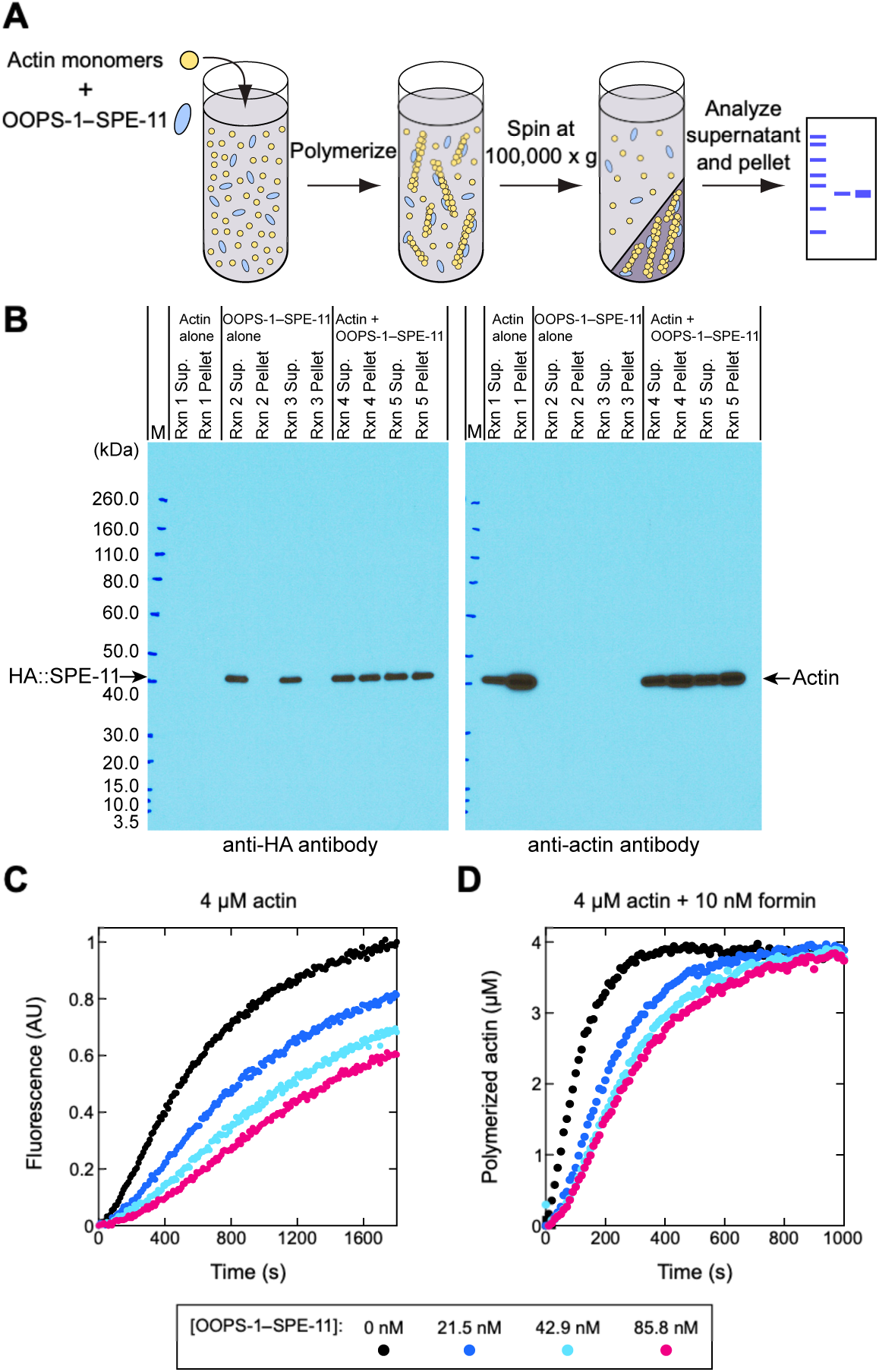
Biochemical evidence that the OOPS-1–SPE-11 complex is an actin regulator. (A, B) The OOPS-1–SPE-11 complex interacts with F-actin in the absence of other proteins. (A) Schematic representation of the co-sedimentation assay used to assess the interaction of the OOPS- 1–SPE-11 complex with F-actin. (B) Western blot analyses of the supernatant and pellet fractions. Filaments were assembled from 4 µM actin monomers in the presence or absence of 223 nM OOPS-1–SPE-11 complex. (C) The OOPS-1–SPE-11 complex slows actin polymerization. Representative time courses of spontaneous polymerization of 4 µM actin (20% pyrene-labeled) in the presence of a range of concentrations of OOPS-1–SPE-11. (D) The OOPS-1–SPE-11 complex inhibits formin-mediated actin assembly. Time courses of polymerization of 4 µM actin (20% pyrene-labeled) in the presence of 10 nM formin and a range of concentrations of OOPS-1– SPE-11. Fluorescence data were normalized and are represented in units of polymerized actin, which facilitates quantification of polymerization rates.

## DISCUSSION

In many organisms, early development is under substantial maternal control. By contrast, the mechanisms by which the sperm promotes embryonic development, beyond contributing a haploid genome and a centriole pair, are less understood. Here, we address the molecular mechanisms of one of the few strict paternal-effect genes known, *C. elegans spe-11*. We show that SPE-11 functions with an oocyte protein, OOPS-1, in a protein complex that forms at fertilization. The paternal and maternal genetic requirements for *spe-11* and *oops-1*, respectively, indicate that the protein complex is required for completion of oocyte meiosis, meiotic cytokinesis, synthesis of the eggshell, the block to polyspermy, and the embryonic divisions.

OOPS-1 and SPE-11 are IDPs that appear to be restricted to Caenorhabditid nematodes. This observation is unsurprising because virtually all reproductive proteins are sexually selected, and “the most interesting examples are likely those that directly bind molecules derived from the other sex (Wilburn and Swanson, 2016).” A single sperm, with ∼1/10,000th the volume of an oocyte, deposits a payload of SPE-11, which must rapidly interact with OOPS-1 to execute meiosis at the anterior cortex. Evolution must have selected for a high-affinity interaction because the eggshell begins to form within approximately 5 min following fertilization and oocyte meiosis is completed within approximately 30 min after fertilization (Ward and Carrel, 1979; McCarter et al., 1999). Indeed, our TAP experiments efficiently recovered the protein complex even though these proteins appear to be found in the same cell only after fertilization. IDPs can exhibit diffusion-limited binding (Borgia et al., 2018), which might be a conserved aspect of fertilization. The purification of the complex from *E. coli* demonstrates that the two proteins can interact in the absence of other *C. elegans* proteins.

In the context of the *C. elegans* embryo, interactions with other proteins might facilitate the formation or function of the OOPS-1–SPE-11 complex. Consistent with this possibility, we recovered dominant mutant alleles of *chs-1* chitin synthase and *egg-3* in a screen for suppressors of a *spe-11(*ts) mutation. As mentioned above, CHS-1 and EGG-3 are required for the formation of polar bodies and share mutant phenotypes with *spe-11* and *oops-1,* including defects in the completion of oocyte meiosis and eggshell synthesis. The fact that disruptions in eggshell synthesis perturb polar body formation was somewhat enigmatic because it was hard to imagine how events happening external to the plasma membrane might affect meiotic cytokinesis on the other side— the chitin layer of the eggshell is deposited external to the plasma membrane just beneath the vitelline layer (Stein and Golden, 2018). Chitin synthases are localized to the plasma membrane and thought to act as channels that export the chitin polymer across the plasma membrane (Ren et al., 2022). Our findings suggest a model in which CHS-1, or its association with the EGG complex, might localize the OOPS-1–SPE-11 complex to the cortex to spatially control the regulation of actin dynamics for the completion of meiosis. In turn, the OOPS-1–SPE-11 complex might activate CHS-1 for eggshell synthesis (Fig. 9). Consistent with this model, CHS-1 localization is normal in *oops-1* and *spe-11* mutants, but they lack chitin deposition. One possibility is that the rare dominant *spe-11(*ts) suppressor mutations restore molecular interactions between the EGG complex and the OOPS-1–SPE-11 complex.

**Fig. 9.**
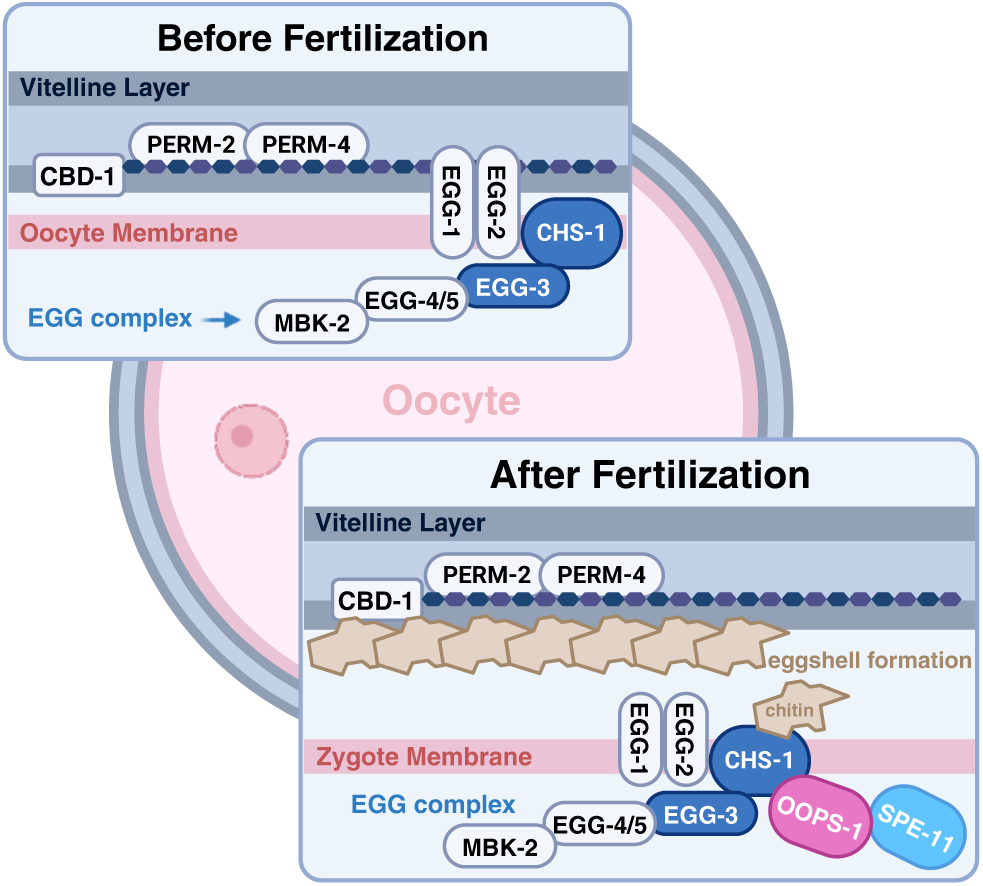
Cortical recruitment model. Components of the EGG complex are proposed to recruit the OOPS-1–SPE-11 complex to the cortex for the completion of oocyte meiosis. In turn, OOPS-1– SPE-11 are required for synthesis of the chitin layer of the eggshell.

Biochemical experiments reported here suggest that the OOPS-1–SPE-11 complex can function as an actin regulator. Specifically, we found that the OOPS-1–SPE-11 complex interacts with F-actin in the absence of other proteins and inhibits formin-mediated actin assembly at sub- stochiometric concentrations. These data suggest that the OOPS-1–SPE-11 complex possesses an elemental biochemical function at the level of the actin filament. These biochemical experiments were motivated partly by the fact that meiotic cytokinesis is an actin-based process, which enables the asymmetric division that forms small polar bodies and a large embryo. Although many studies have examined the role of the actin cytoskeleton during polarization of the *C. elegans* embryo at the pronuclear stage (Munro and Bowerman, 2009; Gubieda et al., 2020), there have been far fewer studies of actin dynamics just after fertilization because of technical challenges—newly fertilized embryos are osmotically sensitive before the eggshell forms, requiring the imaging of thick samples *in utero*. Indeed, both *spe-11* and *oops-1* mutant embryos exhibit osmotic sensitivity.

Actin-based processes, such as meiotic cytokinesis, which is defective in *spe-11* and *oops-1* mutants, require spatially and temporally regulated actin dynamics, with a highly controlled balance of assembly and disassembly (i.e., actin network dynamic instability). Mutations affecting the CYK-1 formin and PFN-1 profilin disrupt polar body formation likely in part by interfering with the assembly and function of the meiotic cytokinetic ring (Swan et al., 1998; Severson et al., 2002). Actomyosin contractility and cortical destabilization appear to drive membrane ingressions at the cortex during the meiotic divisions (Fabritius et al., 2011; Schlientz and Bowerman, 2020; Quiogue et al., 2023). It has been proposed that global actomyosin contractility at the cortex provides the force that pushes the spindle through the actomyosin-free center of the constricting and ingressing contractile ring (Fabritius et al., 2011). Meiotic fidelity also requires cooperation and crosstalk between the actin and microtubule cytoskeletons. A recent study in *C. elegans* presented a model in which a proper balance between cortical actomyosin-generated tension and microtubule-mediated stiffness is important for controlling membrane ingression during polar body extrusion (Quiogue et al., 2023). This model proposes that the assembly and the function of the contractile ring, and polar body formation, depend on an optimal balance between membrane ingression and cortical stiffness, generated by actin–microtubule crosstalk. Cytoplasmic microtubule scaffolding structures, called linear elements, which are closely associated with the cortex, promote cortical stiffness that limits actomyosin-dependent membrane ingressions. F-actin destabilization at the cortex appears to play a role in generating the membrane ingressions because semi-dominant temperature-sensitive mutations in *act-2*, which affect actin’s ATP-binding site, cause abnormal membrane ingressions and incompletely penetrant defects in meiosis (Willis et al., 2006). Mutations in *oops-1* and *spe-11* might disrupt actin–microtubule cross talk at yet another level. Specifically, we observed that approximately 50% of *oops-1* and *spe-11* mutants, as well as the double mutant, arrest in meiosis. One possibility is that these embryos arrest owing to activation of the spindle-assembly checkpoint, which is active during meiosis (Furuta et al., 1999; Kitagawa et al., 2002; Stein et al., 2007; Bohr et al., 2015). Future work will be important for evaluating whether and how the biochemical activities we ascribe to the OOPS-1–SPE-11 complex as an F-actin destabilizer underlie the penetrant defects in meiotic cytokinesis we describe here.

Several observations suggest that the activity of the OOPS-1–SPE-11 complex might be highly regulated. Ectopic expression of SPE-11 in the female germline results in co-localization with OOPS-1 to the cortex of germ cells and oocytes without producing detrimental consequences. In fact, the complex retains its function, which is only manifested after oocyte meiotic maturation and fertilization. If our *in vitro* biochemical experiments have predictive value, the expectation might be that ectopic formation of the complex would result in cortical destabilization in the germline, which is not observed. One possibility is that the activity of the complex is confined to the period following oocyte meiotic maturation and fertilization, potentially by the numerous protein kinases that are known to be activated during this developmental time window. Indeed, both OOPS-1 and SPE-11 are phosphoproteins with multiple modification sites. These sites localize to non-essential regions of the two proteins, which can be deleted without phenotypic consequences, consistent with the possibility that they might be regulatory or modulatory. Mutagenesis studies will be required to determine the consequences of individual and multiple phosphorylations. Potentially, some modifications could be inhibitory and others activating. It is notable that PP1 phosphatases GSP-3/4 co-purify in both OOPS-1 and SPE-11 TAP experiments, and genetic experiments are consistent with the idea that these PP1 phosphatases promote OOPS- 1–SPE-11 function. An attractive possibility is that regulation of the complex’s activity by phosphorylation could modulate actin dynamics.

A surprising finding, first shown by Browning and Strome (1996) and validated here at a quantitative level using the new tools that have become available in the interim, is that SPE-11 can mediate its function when provided via the oocyte. Why then has nature gone to the trouble of separating the complex into gametes of opposite sex? Current data provide few clues and fewer answers, so we can only speculate. It is notable that OOPS-1 is most abundantly expressed in the distal germline where it does not appear to have a required role—*oops-1* null mutants are maternal- effect lethal. Whether *oops-1* has a non-essential or redundant role will require additional analysis. If it does, perhaps SPE-11 might interfere with this function through its high-affinity interaction with OOPS-1? Alternatively, the separation of OOPS-1 and SPE-11 into gametes of opposite sex might relate to the fact that female meiosis in *C. elegans* is under multiple paternal controls: the MSP hormone promotes oocyte meiotic maturation (Miller et al., 2001); SPE-11 promotes meiotic progression and meiotic cytokinesis; and sperm factors, including the GSP-3/4 PP1 phosphatases and the GSKL-1/2 glycogen synthase kinase homologs, promote meiosis II after fertilization (Ataeian et al., 2016; Banerjee and Srayko, 2022). Perhaps this regulation serves to ensure fertilization and prevent parthenogenetic development, which is a derived reproductive strategy in many nematodes.

## MATERIALS AND METHODS

### Strains, genetic analysis, phenotypic analysis, and whole genome sequencing for mutant identification

The genotypes of strains used in this study are reported in File S5. Genes and mutations are described in WormBase (Sternberg *et al*., 2024) or in the indicated references. Culture and genetic manipulations were conducted at 20°C as described (Brenner, 1974), except for the analysis of *spe-11(tn2094 tn2145*ts) or its suppressors, which were conducted at 15°C or 25°C, and RNA interference (RNAi), which was conducted at 22°C. The balancer chromosomes (Dejima et al., 2018) used for *spe-11* and *oops-1* were *tmC18[dpy-5(tmIs1236)]* I and *tmC25[unc-5(tmIs1241)]* IV, respectively. RNAi was performed by feeding with double-stranded RNA (dsRNA)-expressing *E. coli* (Timmons and Fire, 1998) using the culture media described by Govindan *et al*. (2006). The RNAi clone (I-2C18) targeting *mat-1*, which encodes the anaphase-promoting complex CDC27/APC3 subunit (Davis *et al*., 2002), was obtained from Source BioScience (Nottingham, UK), and its identity was validated by Sanger sequencing. Exposure to *mat-1(RNAi)* was initiated during the fourth larval stage.

To isolate dominant mutations able to suppress the *spe-11(tn2094 tn2145*ts) mutation, L4-stage hermaphrodites were mutagenized with 50 mM ethyl methanesulphonate (EMS) at 15°C. Between one and two mutagenized hermaphrodites were cultured on 100- by 15-mm or 150- by 15-mm petri dishes with nematode growth medium (NGM, Brenner, 1974) or peptone-enriched NGM at 15°C, with bacterial strains OP50-1 or NA22 as the food source. The F1 progeny were transferred to 25°C three to five days later and suppressed strains were isolated by their ability to efficiently produce progeny over multiple generations at 25°C. These rare dominant suppressors were sought in the F1 generation by screening approximately ∼1.5x10^6^ EMS-mutagenized genomes. Suppressed strains were outcrossed seven times using *tmC18[dpy-5(tmIs1236)]/+* males for whole genome sequencing.

Genomic DNA was prepared for whole-genome sequencing using the QIAGEN (Valencia, CA) DNeasy Blood and Tissue Kit. Illumina (San Diego, CA) libraries of genomic DNA were prepared and sequenced by Azenta GENEWIZ (South Plainfield, NJ) to approximately 100x coverage. As controls, the N2 strain, the DG5649 parent strain, and the FX30168 *tmC18* strain were also sequenced to filter out differences between the reference genome and our laboratory versions, as well as variants introduced during backcrossing. Reads were trimmed using TrimGalore (0.6.0) and mapped to the BSgenome.Celegans.UCSC.ce11 (1.4.2) genome using BWA mem (0.7.17-r1188). Aligned reads were sorted with Samtools (1.16.1) and duplicates were identified using MarkDuplicates (Picard, 2.18.16). The GATK Haplotype caller (4.1.2.0) was used to identify sequence variations with respect to the reference ce11 genome. The VariantAnnotation package (1.50.0) within R version (4.4.0) was used to filter variants for homozygous, single nucleotide polymorphisms with read depth ≥ 15 that were not present in all samples. These variants were annotated using the TxDb.Celegans.UCSC.ce11.ensGene (3.15.0) annotation package to consider all splice donor/acceptor mutations and non-synonymous protein coding mutations.

The following mutations were found to exhibit linkage to chromosome I, as they were balanced by *tmC18*: *chs-1(tn2189, tn2191, tn2193, tn2195, tn2198, tn2201, tn2210, tn2242,* and *tn2248)* and *gsp-3(tn2202)*. These mutations have not been separated from *spe-11(tn2094 tn2145*ts). *egg- 3(tn2190* and *tn2205)* were found to exhibit linkage to chromosome II, as they were balanced by *mnC1[dpy-10(e128) unc-52(e444) umnIs32]*. The two *egg-3* alleles were separated from *spe- 11(tn2094 tn2145*ts) and found to be viable and fertile at 25°C. The suppressor mutations were isolated in the F1 generation, suggesting they were dominant. To explicitly test dominance for *chs-1(tn2189, tn2191, tn2198, tn2201,* and *tn2210)* and *gsp-3(tn2202)*, *spe-11(tn2094 tn2145*ts*)/tmC18; tnEx265[str-1::gfp]* males were crossed to the suppressed (sup) strains at 25°C [genotypes: *sup spe-11(tn2094 tn2145*ts)] and hermaphrodites of genotypes *sup/+ spe-11(tn2094 tn2145* ts*)]; tnEx265* were examined at 25°C and found to be fertile, indicating dominance. For *egg-3(tn2190* and *tn2205)*, adult hermaphrodites of genotype *egg-3(tn2190* or *tn2205)/mnC1[umnIs32]*; *spe-11(tn2094 tn2145*ts) were generated at 25°C and found to be fertile.

### Genome editing

Plasmids that express single-guide RNAs (sgRNAs) under the control of the U6 promoter were generated as described (Arribere *et al*. 2014). Repair templates used to tag *oops-1* and *spe-11* with *gfp* were also generated as described (Dickinson *et al*. 2015). Repair templates used to tag *spe-11* with mSCARLET-I were as described (Spike *et al*., 2022). Genome editing was performed by injecting wild-type adult gonads with a DNA mix containing a repair template (10 ng/μl), one or more sgRNA plasmids (25 ng/μl each), Cas9-expressing plasmid (pDD162, 50 ng/μl), and injection marker (pMyo2::tdTomato, 4 ng/μl) and selecting for repairs and self-excising cassette (SEC) excisions using standard methods (Dickinson *et al*. 2015). Each repair was balanced and the SEC removed from heterozygotes. Fluorescently tagged *oops-1 and spe-11* alleles were homozygous fertile after SEC excision. All edited loci were validated by sequencing the repair junctions using PCR products as templates. sgRNA plasmids and repair templates used to generate deletions in *oops-1* and *spe-11* are described in File S6. Deletions were constructed using the *dpy- 10* co-conversion method (Arribere *et al*. 2014). The injection mix contained pJA58 (7.5 ng/µl), AF-ZF-827 (500 nM), appropriate sgRNA plasmid (25 ng/µl), the appropriate repair template (500 nM), and pDD162 (50 ng/µl).

Single-copy insertions were generated by combining MosSCI (Frøkjær-Jensen et al., 2008, 2014) and CRISPR/Cas9 approaches. Promoters, fluorescently tagged *oops-1* or *spe-11,* and 3’UTR regions were amplified from genomic DNA by PCR using the oligonucleotides listed in File S6. Amplified fragments were inserted into pKL129 (kind gift from the CGC) by Gibson assembly using NEBuilder HiFi DNA Assembly Master Mix (New England Biolabs, Ipswich, MA) to generate the MosSCI plasmids, which also contained a rescuing copy of *unc-119* (File S6). To generate single-copy insertion strains, *ttTi5605* II; *unc-119(ed3)* III adults from EG6699 (Frøkjær-Jensen et al., 2012) were injected with an injection mix containing MosSCI plasmids (50 ng/ µl), a co-injection marker (pMyo2::tdTomato, 4 ng/μl), and two sgRNA plasmids pXW7.01 and pXW7.02 (50 ng/ µl each, kind gift from Ekaterina Voronina, University of Montana), which were used to excise the *ttTi5605 Mos1* transposon on chromosome II. Injected animals were cultured at 22 °C for approximately 7 days, after which *unc-119*-rescued F2 progeny not containing the pMyo2::tdTomato co-injection marker were individually isolated and their progeny screened for stable transmission. Their progeny were crossed with males heterozygous for the *mnC1[dpy- 10(e128) unc-52(e444) umnIs32]* balancer chromosome, and their progeny that do not segregate *unc-119(ed3)* were identified by PCR using a cleaved amplified polymorphic sequence (CAPS) marker (File S6). Correct targeting was verified by conducting PCR and by Sanger sequencing. Oligonucleotides used as PCR primers or for Sanger sequencing are listed in File S6.

Endogenous fluorescent reporter tag insertions for EGFP::CHS-1 were conducted as described (Vincencio et al., 2019). The injection mix contained Cas9 protein (250 ng/µL), universal tracrRNA (141.97 ng/µL), *dpy-10* crRNA (14.39 ng/µL), allele specific crRNA (59 ng/µL), *dpy- 10* repair oligonucleotide (28 ng/µL), and allele specific oligonucleotide (116 ng/µL). All reagents were obtained from Integrated DNA technologies, Inc. (IDT, Coralville, IA). Guide and repair templates used to generate insertions are described in File S6. Each injection mix was injected into the germline of young adult hermaphrodites and the F1 generation screened with a combination of single-worm propagation and lysis followed by detection of gene deletions or tag insertions by PCR. Each new edited strain was verified by Sanger sequencing. Oligonucleotides used as PCR primers or for Sanger sequencing are listed in File S6.

### Microscopy

#### Widefield microscopy

For the images in Fig. 1, FM4-64 and calcofluor were utilized to examine eggshell permeability and chitin formation, respectively. L4-stage hermaphrodites were placed onto fresh MYOB media and cultured for 24 h. Embryos were dissected in egg buffer [4 mM HEPES (pH 7.4), 94 mM NaCl, 3.2 mM KCl, 2.7 mM CaCl2, and 2.7 mM MgCl2] supplemented with 16 mM of FM4-64 (Bai et al., 2020) or 1:5 ratio of calcofluor:egg buffer respectively. Embryos were dissected in 5 µl of supplemented egg buffer on a cover slip using a scalpel. To prevent pressure on the embryos, a depression microscope slide was fixed to the cover slip using four droplets of Vaseline on the edges. Images were acquired on a Zeiss AxioObserver inverted widefield microscope using a 40x Plan-Neofluar (numerical aperture 1.3) objective lens and Axiocam 503 camera (Carl Zeiss Inc. Gottingen, Germany). Image processing and analysis were conducted using Zen Microscopy (Carl Zeiss Inc.) and ImageJ/Fiji software (Schindelin et al., 2012). All images were obtained using identical parameters, with brightness and contrast adjusted for better visualization.

For the images in Fig. 2 and the movies, live imaging of oocyte meiosis was conducted on a Zeiss AxioObserver inverted widefield microscope using a 100x Plan-Apochromat (numerical aperture 1.4) objective lens and an Axiocam 503 camera (Carl Zeiss Inc.). A microscope slide with a 2% agarose pad was prepared with 40 µl of 2 mM tetramisole. Adult hermaphrodites were placed and immobilized in the tetramisole and covered with a cover slip. For each sample, meiotic events were imaged in a single focal plane with a time lapse of 30 s intervals.

Microscope images in Fig. 4, panels A-D, Fig. 5, panels F and G, Fig. S2, and Fig S5 were acquired on a Nikon Ni-E microscope with either a Plan Apo λ 60x (numerical aperture 1.4) objective or a Plan Fluor 40x Oil (numerical aperture 1.3) objective using an ORCA FLASH sCMOS camera (Hamamatsu, Shizuoka, Japan) and NIS elements software (Nikon Inc., Melville, NY). Fig. S1C was acquired on a Zeiss (Thornwood, NY) motorized Axioplan 2 microscope with a 40x Plan-Neofluar (numerical aperture 1.3) objective lens using a AxioCam MRm camera and Zeiss AxioVision software.

#### Confocal microscopy

Localization and image analysis of Fig. 1, panels D-F, and Fig. 5, panels B-E, were conducted on an Andor Dragonfly spinning disc confocal microscope (Oxford Instruments, Abingdon, UK) using a Plan Apo 63x objective lens (numerical aperture 1.47) and a Zyla sCMOS camera (Oxford Instruments). A microscope slide with a 2% agarose pad was prepared with 40 µl of 20 mM tetramisole. Young adult worms were placed and immobilized in the tetramisole and covered with a cover slip. Germlines were imaged using Z-stack projections of a constant 0.5 µm or 0.2 µm per slice for Fig. 1 and Fig. 5, respectively. Image processing and analysis were conducted using Imaris image analysis software (Oxford Instruments) and ImageJ (Fiji). All images were obtained using identical parameters, with brightness and contrast adjusted for better visualization

Microscope images in Fig. 4, panels E and F, were acquired on a Nikon Ti2 inverted confocal microscope with a Plan ApoIR 60x objective (numerical aperture 1.27), motorized stage and Galvano scanner using a DUG hybrid four-channel detector system that combines gallium arsenide phosphide and multi-alkali photomultiplier tubes and NIS elements software (Nikon Inc.).

### Proteomics

Tandem affinity purification of OOPS-1 and SPE-11 were conducted using strains DG4800, DG5430, and DG5462 using modifications of a previously described protocol (Tsukamoto *et al*. 2017, 2020). The first immunopurification used a mixture of anti-GFP monoclonal antibodies 12A6 and 4C9 (Developmental Studies Hybridoma Bank, University of Iowa) and the second immunopurification used anti-FLAG monoclonal antibody M2 (Sigma-Aldrich, St. Louis, MO). Immunopurified proteins were precipitated with 16.7% trichloroacetic acid (TCA), washed with acetone at –20°C, and separated on a 12% NuPAGE Bis-Tris gel, stained with Colloidal Blue Staining Kit (Invitrogen). Lanes were subdivided into eight gel slices. The excised gel slices were subjected to in-gel trypsin proteolytic digestion as described previously (Thu *et al*., 2016) with the following change. During the alkylation step, 55 mM iodoacetamide was used instead of 55 mM methyl methanethiosulfonate. Post digestion, the peptides in each gel band were purified with a C18 Stage tip (Rappsilber *et al*., 2003). Eluates were vacuum dried. Mass spectrometry for Experiment I was performed at the Taplin Biological Mass Spectrometry Facility (Harvard Medical School) using an LTQ Orbitrap Velos Pro ion-trap mass spectrometer (Thermo Fisher Scientific, Inc., Waltham MA). Experiments II–VII were conducted at the Center for Metabolomics and Proteomics at the University of Minnesota. Experiments II and III utilized an Orbitrap Fusion liquid chromatography mass spectrometer (Thermo Fisher Scientific, Inc.). Experiments IV-VII used an Orbitrap Eclipse liquid chromatography mass spectrometer. The tandem mass spectrometry data was processed using Sequest (Eng *et al*., 1994). The *Caenorhabditis elegans* Universal Proteome UP000001940 protein sequence database was downloaded from Uniprot.org and merged with a common lab contaminant protein database (Frankenfield *et al*., 2022). We applied a 1% protein and peptide false discover rate using the Percolator algorithm (Käll *et al*., 2007). For phosphoprotein analysis, Experiments II-VII were reanalyzed using Proteome Discoverer 3.0 using phosphoserine, phosphothreonine, and phosphotyrosine in the database search parameters. Files S1 and S2 report additional technical details of the proteomic analyses, and the peptides identified in the OOPS-1 and SPE-11 immunopurifications, respectively. File S4 documents high-confidence phosphorylation sites identified in the OOPS-1 and SPE-11 immunopurifications based on multiple diagnostic b- and y- type fragment ions identified by manual inspection of the mass spectra.

### Expression and purification of the OOPS-1–SPE-11 complex

#### Expression plasmid construction

The expression plasmid for production of tagged versions of OOPS-1 and SPE-11 in *E. coli* was constructed as diagramed in Fig. S11, and the oligonucleotide primers used are listed in File S6*. oops-1* cDNA (TT767) and *spe-11* cDNA (TT768) codon optimized for *E. coli* were commercially synthesized using gBlock Hi-Fi (IDT, Coralville, IA) and were used as templates for PCR to amplify cDNA fragments encoding tagged version of OOPS-1 and SPE-11. Q5 DNA polymerase (New England Biolabs) and the primer pair of TT769 / TT770 and the pair of TT771 / TT772 were used to amplify *3xFLAG::TEV::oops-1a* and *S-tag::HA::PreScission::spe-11*, respectively. These PCR products were digested with appropriate restriction enzymes and were gel-purified using QIAprep 2.0 spin columns (Qiagen). Purified PCR fragments were ligated into the pRSFDuet-1 vector (Novagen, Madison, WI) between *Eco*RI and *Hin*dIII sites of the first multiple cloning site, and between *Nde*I and *Avr*II sites of the second multiple cloning site, respectively, (pTT197 and pTT204 plasmids) using T4 DNA ligase (New England Biolabs). *S-tag::HA::PreScission::spe-11* fragment was excised from pTT204 by *Nde*I and *Avr*II digestion and was inserted into pTT197 to generate pTT207, which contains both *6xHis::3xFLAG::TEV::oops-1* and *S- tag::HA::PreScission::spe-11*. To replace the 6xHis tag of OOPS-1 with a HaloTag, HaloTag cDNA fragment was amplified by PCR using the primer pair of TT909 and TT910 and was inserted between *Nco*I and *Hin*dIII digestion sites of pTT219 plasmid. The plasmids were transformed into DH5α cells and their DNA sequences were confirmed by Sanger sequencing.

#### Protein induction and purification

Expression of proteins was carried out in BL21-AI *E. coli* cells (Promega, Madison, WI). The transformed cells were cultured at 30 °C in LB medium containing 50 µg/ml of kanamycin until an OD600 of 0.4–0.6. OOPS-1 and SPE-11 proteins were induced at 30 °C for 4 hours by addition of 1 mM IPTG and 0.2% L-arabinose. Cells were harvested by centrifugation at 3,500 *x g* for 20 minutes at 4 °C, and proteins were extracted using Bacterial Protein Extraction Reagent (B-PER, Thermo Scientific) containing EDTA-free Halt protease inhibitors (Thermo Scientific), 5 units/ml of DNase I (Thermo Scientific) and 100 µg/ml of lysozyme (Thermo Scientific). The cell lysate was centrifuged at 20,000 *x g* for 15 minutes at 4 °C, and supernatant was used for protein purification.

For the primary purification, the supernatant was diluted with an equal volume of TBSN buffer (50 mM Tris-HCl, pH 7.4, 150 mM NaCl, 0.05 % NP-40) and applied ANTI-FLAG M2 affinity gel (Sigma-Aldrich) in batch for 3 hours at 4 °C. After the binding, the affinity gel was loaded onto a column and was washed with 20 column volumes of TBSN buffer. The bound proteins were eluted from the column by competitive elution with 3 column volumes of 150 µg/mL 3x FLAG peptide (Sigma-Aldrich) in TBSN buffer.

For the secondary purification, the eluate from ANTI-FLAG M2 affinity gel was diluted with an equal volume of HaloTag protein purification buffer (50 mM HEPES, pH 7.5, 150 mM NaCl, 0.05% NP-40), and applied HaloLink resin (Promega) in batch overnight at 4 °C plus an hour at room temperature. After the binding, the unbound proteins were removed by centrifugation at 2,500 *x g* for 5 minutes at 4 °C. The resin was washed with the 20 resin volumes of HaloTag protein purification buffer, and bound proteins were cleaved from the HaloLink resin by adding and mixing with the cleavage solution containing 283 units/ml of HaloTEV protease (Promega) in HaloTag protein purification buffer for 1.5 hours at room temperature. The cleaved proteins were collected by centrifugation at 3,200 *x g* for 5 minutes at 4 °C.

For tertiary purification, the proteins were incubated with Pierce Anti-HA Agarose (Thermo Scientific) for 3 hours at 4 °C in batch. After the binding, the agarose resin was loaded onto a column and was washed with 20 column volumes of TBSN buffer. The bound proteins were eluted from the column by competitive elution with 2 column volumes of 1 mg/mL HA peptide (Thermo Scientific) in TBSN buffer after incubation at 30 °C for 30 minutes. The eluted proteins from anti-HA agarose were dialyzed against KMEI buffer [50 mM KCl, 1 mM MgCl2, 1 mM EGTA, 10 mM imidazole (pH 7.0)] using Slide-A-Lyzer dialysis cassettes with 10 kDa molecular weight cut off (Thermo Scientific) overnight at 4 °C, flash-frozen on powered dry ice and stored at –80 °C. Eluted peptides at each step of purification were analyzed by running onto 4–12% Bis- Tris NuPAGE gel (Invitrogen, Waltham, MA) and visualized by staining colloidal blue staining kit (Invitrogen).

### Structure modeling and visualization methods

The amino acid sequences for OOPS-1, isoform 1a (Wormbase: CE43675, NCBI Accession: NP_500868) and SPE-11 (Wormbase: CE10744, NCBI Accession: P54217) were entered into the AlphaFold 3 Server (Abramson et al., 2024) to model a complex with one copy of each protein. Random seeds were used to initiate ten different model complexes. AlphaFold 3 generated five models in each run, and the results for the highest-ranked model from each run were analyzed. AlphaFold 3 calculated the confidence level for the position of each atom in the structure using the predicted local distance difference test (pLDDT). These values vary from 0 to 100, with values above 90 having high confidence and those below 50 indicating that the structure was likely incorrect. The pLDDT scores from the B-factor field of the mmCIF files were extracted using RStudio ver. 2024.12.0+467, and the pLDDT values for the alpha carbons (Ca) for each amino acid were plotted in a heatmap using geom_tile in ggplot2 (Wickham, 2016).

The models for the complexes containing the full-length proteins were visualized using ChimeraX Ver. 1.9 (Meng et al., 2023). The proteins were colored using the pLDDT values ranging from red (low values) to blue (high values). In each model, the N- and C-termini of each protein were mostly disordered, but the central part of the complex where the proteins were predicted to interact had a consistent structure in each model with reasonably high-confidence levels (pLDDT > 70). In addition, the predicted aligned error (PAE) in this part of the structure was low, suggesting that there was high confidence in the relative positions of the amino acids in the two proteins in this region. This again suggested that the proteins likely interact in this region. The structural models for the suppressor mutants were visualized using ChimeraX Ver. 1.9 as described above. The AlphaFold models for CHS-1 (https://alphafold.ebi.ac.uk/search/text/G5ECD6) and EGG-3 (https://alphafold.ebi.ac.uk/search/text/Q20402) were used. The AlphaFold 3 Server was used to model the structure of GSP-3 (WormBase:CE14754, NCBI accession O02658) with 2 Mn^2+^ ions because crystal structures of homologous mammalian serine/threonine PP1 phosphatases contained two metal ions in the active site (Egloff et al., 1995; Goldberg et al. 1995) that are required for enzyme activity (Zhang et al. 1996).

### Actin biochemistry

#### Purification of actin and formin

Actin was purified from an acetone powder of frozen chicken breast muscle (Trader Joe’s, Minneapolis, MN) by one cycle of polymerization and depolymerization (Spudich and Watt, 1971), followed by gel filtration on Sephacryl S-300 resin in G-Buffer (2 mM Tris-HCl (pH 8.0), 0.5 mM ATP, 0.5 mM DTT, and 0.1 mM CaCl2). Actin monomers were polymerized by dialyzing in 100 mM KCl, 2 mM MgCl2, 25 mM Tris-HCl (pH 7.5), and 0.3 mM ATP, and incubated overnight at 4 °C with a 1:10 molar ratio of actin to pyrenyl iodoacetamide (Product number P29, Thermo Fisher Scientific). Labeled F-actin was pelleted by ultracentrifugation at 120,000 x *g*, depolymerized, clarified, and gel-filtered in G-Buffer. We used extinction coefficients of 26,000 M^-1^cm^-1^ at λ = 290 nm for unlabeled actin and 22,000 M^-1^cm^-1^ at λ = 344 nm for pyrene, and the following relation to calculate the concentration of pyrene-labeled actin: [total actin] = (A290- (A344•0.127)) 26,000 M^-1^cm^-1^.

A construct encoding the FH1 and FH2 domains of the *S. cerevisiae* formin Bni1p (residues 1227-1776) was cloned into a pGEX-4T-3 plasmid (GE Healthcare Life Sciences, Pittsburgh, PA), which was modified to encode an N-terminal TEV protease recognition sequence and a C-terminal 6xHis tag. The protein was expressed overnight at 16 °C in a 1-liter culture of BL21(DE3) RP Codon Plus cells (Agilent Technologies, Santa Clara, CA). Resuspended cell pellets were lysed via sonication, clarified by centrifugation, and incubated with glutathione-Sepharose resin (Gold Biotechnology, Olivette, MO). The protein was eluted with 100 mM GSH (pH 8.0) in 50 mM Tris (pH 8.0), 100 mM NaCl, and 1 mM DTT and incubated with 2-5 µM MBP-tagged TEV protease overnight at 4 °C to remove the GST tag. The purified protein was separated from the TEV protease and cleaved GST by nickel affinity chromatography, concentrated using a 30,000 molecular- weight cutoff spin column (EMD Millipore, Burlington, MA), dialyzed into KMEI buffer (10 mM imidazole (pH 7.0), 50 mM KCl, 1 mM MgCl2, 1 mM EGTA) with 1 mM DTT, flash-frozen and stored at –80 °C. We used ProtParam (http://web.expasy.org/protparam) to calculate the extinction coefficient (Gasteiger et al., 2005).

#### Co-sedimentation assays

Ca^2+^-actin monomers were converted to Mg^2+^-actin via the addition of 0.05 mM MgCl2 and 0.2 mM EGTA. Samples containing 4 µM Mg^2+^-actin monomers were polymerized in KMEI buffer (10 mM imidazole (pH 7.0), 50 mM KCl, 1 mM MgCl2, 1 mM EGTA) for 1 hour at 22 °C in the absence or presence of 223 nM OOPS-1–SPE-11 complex. Polymerized samples were centrifuged for 30 min at 100,000 x *g*. Supernatants and pellets were separated and analyzed via Western blot. The supernatant fraction was first TCA precipitated as described above and resuspended in LDS sample buffer supplemented with a reducing agent (Invitrogen, Carlsbad, CA). Equivalent fractions of the supernatant and pellet fractions were separated using NuPAGE 4–12% Bis-Tris gels (Invitrogen, Carlsbad, CA) and visualized after western blotting. Blots were blocked with 5% nonfat dried milk. The primary antibodies used were mouse anti-HA.11 monoclonal 16B12 (Enzo Life Sciences, Farmingdale, NY) at a 1:30,000 dilution and mouse anti-actin monoclonal clone C4 (MP Biomedicals, Santa Ana, CA) at a 1:10,000 dilution. The secondary antibodies used were peroxidase-conjugated goat anti-mouse (Jackson ImmunoResearch Laboratories, West Grove, PA) at a 1:100,000 dilution. Detection was performed using SuperSignal West Femto Maximum Sensitivity Substrate and CL-XPosure film (Thermo Scientific).

#### Pyrene-actin assembly assays

Time courses of actin polymerization were collected by measuring fluorescence emission with a Molecular Devices SpectraMax Gemini EM fluorescence plate reader using Corning 96-well flat- bottom plates. Reactions containing 4 µM actin (20% pyrene-labeled) and a range of concentrations of OOPS-1–SPE-11 complex were polymerized in the absence or presence of formin in 10 mM imidazole (pH 7.0), 50 mM KCl, 1 mM MgCl2, 1 mM EGTA, 0.17 mM ATP, 0.5 mM DTT, 0.03 mM CaCl2, and 0.17 mM Tris-HCl (pH 8.0). Samples were excited at 365 nm and the fluorescence emission intensity was measured every 10 s at 407 nm over a period of 30 min. In reactions containing formin, the fluorescence signal was converted to polymer concentration by normalizing the fluorescence intensity to the final predicted actin polymer concentration, assuming a critical concentration of 0.17 µM. Reactions performed in the absence of formin did not attain equilibrium, thus precluding calculations of polymer concentration. The polymerization rate was calculated from the slope of the change in fluorescence signal at the point where half of the actin is polymerized.

## Data availability

All relevant data can be found within the article and its supplementary information.

## Author contributions

Conceptualization: T.T., A. J.-L., N.C., D.G.; Methodology, formal analysis, and investigation: T.T., J.K.K., M.E.Z., N.C., M.D.G., K.M.W., A.J.-L., D.G.; Writing, review, and editing, T.T., J.K.K., M.E.Z., N.C., M.D.G, K.M.W., A.J.-L., D.G.; Supervision, project administration and funding acquisition: N.C., A.J.-L., D.G.

## Funding Group

Award Group:

Funder(s): National Institutes of Health

Award Id(s): R35GM144029, R01GM122787, R35GM142524

## ACKNOWLEDGMENTS

This manuscript is dedicated to the memory of Andy Golden, our colleague and friend, whose encouragement, enthusiasm, and advice was pivotal to this study. We thank Shohei Mitani for sending many of the knockout alleles we screened in this study. Some strains and reagents were provided by the Caenorhabditis Genetics Center, which is funded by grant P40OD010440 from the NIH Office of Research Infrastructure Programs. We also thank WormBase for sequences and annotations. We also thank Margaret Titus for providing reagents for protein expression and purification, Ekaterina Voronina for sgRNAs for MosSCI, and Sara Olson for strains. Caroline Spike and Todd Starich provided many helpful suggestions.

## SUPPLEMENTAL INFORMATION

**Fig. S1.**
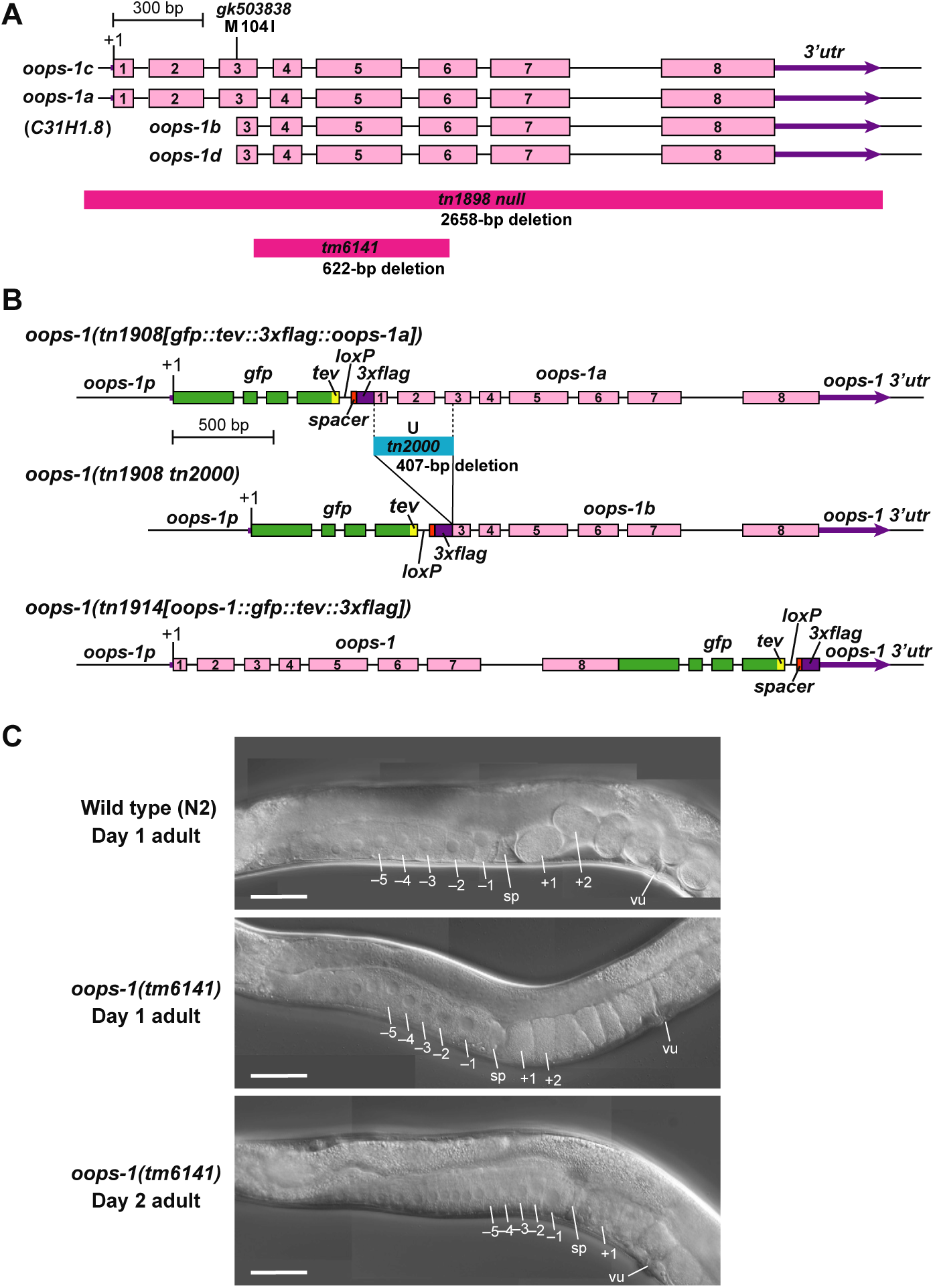
(Related to Fig. 1). (A) Genomic structure of *oops-1* showing several mutant alleles used in this study. (B) N- and C-terminal GFP-tagged alleles of *oops-1* generated by genome editing. The *oops-1(tn1908tn2000)* allele removes 407 bp, including exons 1 and 2 and a portion of exon 3, as well as introns 1 and 2, such that only *oops-1b/d* can be expressed from the locus. (C) DIC images of wild-type and *oops-1(tm6141)* adult hermaphrodites. Day-1 adult *oops-1(tm6141)* hermaphrodites contain 1-cell arrested embryos in the uterus [e.g., the region between the spermatheca (sp) and the vulva (vu)]. Day 2 adult *oops-1(tm6141)* hermaphrodites contain many arrested oocytes in the gonad arm and fewer embryos in the uterus owing to the depletion of sperm via polyspermy (stacked oocyte phenotype—see Table 1 in the main text). Proximal oocytes (–1 to –5) and newly fertilized embryos (+1 and +2) are indicated. Bars, 50 μm.

**Fig. S2.**
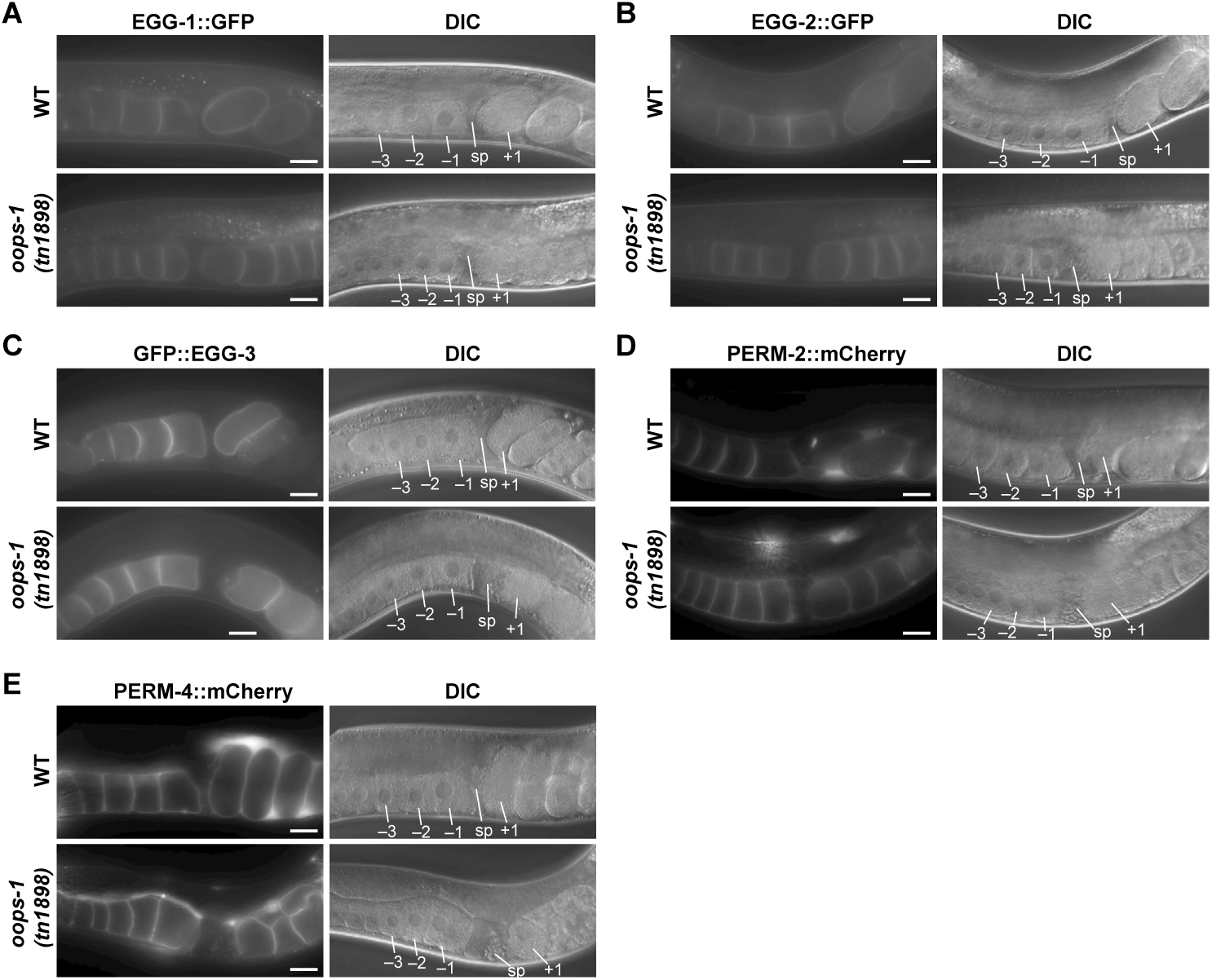
(Related to Fig. 1). ***oops-1* mutant oocytes and embryos exhibit normal localization of the EGG complex and PERM complex proteins investigated.** (A-E) Day 1 adult hermaphrodites were examined for the expression of EGG-1::GFP (A), EGG-2::GFP (B), GFP::EGG-3 (C), PERM-2::mCherry (D), and PERM-4::mCherry (E) in wild-type and *oops- 1(tn1898)* mutants. Proximal oocytes (–1 to –3), the spermatheca (sp), and newly fertilized embryos (+1) are indicated. Bars, 20µm.

**Fig. S3.**
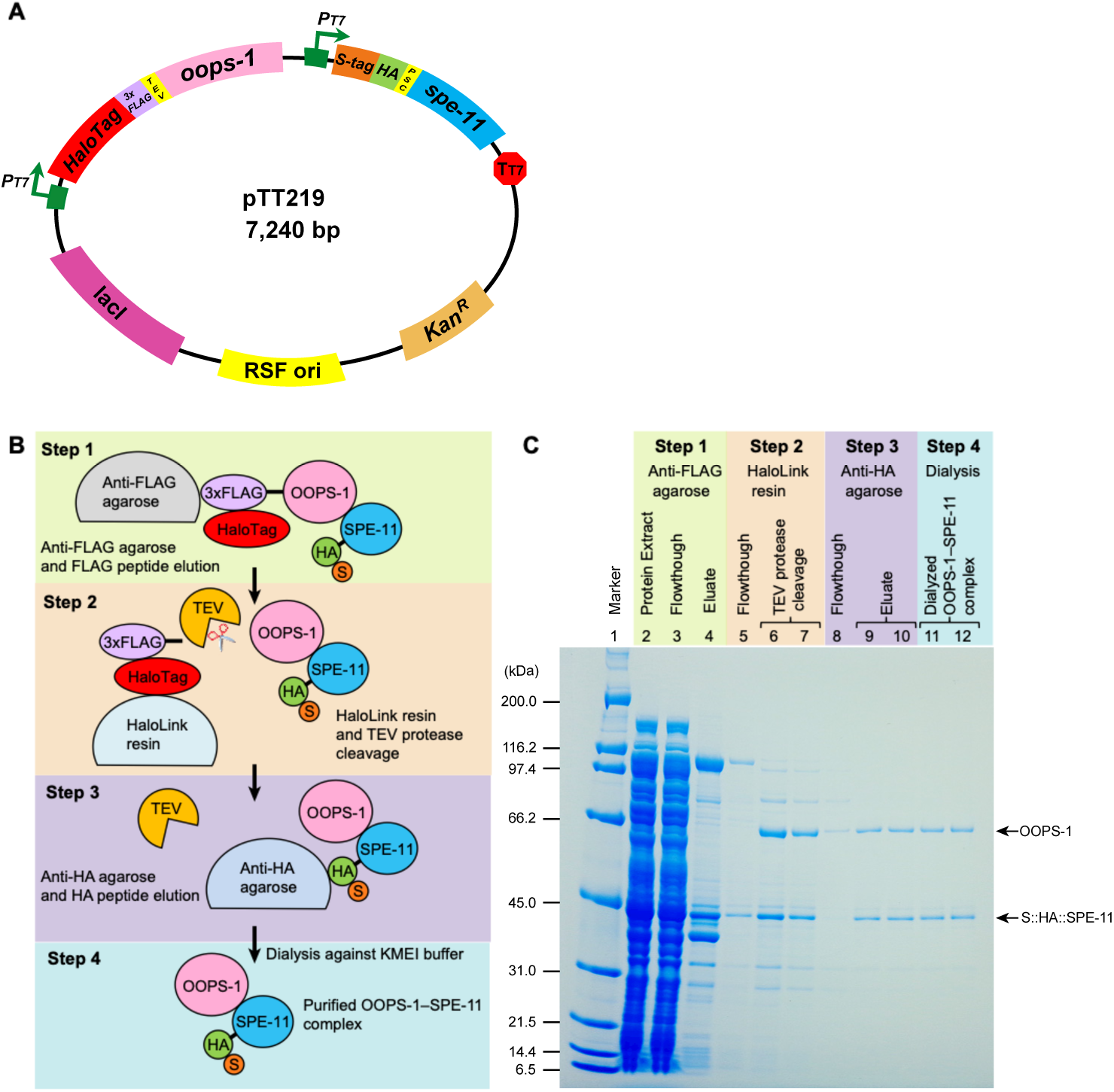
(Related to Fig. 3). Expression and purification of the OOPS-1–SPE-11 complex. (A) Plasmid for co-expression of tagged versions of OOPS-1 and SPE-11 in *E. coli*. (B) Flow chart of steps used for affinity purification of the complex. (C) Colloidal Coomassie-stained protein gel showing intermediate and final steps of the affinity purification of the complex.

**Fig. S4.**
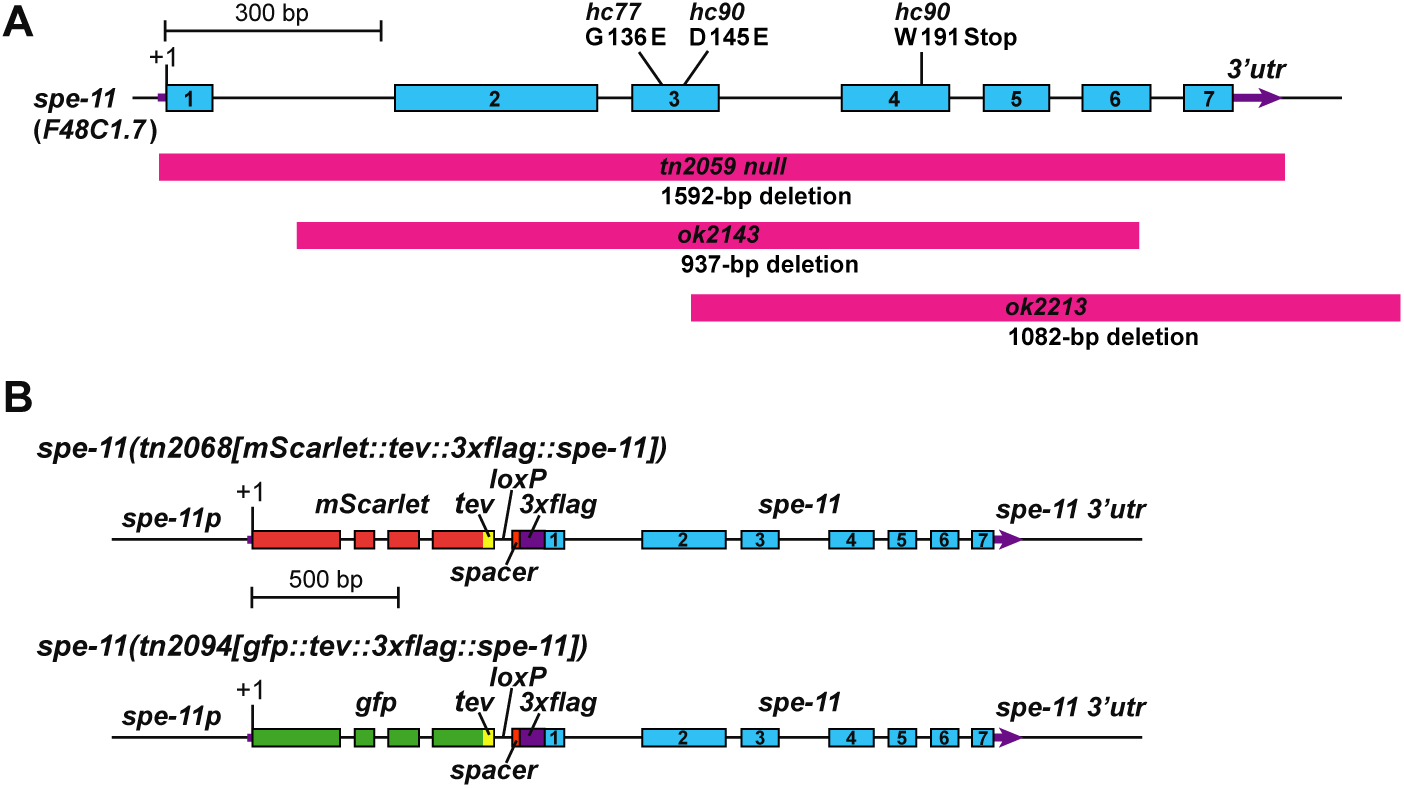
(Related to Figs 2 and 4). (A) Genomic structure of *spe-11* showing several mutant alleles used in this study. (B) Alleles of *spe-11* tagged with mScarlet or GFP generated by genome editing.

**Fig. S5.**
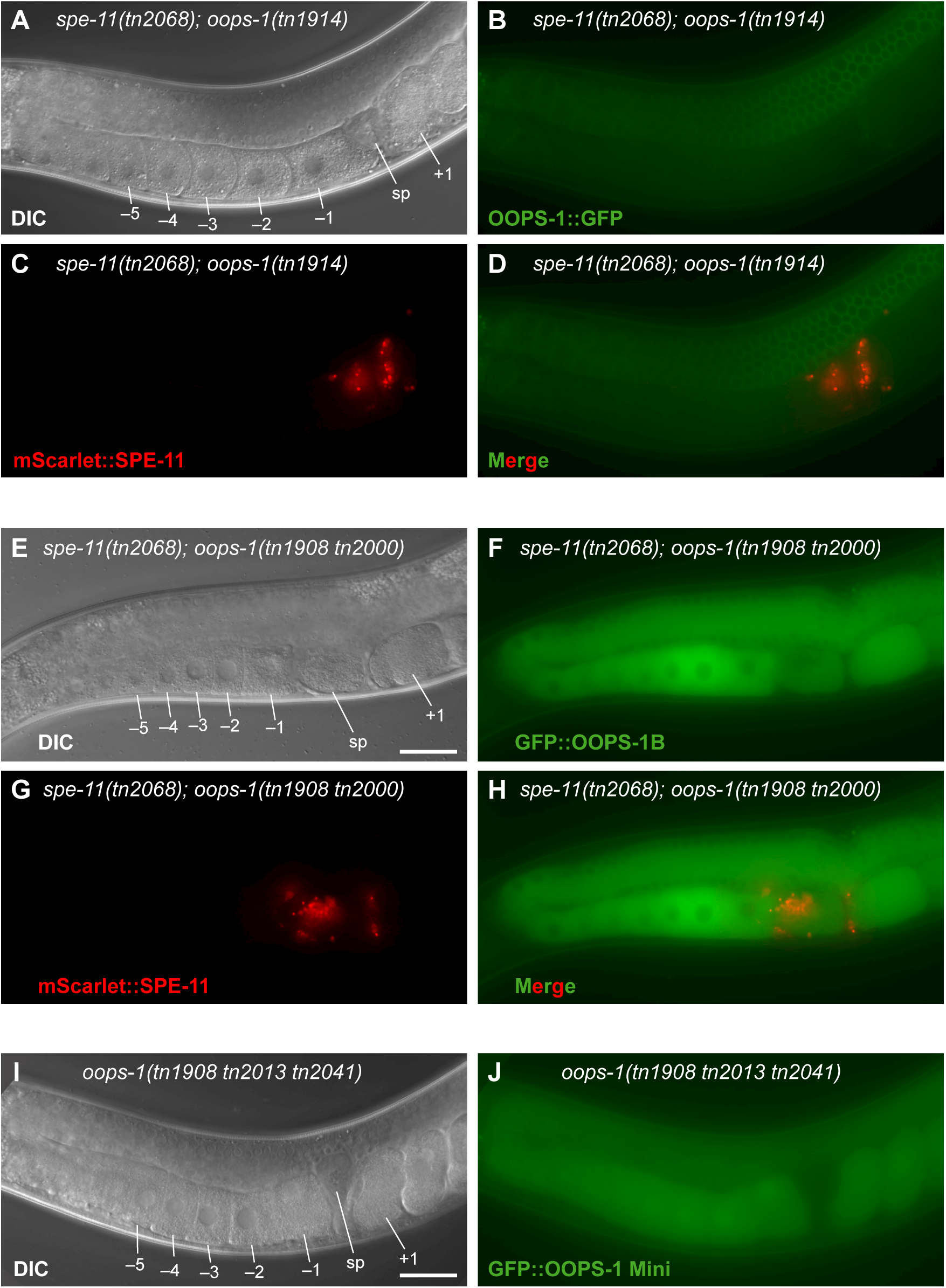
(Related to Fig. 4). DIC (A) and wide-field fluorescence micrographs (B-D) of a *spe- 11(tn2068[mScarlet::tev::3xflag::spe-11]); oops-1(tn1914[oops-1::gfp::tev::3xflag])* adult hermaphrodite (A-D). Tagging at the C-terminus with GFP should label all isoforms. DIC (E) and wide-field fluorescence micrographs (F-H) of a *spe-11(tn2068[mScarlet::tev::3xflag::spe-11]); oops-1(tn1908tn2000[gfp::tev::3xflag::oops-1b/d])* adult hermaphrodite. The deletion in *oops- 1(tn1908tn2000)* results in the expression of only OOPS-1B/D isoforms. (I, J) DIC (I) and a wide- field fluorescence micrographs (J) of GFP::OOPS-1(139-349) Mini [*oops- 1(tn1908tn2013tn2041)*]. Note that GFP::OOPS-1(139-349) Mini exhibits both cytoplasmic and nuclear localization. Proximal oocytes (–1 to –5), spermatheca (sp), and a newly fertilized embryo (+1) are indicated. Bars, 30 μm.

**Fig. S6.**
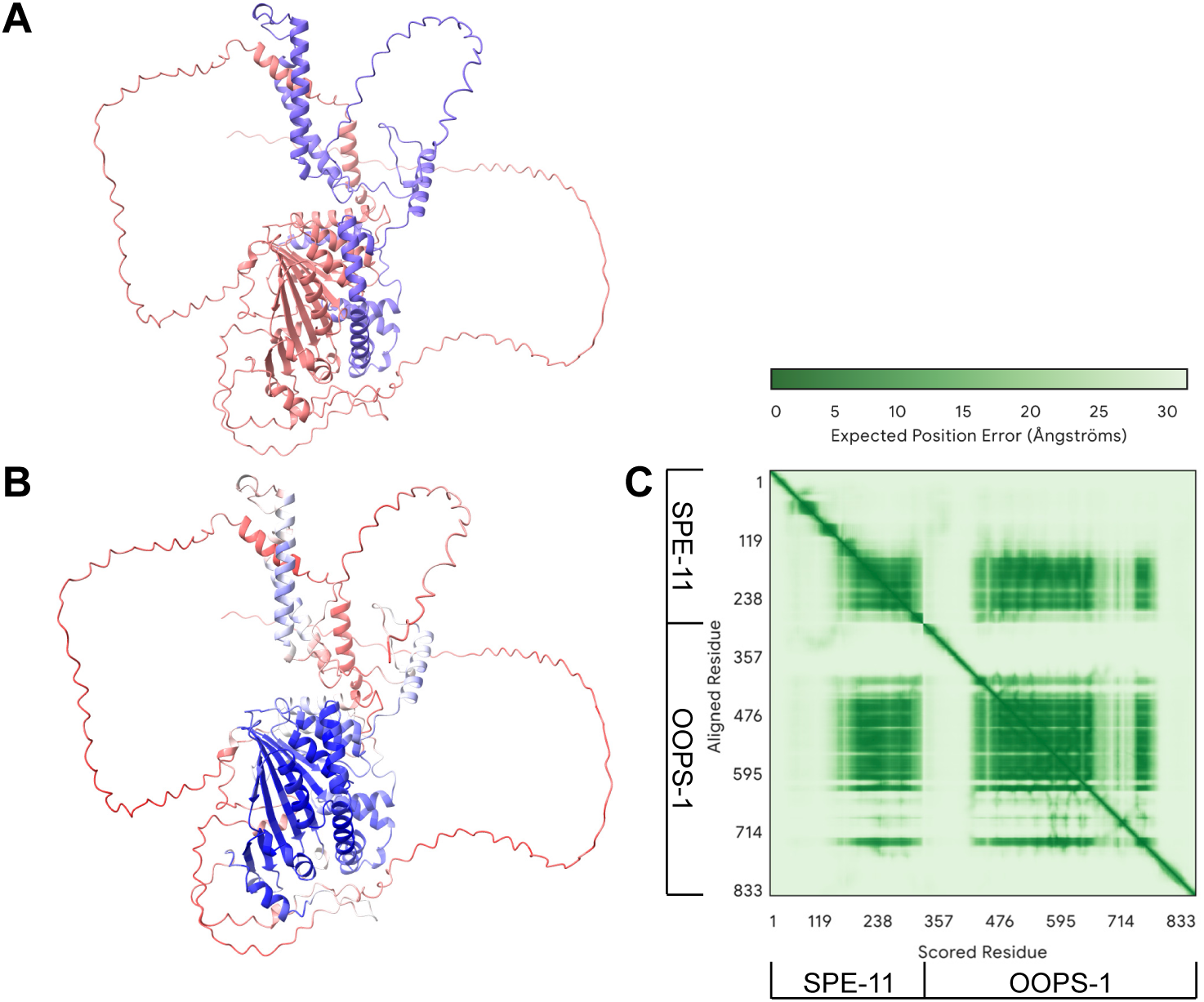
(Related to Fig. 6). Structures for the OOPS-1 and SPE-11 complex were modeled using the AlphaFold 3 server. The highest-ranked structure from model 1 is shown and colored based on (A) the protein with OOPS-1 in peach and SPE-11 in purple and (B) the pLDDT values, with red indicating low values and blue indicating high values. (C) Predicted aligned error (PAE) plot for a top-scoring model (model 1) of the OOPS-1–SPE-11 complex. PAE is a confidence measure for the relative position of any two residues within the predicted structure.

**Fig. S7.**
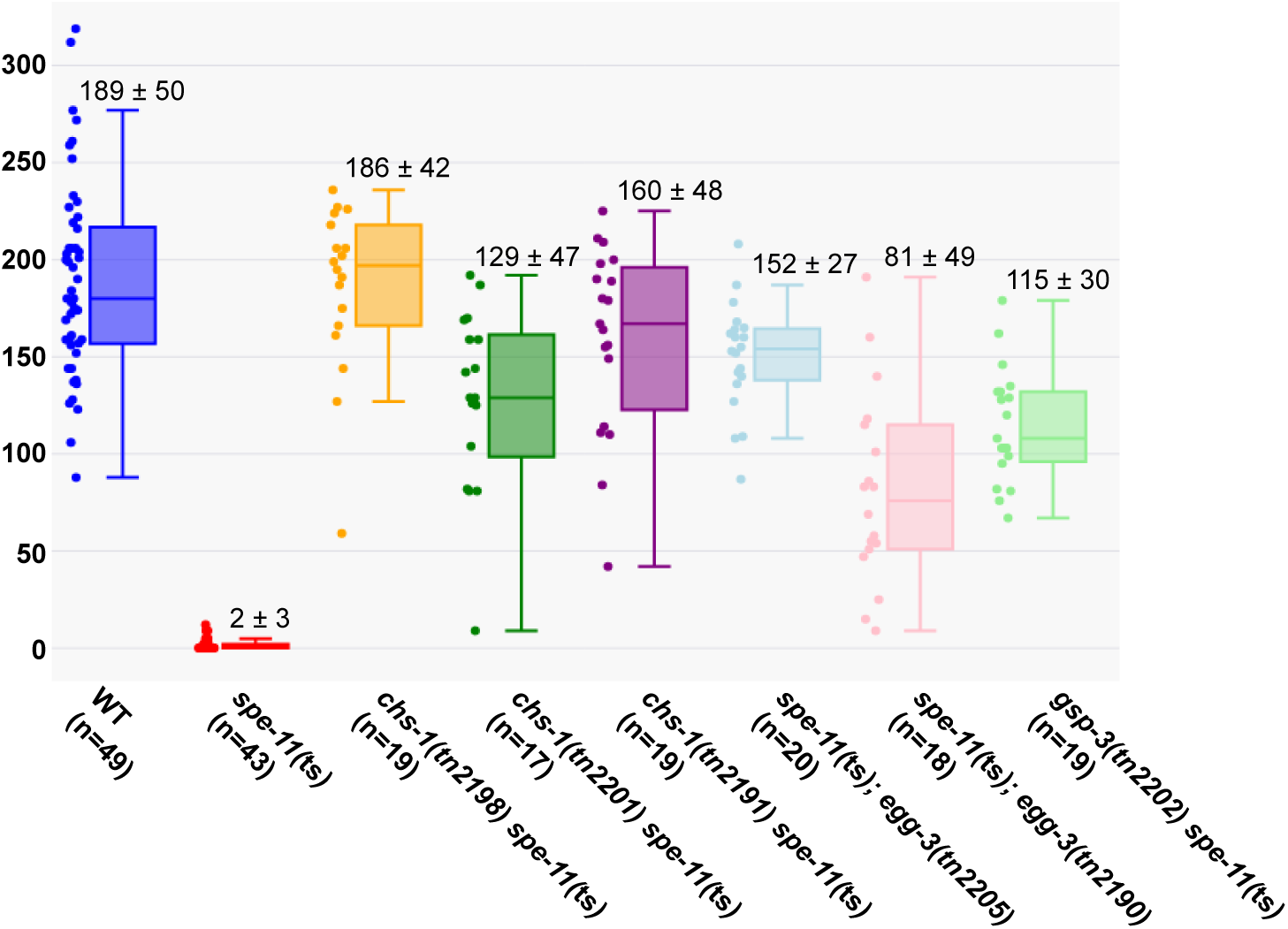
(Related to Table 5). Brood counts (total viable progeny) of the N2 wild type, DG5649 *spe-11*(ts), and suppressor strains of the genotypes listed (in order DG5830, DG5829, DG5837, DG5843, DG6032, and DG5842). Boxplots, showing means, quartiles calculated using the linear method, and all data points, were constructed using software from http://www.statskingdom.com.The average brood sizes for the suppressed strains were compared to that of DG5649 *spe-11*(ts) using a two-sample t-test with Welch’s correction, and in each case p<0.0001

**Fig. S8.**
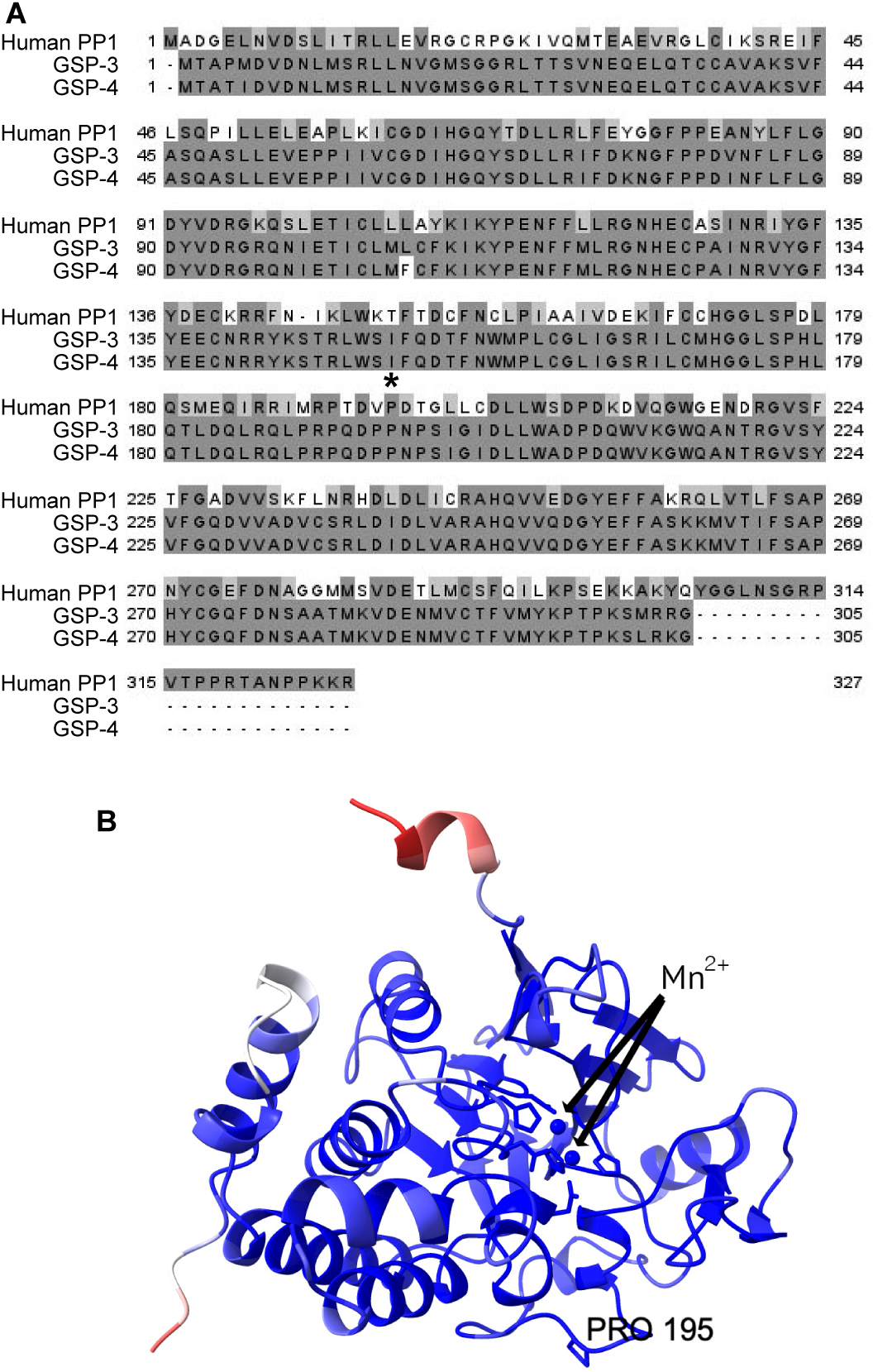
(Related to Table 5). (A) ClustalW alignment of human PP1B and GSP-3 and GSP-4 from *C. elegans*. The position of the P195S mutation in *gsp-3(tn2202)* is indicated by an asterisk. (B) AlphaFold predicted structure of GSP-3 showing the predicted location of P195 and the phosphatase active site and the metals in the phosphatase active site. The coloring is based on pLDDT values.

**Fig. S9.**
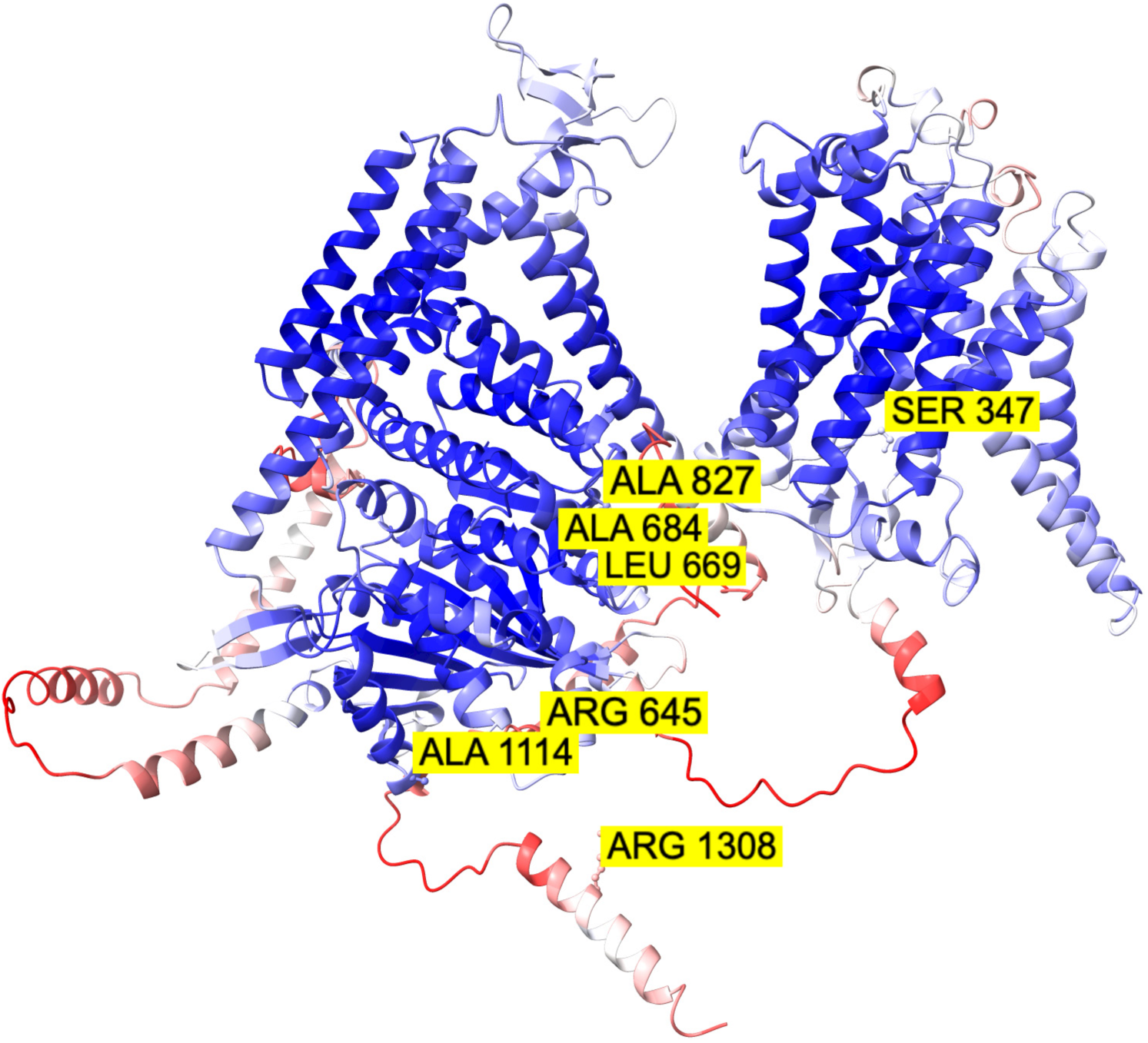
(Related to Table 5). AlphaFold predicted structure of CHS-1 showing the predicted location of the *spe-11(*ts) suppressor mutations. The coloring is based on pLDDT values.

**Fig. S10.**
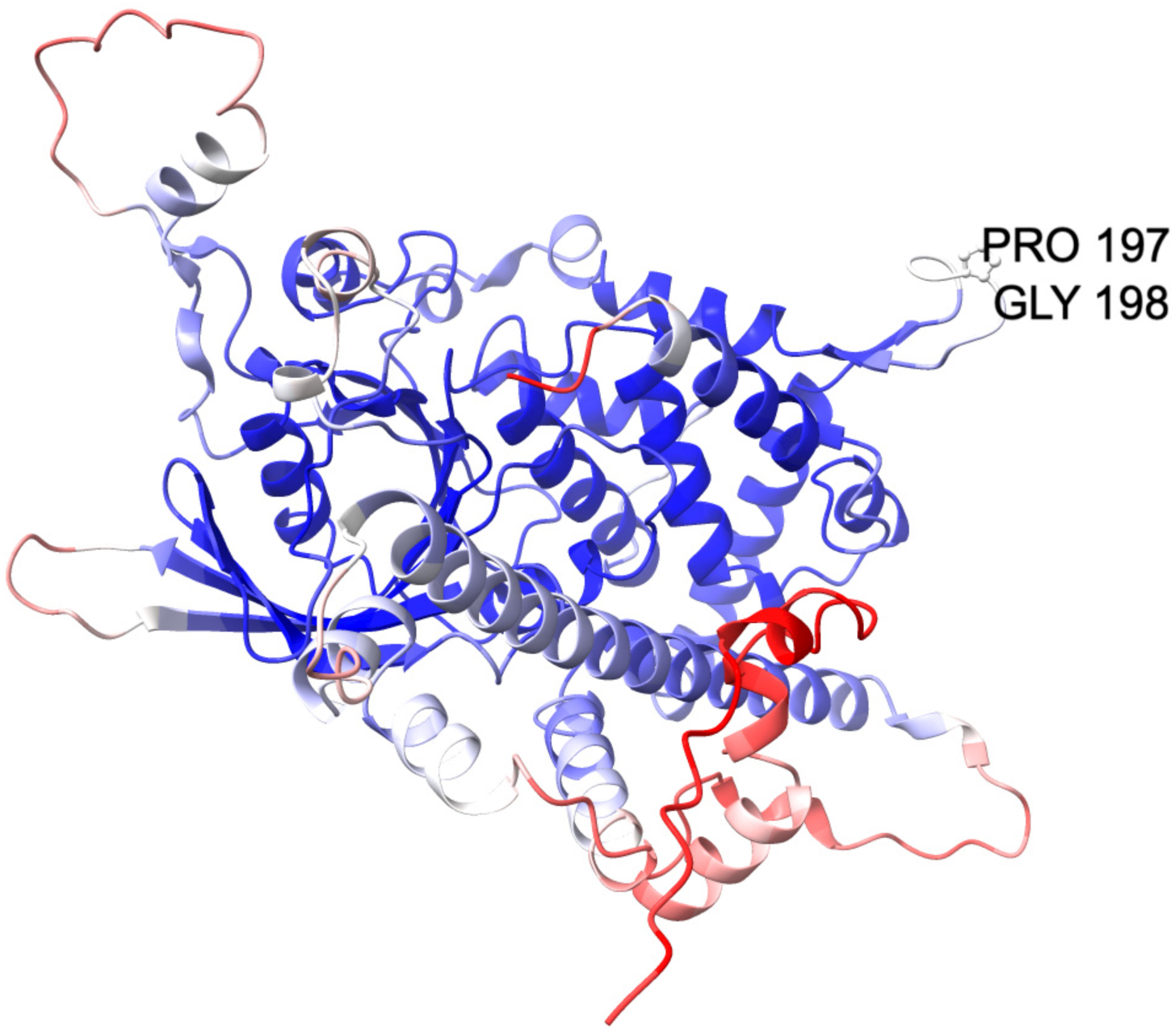
(Related to Table 5). AlphaFold predicted structure of EGG-3 showing the predicted location of the *spe-11(*ts) suppressor mutations. The coloring is based on pLDDT values.

**Fig. S11.**
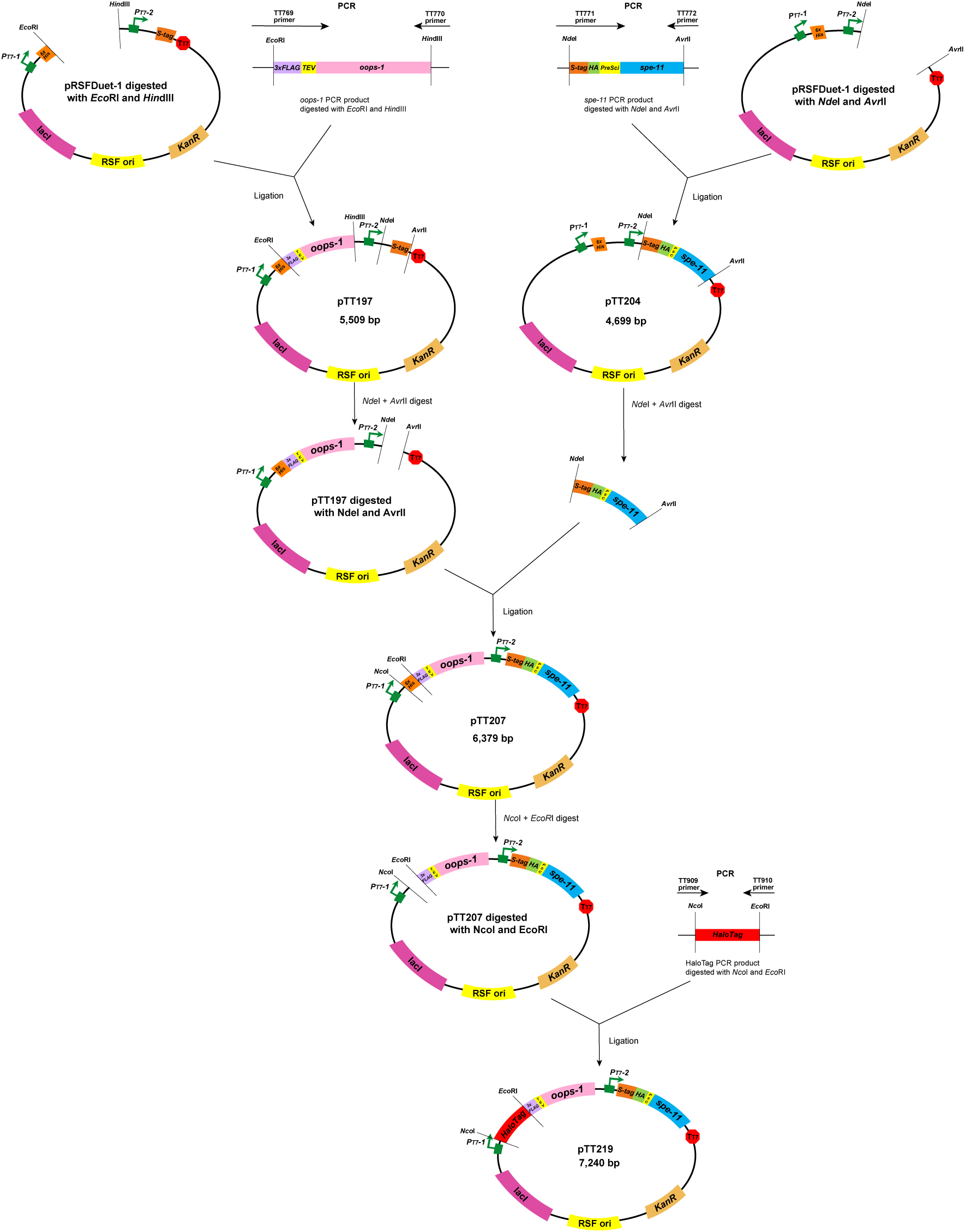
(Related to Fig. 3). Schematic of cloning methods to generate plasmids for expression of tagged versions of OOPS-1 and SPE-11 in *E. coli*.

## MOVIE LEGENDS

**Movie 1.** Time-lapse imaging of oocyte meiosis in the wild type. Chromatin is labelled in red, and microtubules are labelled in green. Time-lapse taken at 30 sec intervals for 49.5 min (100 frames).

**Movie 2.** Time-lapse imaging of oocyte meiosis in an *oops-1(tn1898)* null mutant. Segregation of chromosomes at meiosis I occurs but the first polar body fails to form, and the chromosomes collapse back into the oocyte. The meiosis II segregation event occurs with double the number of chromosomes, and the second polar body fails to form. This mutant phenotype is referred to the “completion phenotype” in the main text. Chromatin is labelled in red, and microtubules are labelled in green. Time-lapse taken at 30 sec intervals for 41.5 min (84 frames).

**Movie 3.** Time-lapse imaging of oocyte meiosis in an *oops-1(tn1898)* null mutant showing an example of the meiotic arrest phenotype. Segregation of chromosomes at meiosis I occurs but the first polar body fails to form, and the chromosomes collapse back into the oocyte. Chromosome segregation during meiosis II does not occur and the meiosis II spindle that encircles the chromosomes appears to fall apart. Chromatin is labelled in red, and microtubules are labelled in green. Time-lapse taken at 30 sec intervals for 64.5 min (130 frames).

**Movie 4.** Time-lapse imaging of oocyte meiosis in a *spe-11(hc90)* showing an example of the completion phenotype. Chromatin is labelled in red, and microtubules are labelled in green. Time-lapse taken at 30 sec intervals for 54.5 min (110 frames).

**Movie 5.** Time-lapse imaging of oocyte meiosis in a *spe-11(tn2059)* null mutant showing an example of the completion phenotype. Chromatin is labelled in red, and microtubules are labelled in green. Time-lapse taken at 30 sec intervals for 39.5 min (80 frames).

**Movie 6.** Time-lapse imaging of oocyte meiosis in *spe-11(tn2059)* null mutant showing an example of the meiotic arrest phenotype. Homologous chromosome segregation during meiosis I fails and the spindle fibers that encircle the chromosomes appear to fall apart. Chromatin is labelled in red, and microtubules are labelled in green. Time-lapse taken at 30 sec intervals for 47 min (95 frames).

## SUPPLEMENTAL FILES

**File S1.** Proteins identified in OOPS-1 TAP experiments.

**File S2.** Proteins identified in SPE-11 TAP experiments.

**File S3.** Results of whole-genome sequencing for the identification of *spe-11(*ts) suppressor mutations.

**File S4.** High-confidence phosphorylation sites identified in OOPS-1 and SPE-11 TAP data. Mass spectra showing phosphorylation identifications based on multiple diagnostic b- and y-type fragment ions.

**File S5.** *C. elegans* strains used in this study.

**File S6.** Sequence of oligonucleotides used in this study for generation of plasmids, genome editing, PCR, and sequencing.

