## Supplemental File 4 for "A sperm–oocyte protein partnership required for egg activation in *Caenorhabditis* elegans"

Tsukamoto, Kwah et al.
File S4
Mass spectral analysis identifying high-confidence sites of phosphorylation
Supported by multiple diagnostic b- and y-type fragment ions.
These are reported in Fig. 7 in the main text.

Several additional medium- and low- confidence sites of potential phosphorylation were also detected but are not reported in Fig. 7

### OOPS-1

Peptide tandem MS inspection for phosphorylation sites

OOcyte Partner of Spe-11 (Oops-1) OS=Caenorhabditis elegans OX=6239 GN=oops-1 PE=4 SV=3

☐ Annotate PTMs reported in Uniprot

☐ Show only PTMs

☐ Include PSMs that are Filtered Out

**Coverage: 54.87%**

**Found Modifications:**

**C** Carbamidomethyl (C)
**D** Deamidated (N,Q)
**G** Gln->pyro-Glu (N-term)
**O** Oxidation (M)
**P** Phospho (S,T)

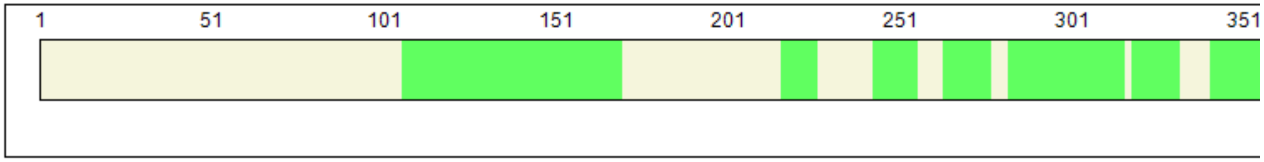


Sequence | Modification List

| Position | Target | Modification | Classification | Highest Peptide Confidence | Sequence Motif |
|---|---|---|---|---|---|
| 311 | N | Deamidated | Artefact | High | CCLKPCnSDAKTK |
| 318 | M | Oxidation | Artefact | High | SDAKTKmRQISVQ |
| 320 | Q | Gln->pyro-Glu | Artefact | High | AKTKMRqISVQVD |
| 324 | Q | Deamidated | Artefact | High | MRQISVqVDHVTG |
| 341 | Q | Gln->pyro-Glu | Artefact | High | SLPDVRqHEGLVC |
| 347 | C | Carbamidomethyl | Chemical derivative | High | QHEGLVcECLGDF |
| 349 | C | Carbamidomethyl | Chemical derivative | High | EGLVCEcLGDFSL |
| 361 | T | Phospho | Post-translational | High | LIRHRPtSVPVTK |
| 362 | S | Phospho | Post-translational | High | IRHRPTsVPVTKI |
| 439 | Q | Deamidated | Artefact | High | LEKAGIqELEDGD |
| 459 | S | Phospho | Post-translational | High | KYPKKRsIVDGAS |
| 465 | S | Phospho | Post-translational | High | SIVDGAsSETCED |
| 466 | S | Phospho | Post-translational | High | IVDGASsETCEDT |
| 468 | T | Phospho | Post-translational | High | DGASSEtCEDTQS |
| 469 | C | Carbamidomethyl | Chemical derivative | High | GASSETcEDTQSV |
| 473 | Q | Deamidated | Artefact | High | ETCEDTqSVGSSE |
| 474 | S | Phospho | Post-translational | High | TCEDTQsVGSSES |
| 477 | S | Phospho | Post-translational | High | DTQSVGsSESSEP |
| 478 | S | Phospho | Post-translational | High | TQSVGSsESSEPR |
| 480 | S | Phospho | Post-translational | High | SVGSSEsSEPRSP |
| 481 | S | Phospho | Post-translational | High | VGSSESsEPRSPE |
| 510 | S | Phospho | Post-translational | High | SDSESSsEDEKEN |
| 516 | N | Deamidated | Artefact | High | SEDEKEnMKSSIK |
| 517 | M | Oxidation | Artefact | High | EDEKENmKSSIKE |

### Experiment VII (lab code 18902)
### RLsSGDLNVIR (S108)
### <u>High confidence level</u>

OOPS-1

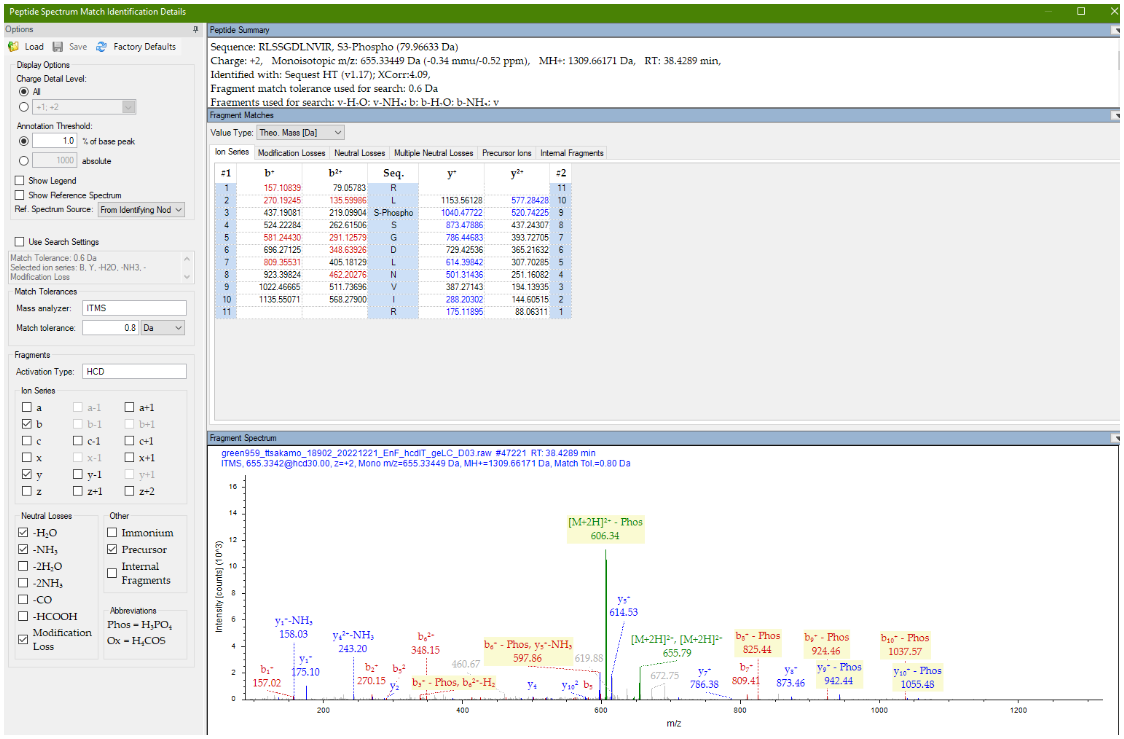


Diagnostic fragment ions detected

### Experiment V (lab code 18850)
### LSsGDLNVIR (S109)
### <u>High confidence level</u>

OOPS-1

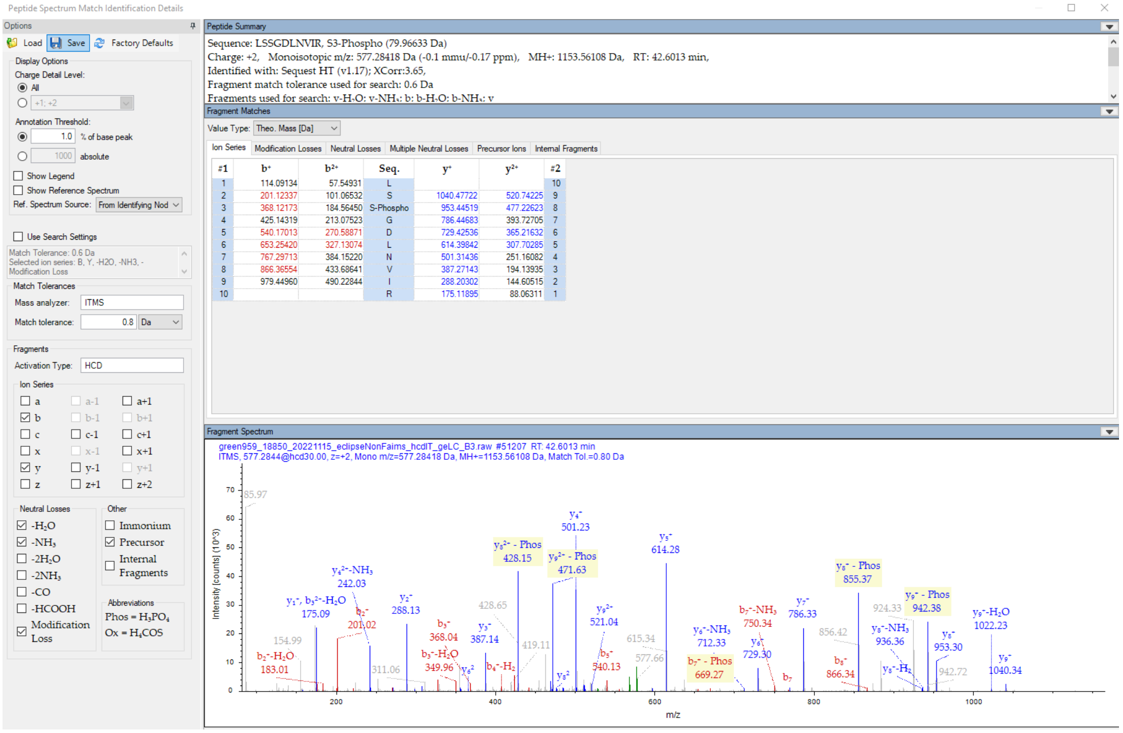


Diagnostic fragment ions detected

### Experiment VII (lab code 18902)
### RLssGDLNVIR (S108, S109)
### <u>High confidence level</u>

### OOPS-1

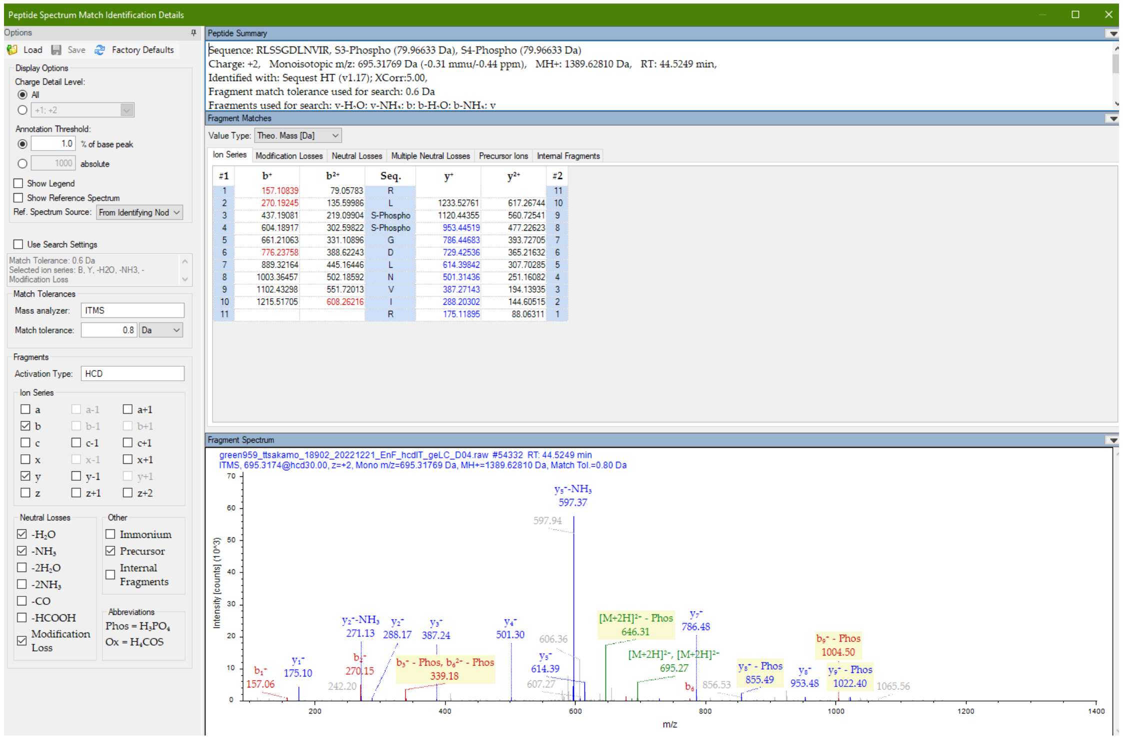


| #1 | $b^+$ | $b^{2+}$ | Seq. | $y^+$ | $y^{2+}$ | #2 |
|---|---|---|---|---|---|---|
| 1 | 157.10839 | 79.05783 | R | | | 11 |
| 2 | 270.19245 | 135.59986 | L | 1233.52761 | 617.26744 | 10 |
| 3 | 437.19081 | 219.09904 | S-Phospho | 1120.44355 | 560.72541 | 9 |
| 4 | 604.18917 | 302.59822 | S-Phospho | 953.44519 | 477.22623 | 8 |
| 5 | 661.21063 | 331.10896 | G | 786.44683 | 393.72705 | 7 |
| 6 | 776.23758 | 388.62243 | D | 729.42536 | 365.21632 | 6 |
| 7 | 889.32164 | 445.16446 | L | 614.39842 | 307.70285 | 5 |
| 8 | 1003.36457 | 502.18592 | N | 501.31436 | 251.16082 | 4 |
| 9 | 1102.43298 | 551.72013 | V | 387.27143 | 194.13935 | 3 |
| 10 | 1215.51705 | 608.26216 | I | 288.20302 | 144.60515 | 2 |
| 11 | | | R | 175.11895 | 88.06311 | 1 |

Diagnostic fragment ions detected

### Experiment VI (lab code 18879)
### RMtEEIDFDLK (T119)
### <u>High confidence level</u>

OOPS-1

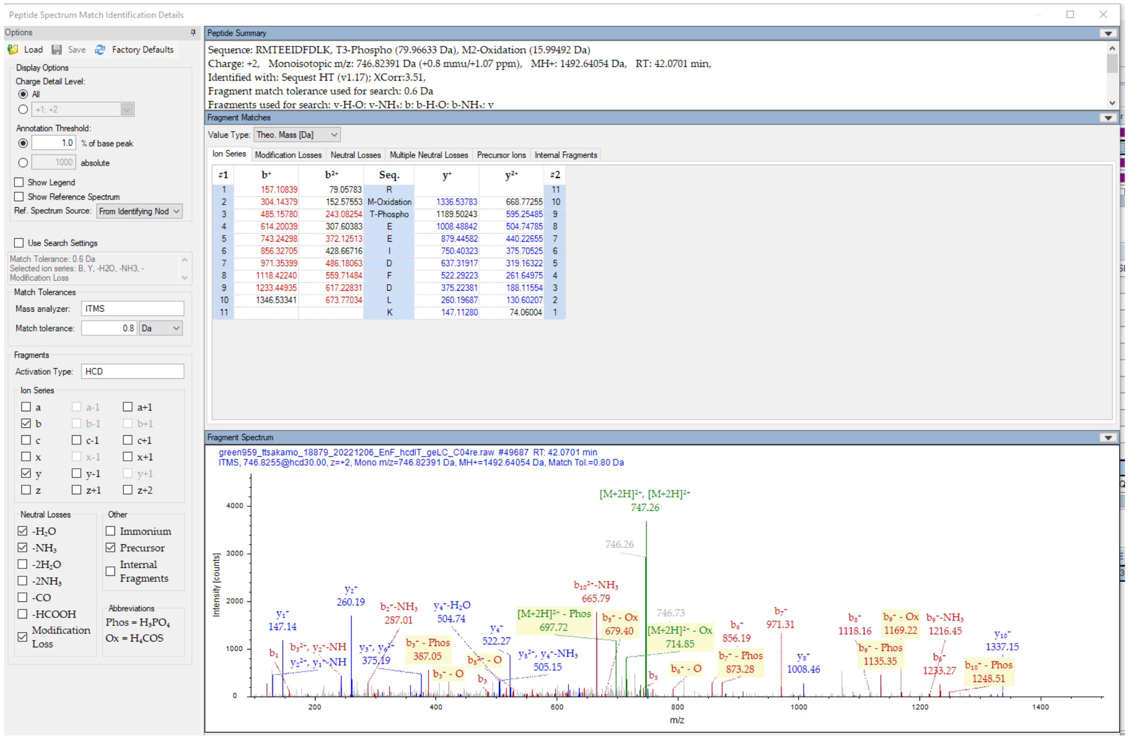


Diagnostic fragment ions detected

### Experiment V (lab code 18850)
### HRPtSVPVTK (T361)
### High confidence level

OOPS-1

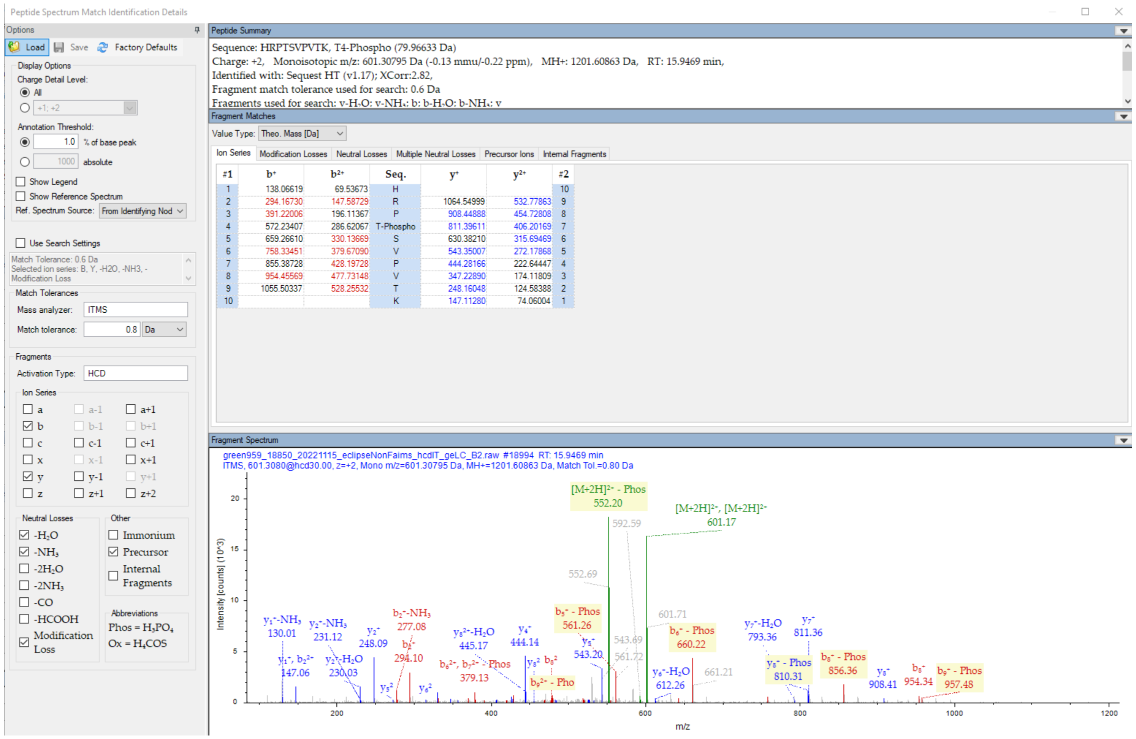


Diagnostic fragment ions detected

### Experiment V (lab code 18850)
### HRPTsVPVTK (S362)
### <u>High confidence level</u>

### OOPS-1

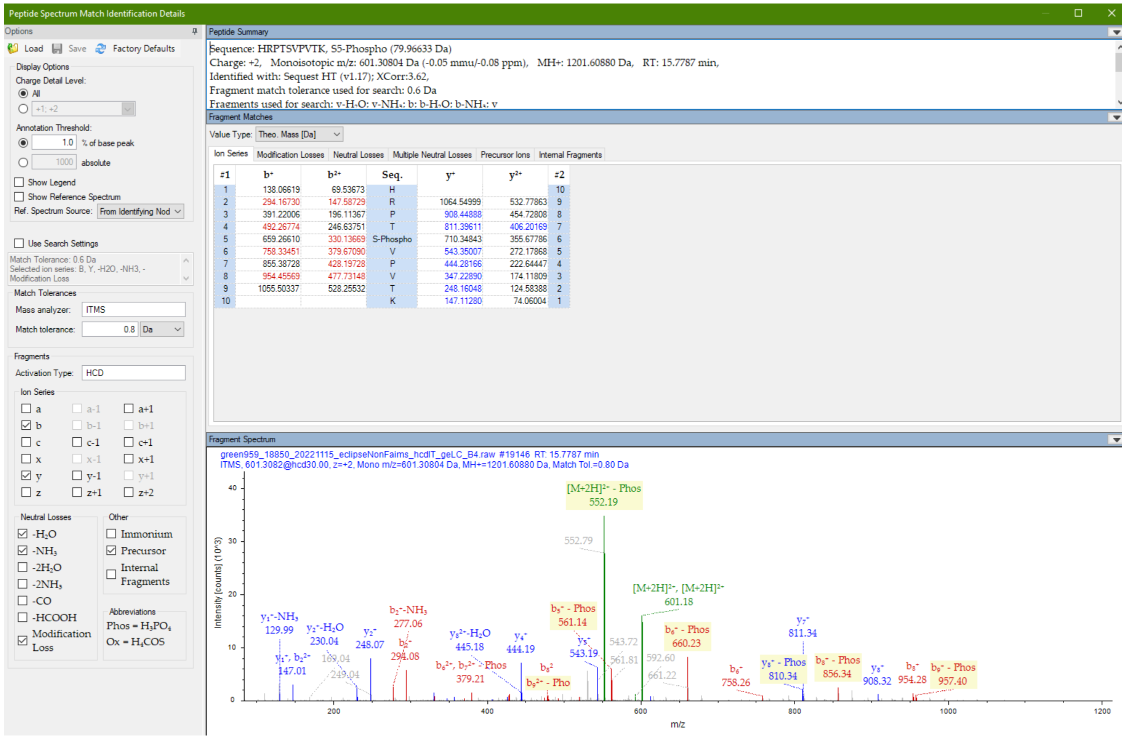


| #1 | $b^+$ | $b^{2+}$ | Seq. | $y^+$ | $y^{2+}$ | #2 |
|---|---|---|---|---|---|---|
| 1 | 138.06619 | 69.53673 | H | | | 10 |
| 2 | 294.16730 | 147.58729 | R | 1064.54999 | 532.77863 | 9 |
| 3 | 391.22006 | 196.11367 | P | 908.44888 | 454.72808 | 8 |
| 4 | 492.26774 | 246.63751 | T | 811.39611 | 406.20169 | 7 |
| 5 | 659.26610 | 330.13669 | S-Phospho | 710.34843 | 355.67786 | 6 |
| 6 | 758.33451 | 379.67090 | V | 543.35007 | 272.17868 | 5 |
| 7 | 855.38728 | 428.19728 | P | 444.28166 | 222.64447 | 4 |
| 8 | 954.45569 | 477.73148 | V | 347.22890 | 174.11809 | 3 |
| 9 | 1055.50337 | 528.25532 | T | 248.16048 | 124.58388 | 2 |
| 10 | | | K | 147.11280 | 74.06004 | 1 |

Diagnostic fragment ions detected with low intensity

### Experiment V (lab code 18850)

SIVDGASSETcEDTQSVGsSESSEPR (S477)

<u>High confidence level</u>

OOPS-1

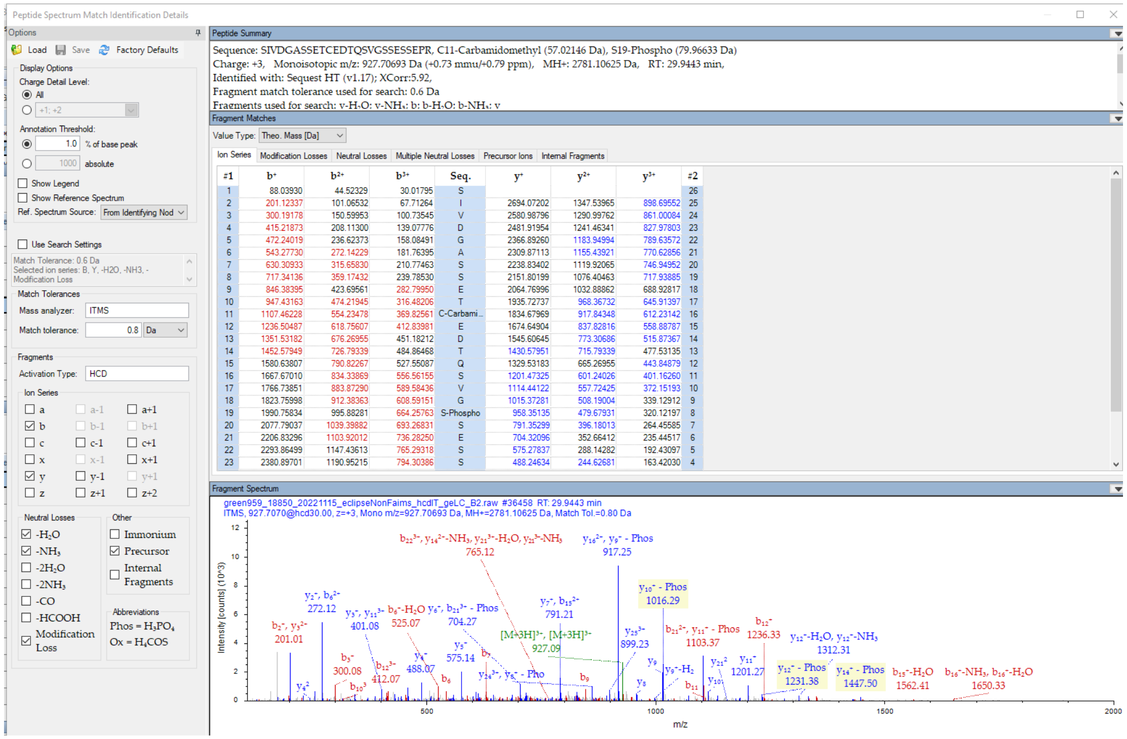


Diagnostic fragment ions detected

### Experiment V (lab code 18850)
### SIVDGASSETcEDTQSVGSSEsSEPR (S480)
### High confidence level

OOPS-1

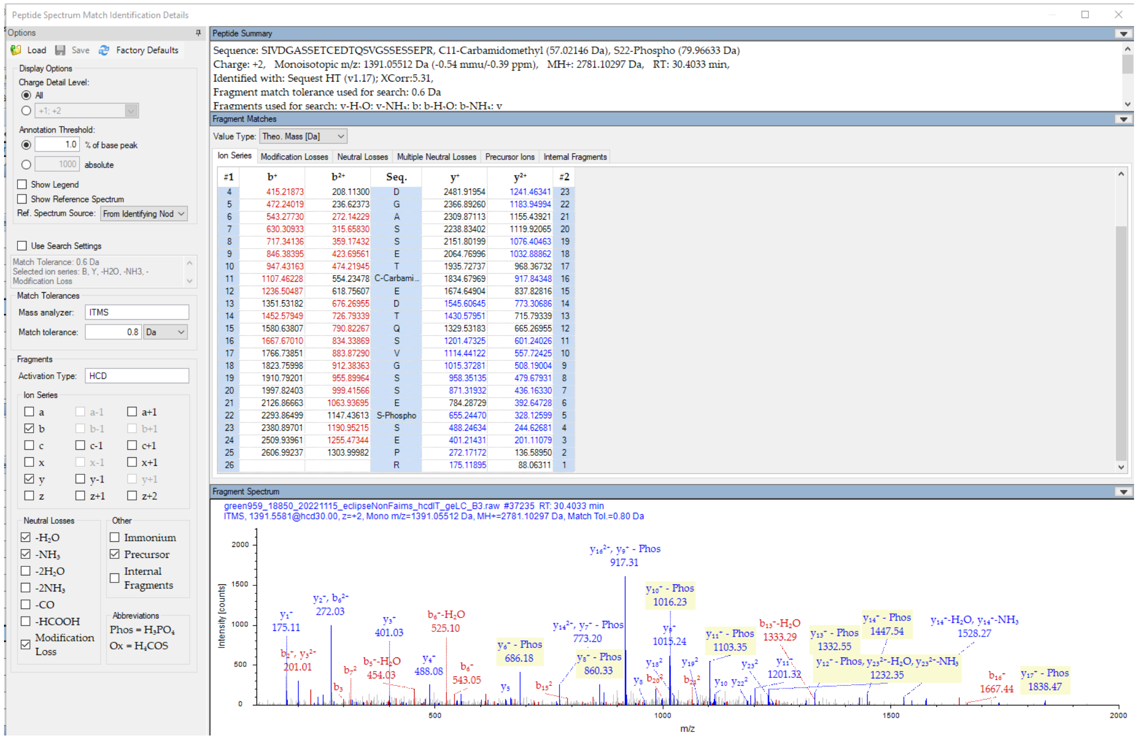


Diagnostic fragment ions detected

Experiment VI (lab code 18879)

SIVDGASSETcEDTQSVGSSESsEPR (S481)

<u>High confidence level</u>

OOPS-1

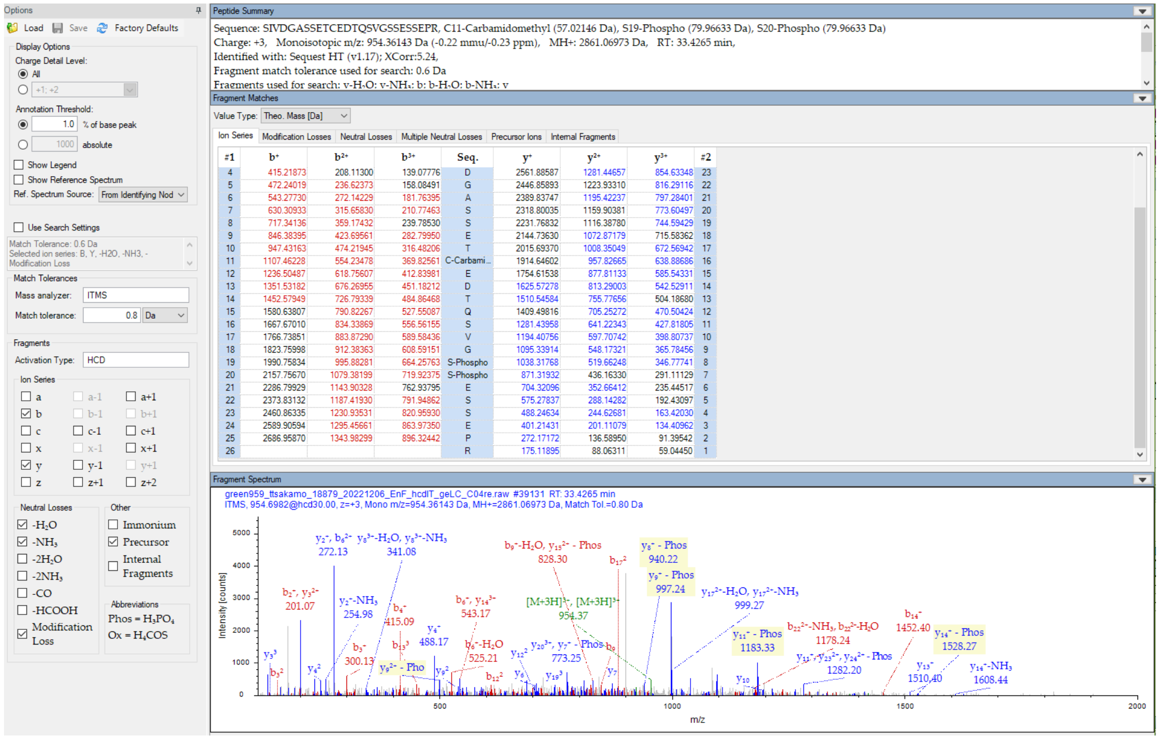


Diagnostic fragment ions detected

### Experiment VI (lab code 18879)

### SIVDGASSETcEDTQSVGSsEsSEPR (S478, S480)

### <u>High confidence level</u>

#### OOPS-1

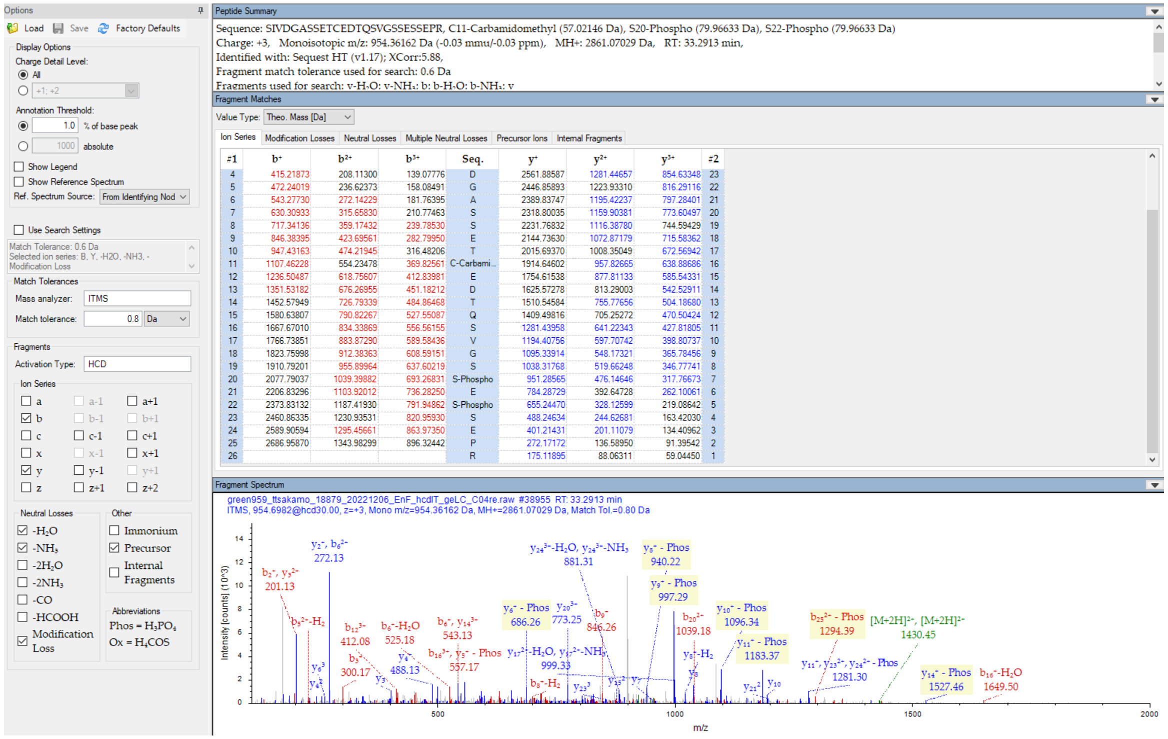


| #1 | b+ | b2+ | b3+ | Seq. | y+ | y2+ | y3+ | #2 |
|---|---|---|---|---|---|---|---|---|
| 4 | 415.21873 | 208.11300 | 139.07776 | D | 2561.88587 | 1281.44657 | 854.63348 | 23 |
| 5 | 472.24019 | 236.62373 | 158.08491 | G | 2446.85893 | 1223.93310 | 816.29116 | 22 |
| 6 | 543.27730 | 272.14229 | 181.76395 | A | 2389.83747 | 1195.42237 | 797.28401 | 21 |
| 7 | 630.30933 | 315.65830 | 210.77463 | S | 2318.80035 | 1159.90381 | 773.60497 | 20 |
| 8 | 717.34136 | 359.17432 | 239.78530 | S | 2231.76832 | 1116.38780 | 744.59429 | 19 |
| 9 | 846.38395 | 423.69561 | 282.79950 | E | 2144.73630 | 1072.87179 | 715.58362 | 18 |
| 10 | 947.43163 | 474.21945 | 316.48206 | T | 2015.69370 | 1008.35049 | 672.56942 | 17 |
| 11 | 1107.46228 | 554.23478 | 369.82561 | C-Carbami... | 1914.64602 | 957.82665 | 638.88686 | 16 |
| 12 | 1236.50487 | 618.75607 | 412.83981 | E | 1754.61538 | 877.81133 | 585.54331 | 15 |
| 13 | 1351.53182 | 676.26955 | 451.18212 | D | 1625.57278 | 813.29003 | 542.52911 | 14 |
| 14 | 1452.57949 | 726.79339 | 484.86468 | T | 1510.54584 | 755.77656 | 504.18680 | 13 |
| 15 | 1580.63807 | 790.82267 | 527.55087 | Q | 1409.49816 | 705.25272 | 470.50424 | 12 |
| 16 | 1667.67010 | 834.33869 | 556.56155 | S | 1281.43958 | 641.22343 | 427.81805 | 11 |
| 17 | 1766.73851 | 883.87290 | 589.58436 | V | 1194.40756 | 597.70742 | 398.80737 | 10 |
| 18 | 1823.75998 | 912.38363 | 608.59151 | G | 1095.33914 | 548.17321 | 365.78456 | 9 |
| 19 | 1910.79201 | 955.89964 | 637.60219 | S | 1038.31768 | 519.66248 | 346.77741 | 8 |
| 20 | 2077.79037 | 1039.39882 | 693.26831 | S-Phospho | 951.28565 | 476.14646 | 317.76673 | 7 |
| 21 | 2206.83296 | 1103.92012 | 736.28250 | E | 784.28729 | 392.64728 | 262.10061 | 6 |
| 22 | 2373.83132 | 1187.41930 | 791.94862 | S-Phospho | 655.24470 | 328.12599 | 219.08642 | 5 |
| 23 | 2460.86335 | 1230.93531 | 820.95930 | S | 488.24634 | 244.62681 | 163.42030 | 4 |
| 24 | 2589.90594 | 1295.45661 | 863.97350 | E | 401.21431 | 201.11079 | 134.40962 | 3 |
| 25 | 2686.95870 | 1343.98299 | 896.32442 | P | 272.17172 | 136.58950 | 91.39542 | 2 |
| 26 | | | | R | 175.11895 | 88.06311 | 59.04450 | 1 |

Diagnostic fragment ions detected

### Experiment V (lab code 18850)

### KRsIVDGASSETcEDTqSVGSSESSEPR (S459)

### <u>Medium confidence level</u>

OOPS-1

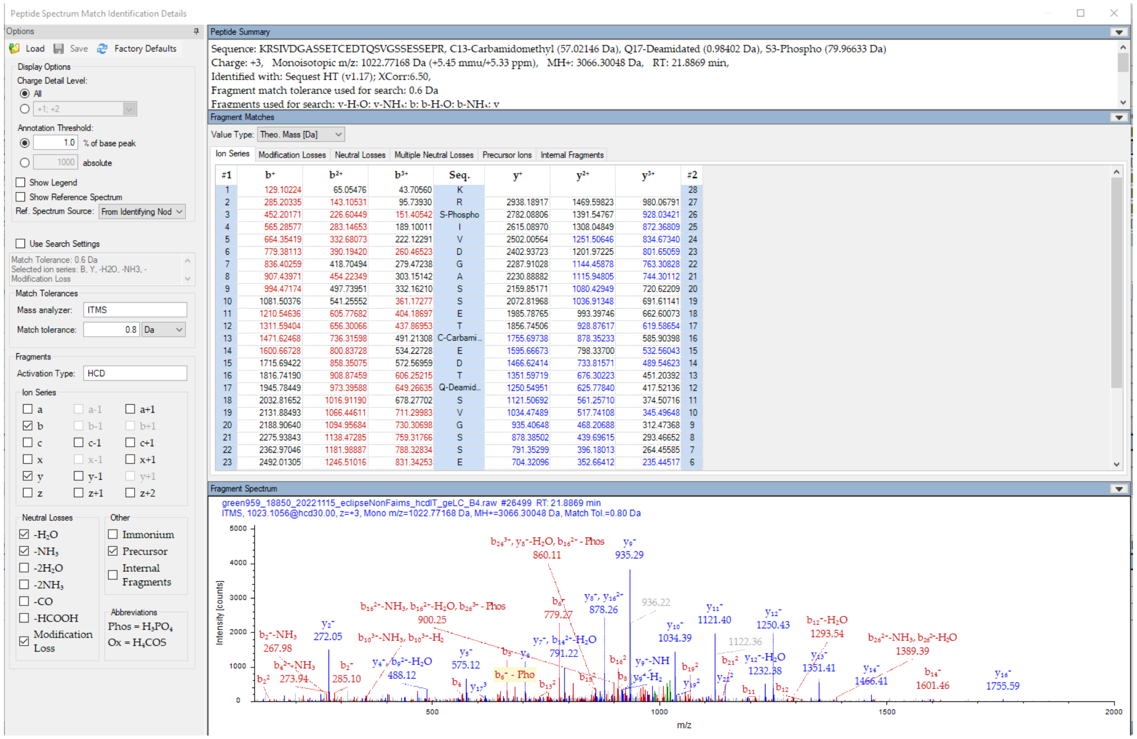


Diagnostic fragment ions detected with low intensity, some indistinguishable from other fragment ions of same m/z

### Experiment V (code 18850)
### KRSIVDGASsETcEDTqSVGSSESSEPR (S466)
### <u>Medium confidence level</u>

OOPS-1

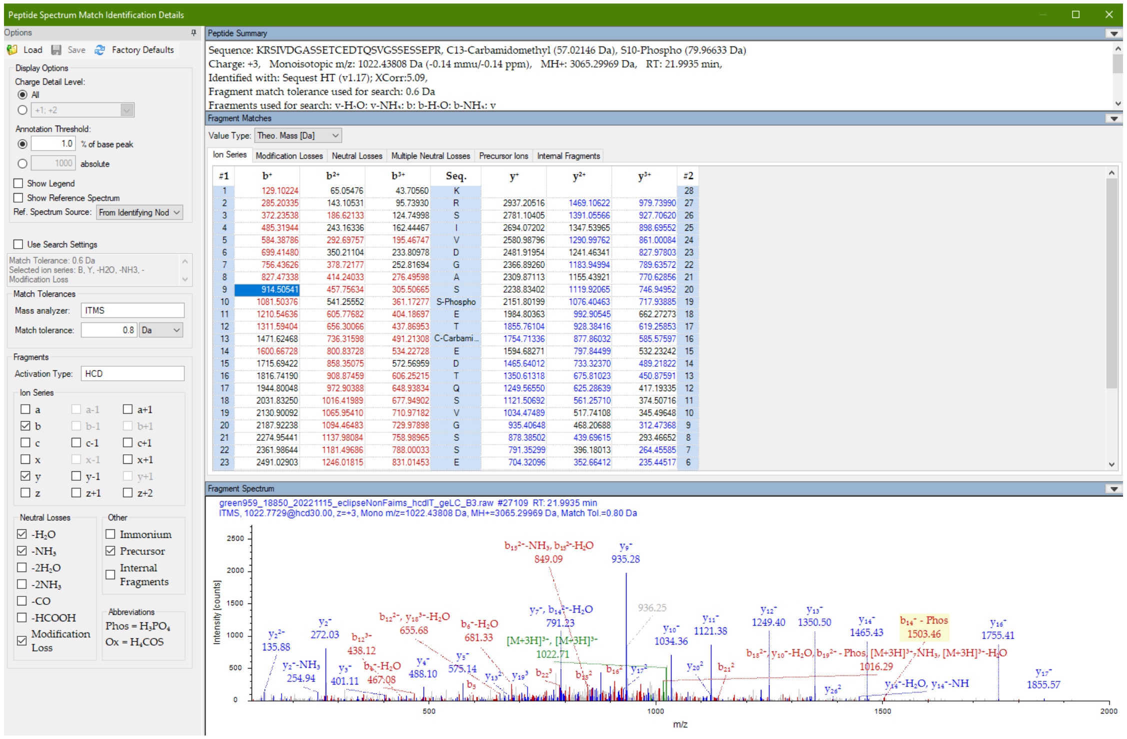


Diagnostic fragment ions detected with low intensity, some indistinguishable from other fragment ions of same m/z

### Experiment V (lab code 18850)
### SIVDGASSETcEDTQSVGSsESSEPR (S478)
### <u>Medium confidence level</u>

OOPS-1

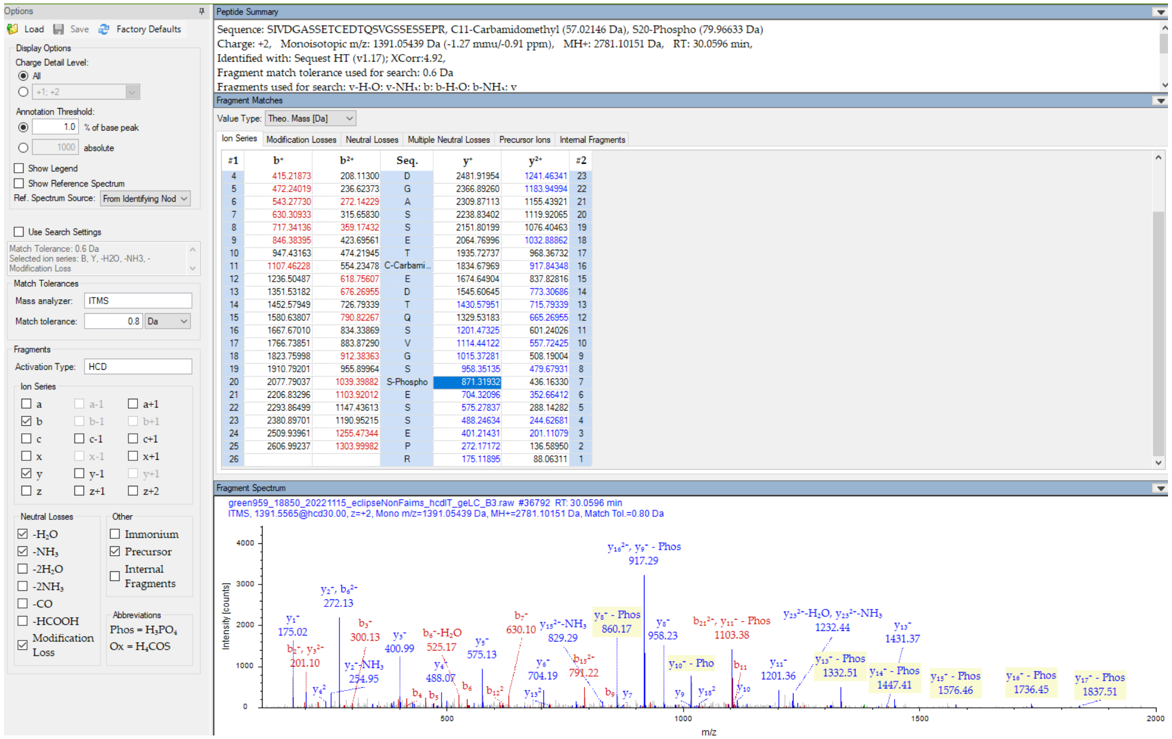


Diagnostic fragment ion 871 *m/z* in the noise; cannot discern between S478 and S477 localization; no supporting b ions

### Experiment V (18850)
### KRSIVDGASsETcEDTqsVGSSESSEPR (S466, S474)
### Low confidence level

OOPS-1

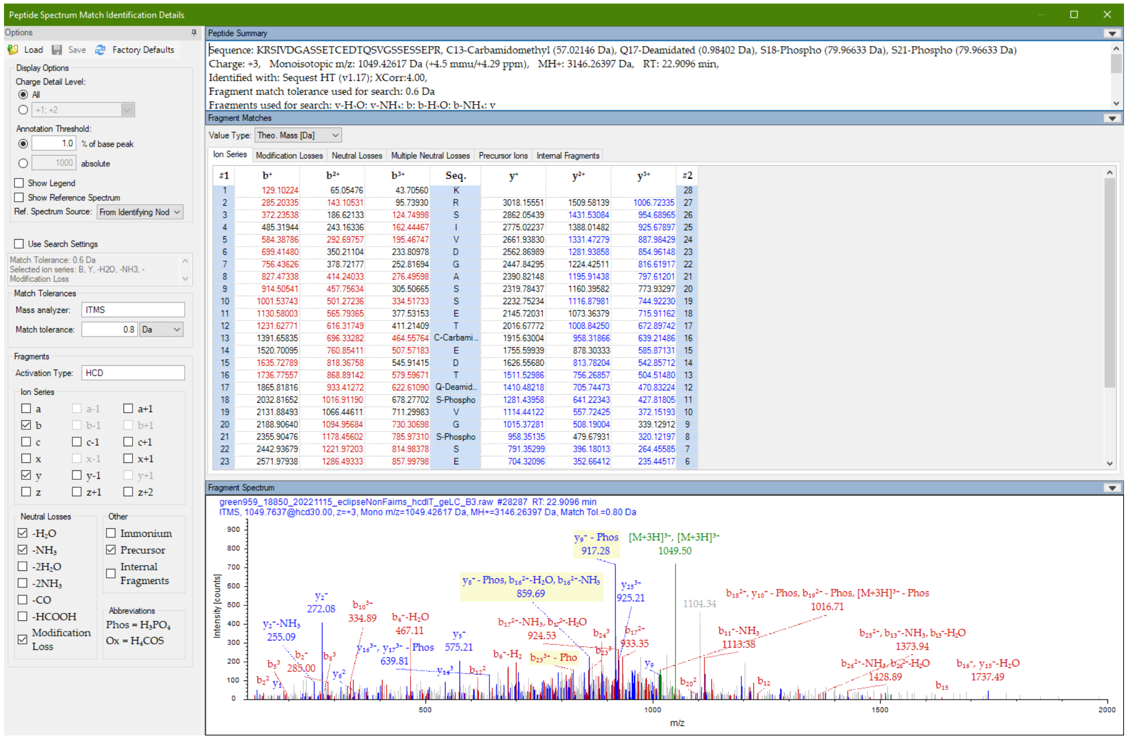


Diagnostic fragment ions detected with low intensity, some indistinguishable from other fragment ions of same m/z

### Experiment V (lab code 18850)
### SIVDGASSETcEDTQSVGSSESsEPR (S481)
### <u>Low confidence level</u>

#### OOPS-1

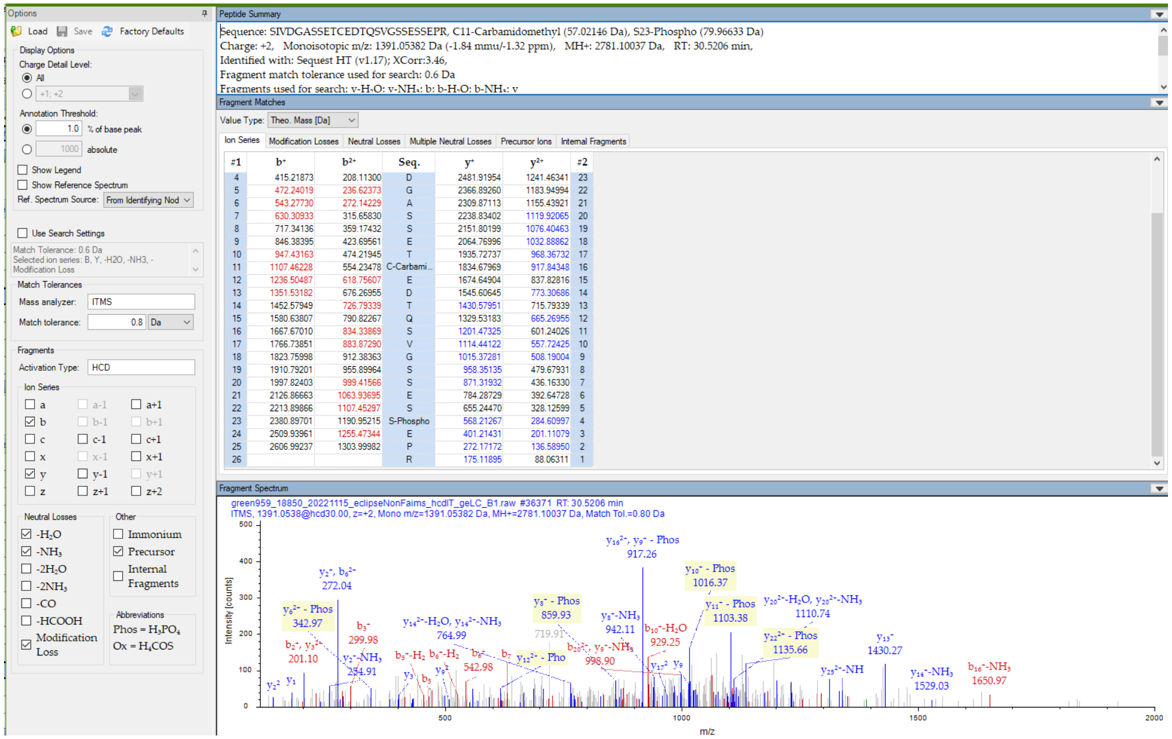


| #1 | $b^+$ | $b^{2+}$ | Seq. | $y^+$ | $y^{2+}$ | #2 |
|---|---|---|---|---|---|---|
| 4 | 415.21873 | 208.11300 | D | 2481.91954 | 1241.46341 | 23 |
| 5 | 472.24019 | 236.62373 | G | 2366.89260 | 1183.94994 | 22 |
| 6 | 543.27730 | 272.14229 | A | 2309.87113 | 1155.43921 | 21 |
| 7 | 630.30933 | 315.65830 | S | 2238.83402 | 1119.92065 | 20 |
| 8 | 717.34136 | 359.17432 | S | 2151.80199 | 1076.40463 | 19 |
| 9 | 846.38395 | 423.69561 | E | 2064.76996 | 1032.88862 | 18 |
| 10 | 947.43163 | 474.21945 | T | 1935.72737 | 968.36732 | 17 |
| 11 | 1107.46228 | 554.23478 | C-Carbami... | 1834.67969 | 917.84348 | 16 |
| 12 | 1236.50487 | 618.75607 | E | 1674.64904 | 837.82816 | 15 |
| 13 | 1351.53182 | 676.26955 | D | 1545.60645 | 773.30686 | 14 |
| 14 | 1452.57949 | 726.79339 | T | 1430.57951 | 715.79339 | 13 |
| 15 | 1580.63807 | 790.82267 | Q | 1329.53183 | 665.26955 | 12 |
| 16 | 1667.67010 | 834.33869 | S | 1201.47325 | 601.24026 | 11 |
| 17 | 1766.73851 | 883.87290 | V | 1114.44122 | 557.72425 | 10 |
| 18 | 1823.75998 | 912.38363 | G | 1015.37281 | 508.19004 | 9 |
| 19 | 1910.79201 | 955.89964 | S | 958.35135 | 479.67931 | 8 |
| 20 | 1997.82403 | 999.41566 | S | 871.31932 | 436.16330 | 7 |
| 21 | 2126.86663 | 1063.93695 | E | 784.28729 | 392.64728 | 6 |
| 22 | 2213.89866 | 1107.45297 | S | 655.24470 | 328.12599 | 5 |
| 23 | 2380.89701 | 1190.95215 | S-Phospho | 568.21267 | 284.60997 | 4 |
| 24 | 2509.93961 | 1255.47344 | E | 401.21431 | 201.11079 | 3 |
| 25 | 2606.99237 | 1303.99982 | P | 272.17172 | 136.58950 | 2 |
| 26 | | | R | 175.11895 | 88.06311 | 1 |

Diagnostic fragment ions in the noise

### Experiment V (lab code 18850)
### SIVDGASSETcEDTQSVGSSEssEPR (S480, S481)
### <u>Low confidence level</u>

OOPS-1

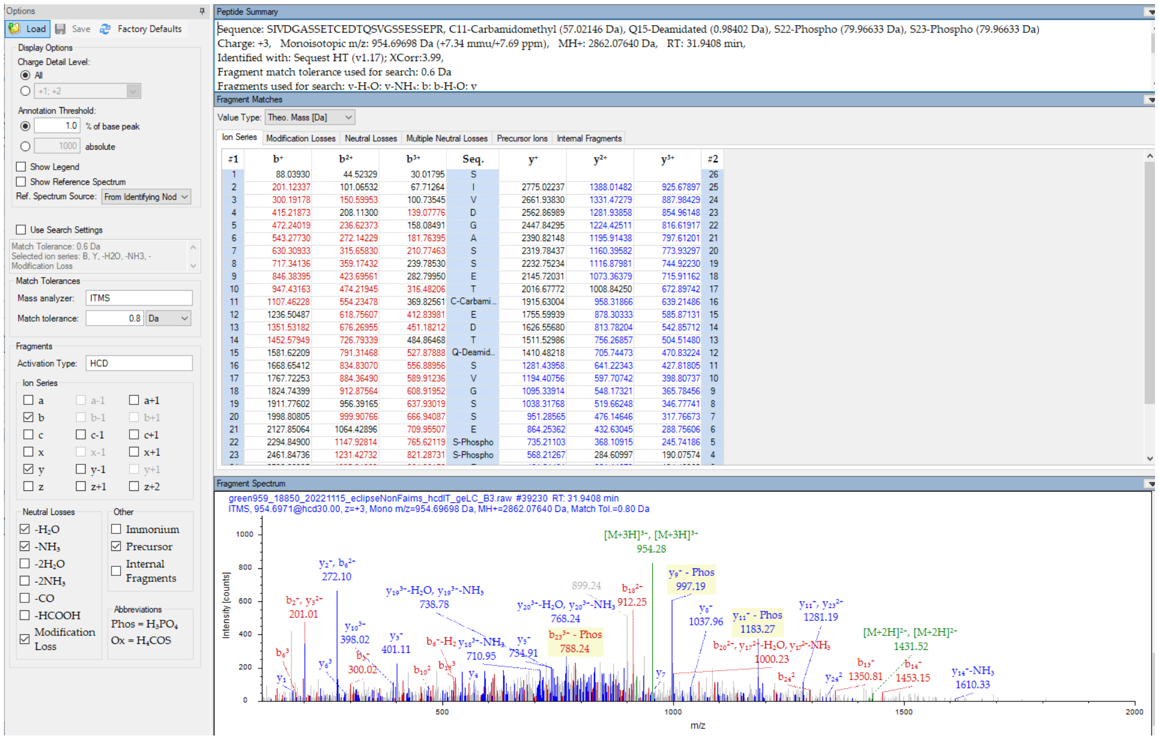


Diagnostic fragment ions in the noise

### Experiment V (lab code 18850)
### SIVDGASSEtcEDTQSVGSSESsEPR (T468, S481)
### <u>Low confidence level</u>

OOPS-1

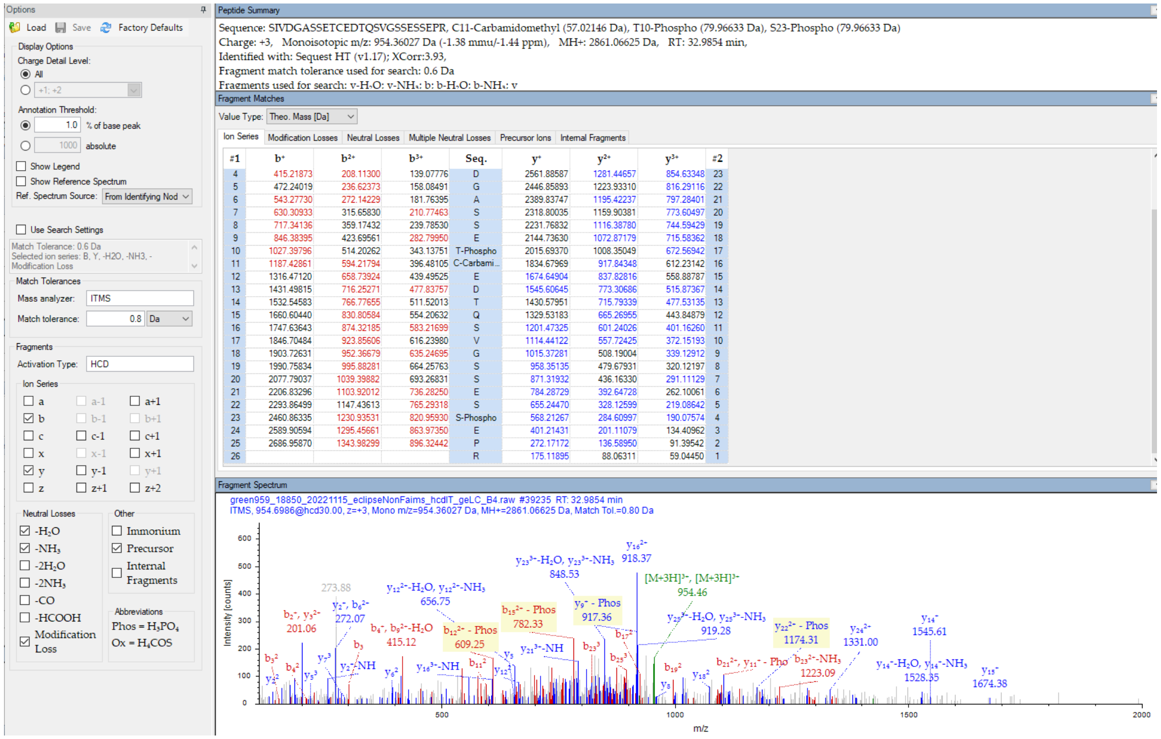


| #1 | b+ | b2+ | b3+ | Seq. | y+ | y2+ | y3+ | #2 |
|---|---|---|---|---|---|---|---|---|
| 4 | 415.21873 | 208.11300 | 139.07776 | D | 2561.88587 | 1281.44657 | 854.63348 | 23 |
| 5 | 472.24019 | 236.62373 | 158.08491 | G | 2446.85893 | 1223.93310 | 816.29116 | 22 |
| 6 | 543.27730 | 272.14229 | 181.76395 | A | 2389.83747 | 1195.42237 | 797.28401 | 21 |
| 7 | 630.30933 | 315.65830 | 210.77463 | S | 2318.80035 | 1159.90381 | 773.60497 | 20 |
| 8 | 717.34136 | 359.17432 | 239.78530 | S | 2231.76832 | 1116.38780 | 744.59429 | 19 |
| 9 | 846.38395 | 423.69561 | 282.79950 | E | 2144.73630 | 1072.87179 | 715.58362 | 18 |
| 10 | 1027.39796 | 514.20262 | 343.13751 | T-Phospho | 2015.69370 | 1008.35049 | 672.56942 | 17 |
| 11 | 1187.42861 | 594.21794 | 396.48105 | C-Carbami... | 1834.67969 | 917.84348 | 612.23142 | 16 |
| 12 | 1316.47120 | 658.73924 | 439.49525 | E | 1674.64904 | 837.82816 | 558.88787 | 15 |
| 13 | 1431.49815 | 716.25271 | 477.83757 | D | 1545.60645 | 773.30686 | 515.87367 | 14 |
| 14 | 1532.54583 | 766.77655 | 511.52013 | T | 1430.57951 | 715.79339 | 477.53135 | 13 |
| 15 | 1660.60440 | 830.80584 | 554.20632 | Q | 1329.53183 | 665.26955 | 443.84879 | 12 |
| 16 | 1747.63643 | 874.32185 | 583.21699 | S | 1201.47325 | 601.24026 | 401.16260 | 11 |
| 17 | 1846.70484 | 923.85606 | 616.23980 | V | 1114.44122 | 557.72425 | 372.15193 | 10 |
| 18 | 1903.72631 | 952.36679 | 635.24695 | G | 1015.37281 | 508.19004 | 339.12912 | 9 |
| 19 | 1990.75834 | 995.88281 | 664.25763 | S | 958.35135 | 479.67931 | 320.12197 | 8 |
| 20 | 2077.79037 | 1039.39882 | 693.26831 | S | 871.31932 | 436.16330 | 291.11129 | 7 |
| 21 | 2206.83296 | 1103.92012 | 736.28250 | E | 784.28729 | 392.64728 | 262.10061 | 6 |
| 22 | 2293.86499 | 1147.43613 | 765.29318 | S | 655.24470 | 328.12599 | 219.08642 | 5 |
| 23 | 2460.86335 | 1230.93531 | 820.95930 | S-Phospho | 568.21267 | 284.60997 | 190.07574 | 4 |
| 24 | 2589.90594 | 1295.45661 | 863.97350 | E | 401.21431 | 201.11079 | 134.40962 | 3 |
| 25 | 2686.95870 | 1343.98299 | 896.32442 | P | 272.17172 | 136.58950 | 91.39542 | 2 |
| 26 | | | | R | 175.11895 | 88.06311 | 59.04450 | 1 |

Diagnostic fragment ions in the noise

### SPE-11

Peptide tandem MS inspection for phosphorylation sites

Spermatocyte protein spe-11 OS=Caenorhabditis elegans OX=6239 GN=spe-11 PE=2 SV=1

☐ Annotate PTMs reported in Uniprot
☐ Show only PTMs
☐ Include PSMs that are Filtered Out

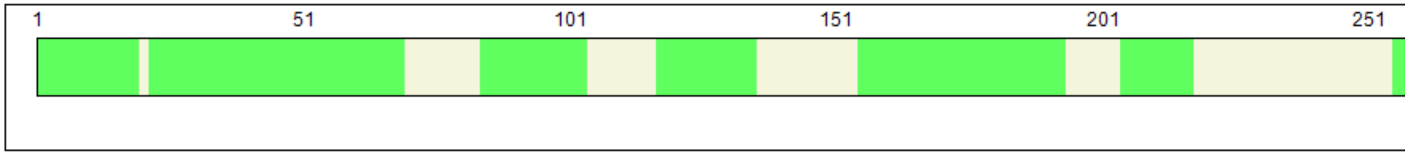


**Coverage: 61.20%**

**Found Modifications:**

**C** Carbamidomethyl (C)
**D** Deamidated (N,Q)
**O** Oxidation (M)
**P** Phospho (S,Y)

[Partial list, from 18850]

Sequence | Modification List

| Position | Target | Modification | Classification | Highest Peptide Confidence | Sequence Motif |
|---|---|---|---|---|---|
| 1 | M | Oxidation | Artefact | High | mSDEEID |
| 13 | N | Deamidated | Artefact | High | DISTALnNKTTPK |
| 14 | N | Deamidated | Artefact | High | ISTALNnKTTPKK |
| 22 | S | Phospho | Post-translational | High | TTPKKKsLKRNSN |
| 26 | N | Deamidated | Artefact | High | KKSLKRnSNSQEG |
| 27 | S | Phospho | Post-translational | High | KSLKRNsNSQEGY |
| 28 | N | Deamidated | Artefact | High | SLKRNSnSQEGYE |
| 29 | S | Phospho | Post-translational | High | LKRNSNsQEGYES |
| 30 | Q | Deamidated | Artefact | High | KRNSNSqEGYESP |
| 33 | Y | Phospho | Post-translational | High | SNSQEGyESPEER |
| 35 | S | Phospho | Post-translational | High | SQEGYEsPEEREI |
| 54 | M | Oxidation | Artefact | High | GAIGTPmAKSDNA |
| 88 | M | Oxidation | Artefact | High | LRSKWDmTQGHLP |
| 117 | M | Oxidation | Artefact | High | YNSKARmDILDGL |
| 133 | C | Carbamidomethyl | Chemical derivative | High | NEGFFNcGKGAAM |
| 164 | M | Oxidation | Artefact | High | ATVETAmKKAGNP |
| 169 | N | Deamidated | Artefact | High | AMKKAGnPTMEQM |
| 172 | M | Oxidation | Artefact | High | KAGNPTmEQMMTD |
| 174 | Q | Deamidated | Artefact | High | GNPTMEqMMTDDL |
| 175 | M | Oxidation | Artefact | High | NPTMEQmMTDDLD |
| 176 | M | Oxidation | Artefact | High | PTMEQMmTDDLDE |
| 209 | M | Oxidation | Artefact | High | RAYDAAmDEREDD |
| 260 | N | Deamidated | Artefact | High | KVTHKFnAYQLDL |
| 263 | Q | Deamidated | Artefact | High | HKFNAYqLDLKCL |
| 268 | C | Carbamidomethyl | Chemical derivative | High | YQLDLKcLDEDAF |
| 276 | N | Deamidated | Artefact | High | DEDAFSnKKSLKS |

### Experiment VI (lab code 18879)
### MsDEEIDISTALNNKTTPK (S2)
### High confidence level

SPE-11

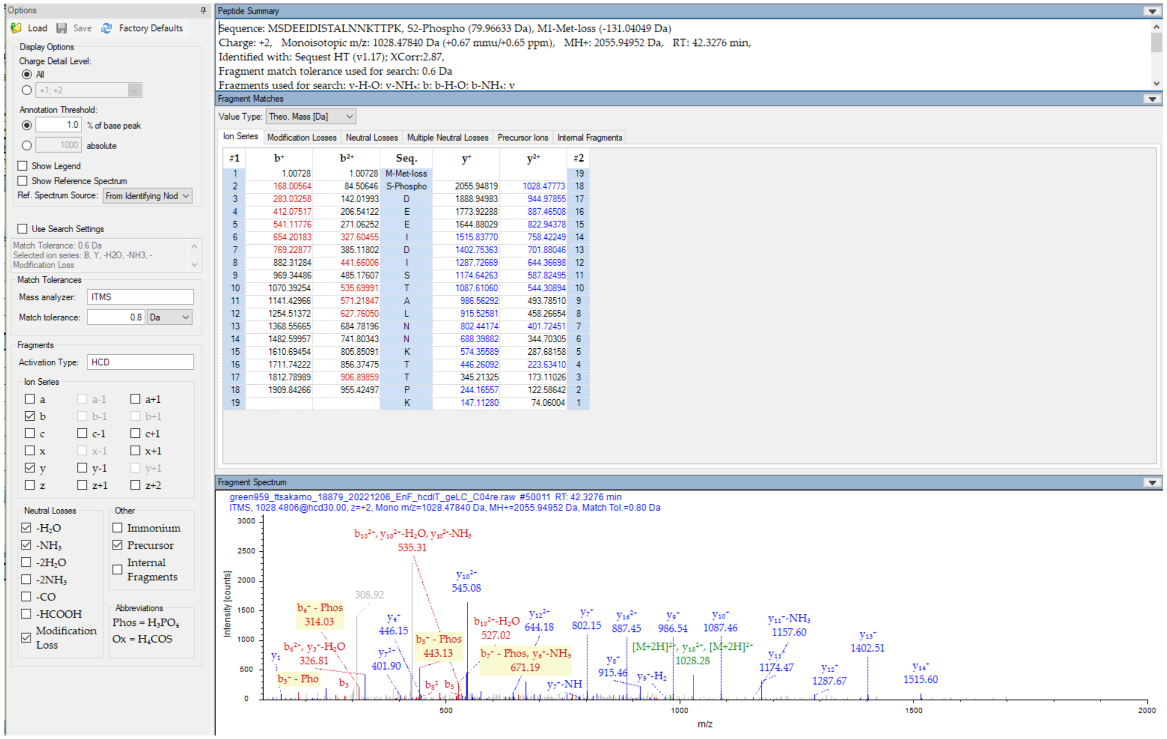


Diagnostic fragment ions detected (y-1 to y-14 support no phos from C-term)

### Experiment VI (lab code 18879)
### MSDEEIDISTALNNKTtPK (S2)
### <u>High confidence level</u>

SPE-11

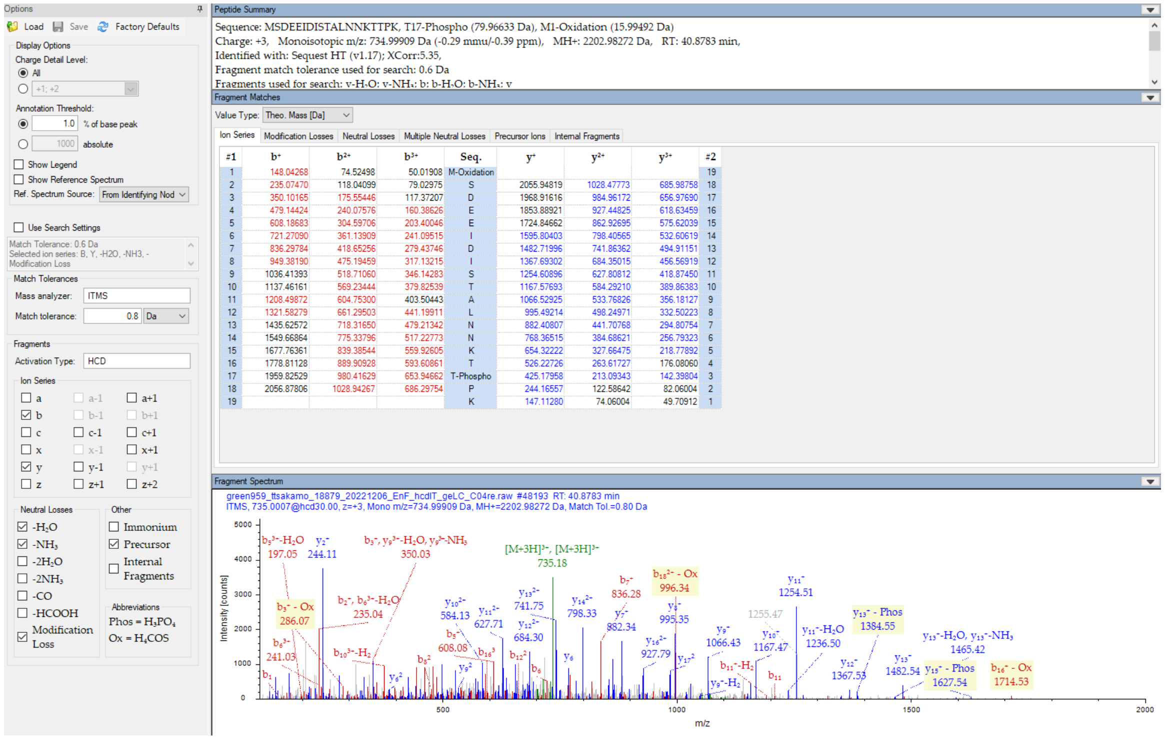


Diagnostic fragment ions detected

### Experiment VI (lab code 18879)
### RNsNSQEGYESPEER (S27)
### <u>High confidence level</u>

SPE-11

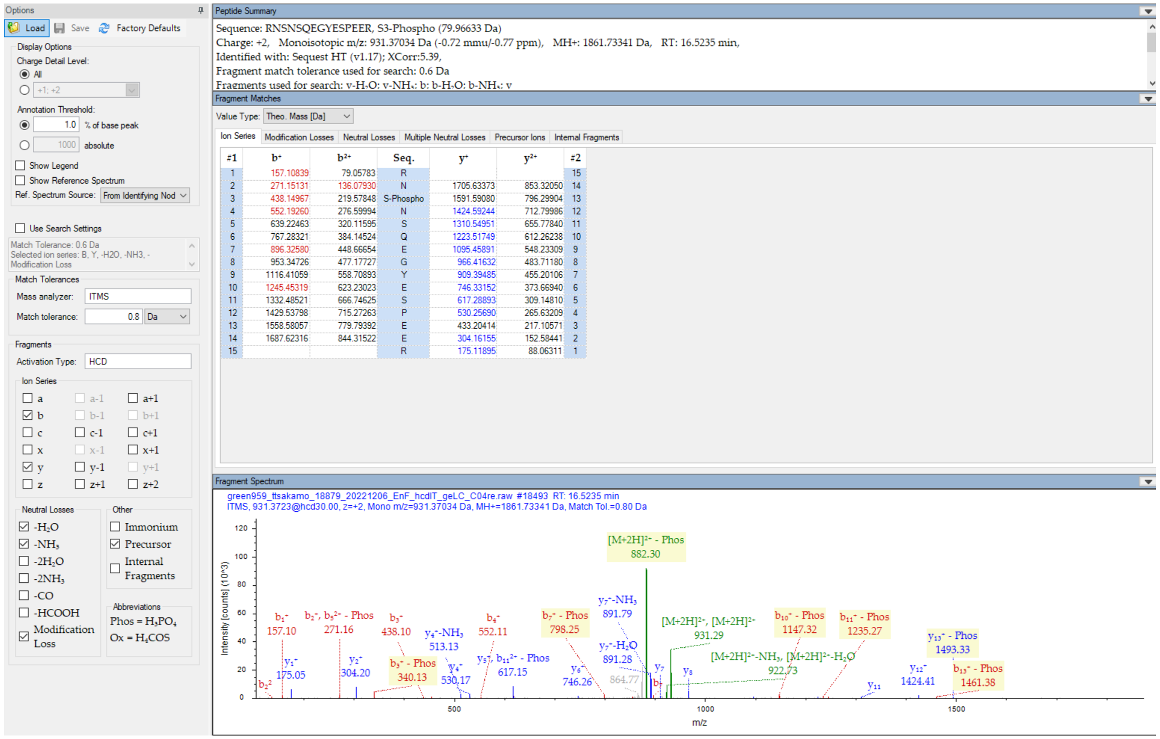


| #1 | b+ | b2+ | Seq. | y+ | y2+ | #2 |
|---|---|---|---|---|---|---|
| 1 | 157.10839 | 79.05783 | R | | | 15 |
| 2 | 271.15131 | 136.07930 | N | 1705.63373 | 853.32050 | 14 |
| 3 | 438.14967 | 219.57848 | S-Phospho | 1591.59080 | 796.29904 | 13 |
| 4 | 552.19260 | 276.59994 | N | 1424.59244 | 712.79986 | 12 |
| 5 | 639.22463 | 320.11595 | S | 1310.54951 | 655.77840 | 11 |
| 6 | 767.28321 | 384.14524 | Q | 1223.51749 | 612.26238 | 10 |
| 7 | 896.32580 | 448.66654 | E | 1095.45891 | 548.23309 | 9 |
| 8 | 953.34726 | 477.17727 | G | 966.41632 | 483.71180 | 8 |
| 9 | 1116.41059 | 558.70893 | Y | 909.39485 | 455.20106 | 7 |
| 10 | 1245.45319 | 623.23023 | E | 746.33152 | 373.66940 | 6 |
| 11 | 1332.48521 | 666.74625 | S | 617.28893 | 309.14810 | 5 |
| 12 | 1429.53798 | 715.27263 | P | 530.25690 | 265.63209 | 4 |
| 13 | 1558.58057 | 779.79392 | E | 433.20414 | 217.10571 | 3 |
| 14 | 1687.62316 | 844.31522 | E | 304.16155 | 152.58441 | 2 |
| 15 | | | R | 175.11895 | 88.06311 | 1 |

Diagnostic fragment ions detected

### Experiment IV (lab code 18822)
### NSNsQEGYESPEER (S29)
### High confidence level

#### SPE-11

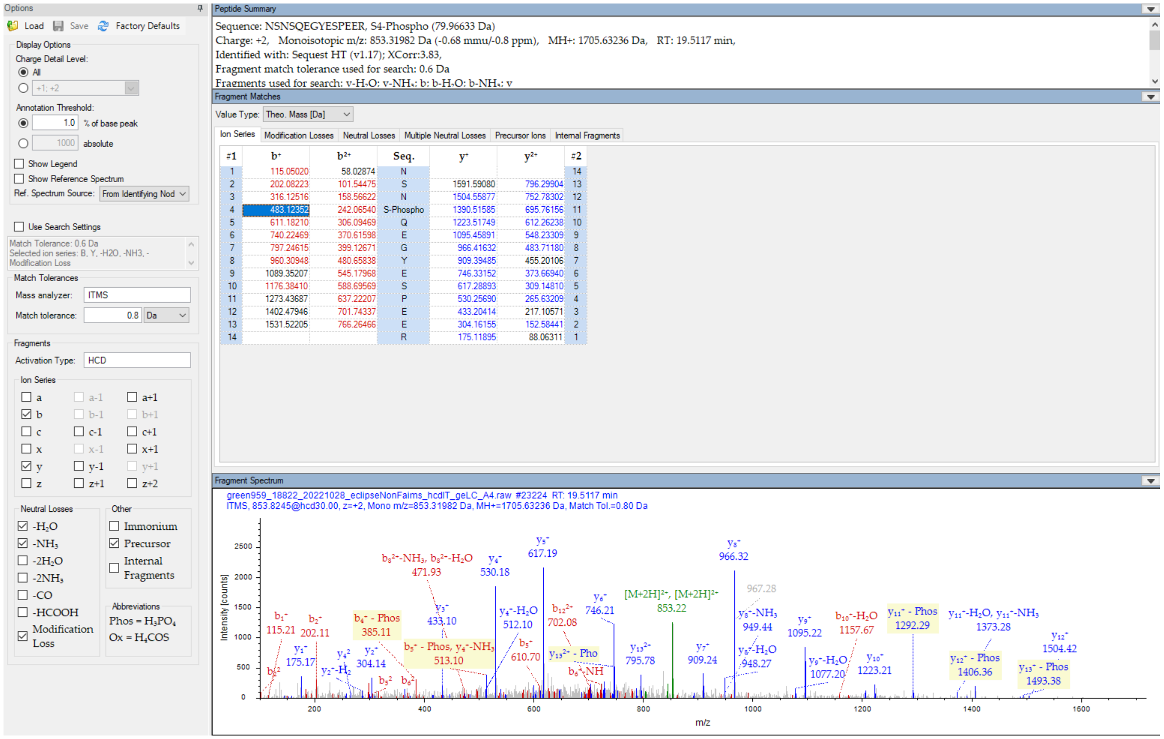


Diagnostic fragment ions detected

### Experiment VII (lab code 18902)
### RNsNsQEGYESPEER (S27, S29)
### <u>High confidence level</u>

SPE-11

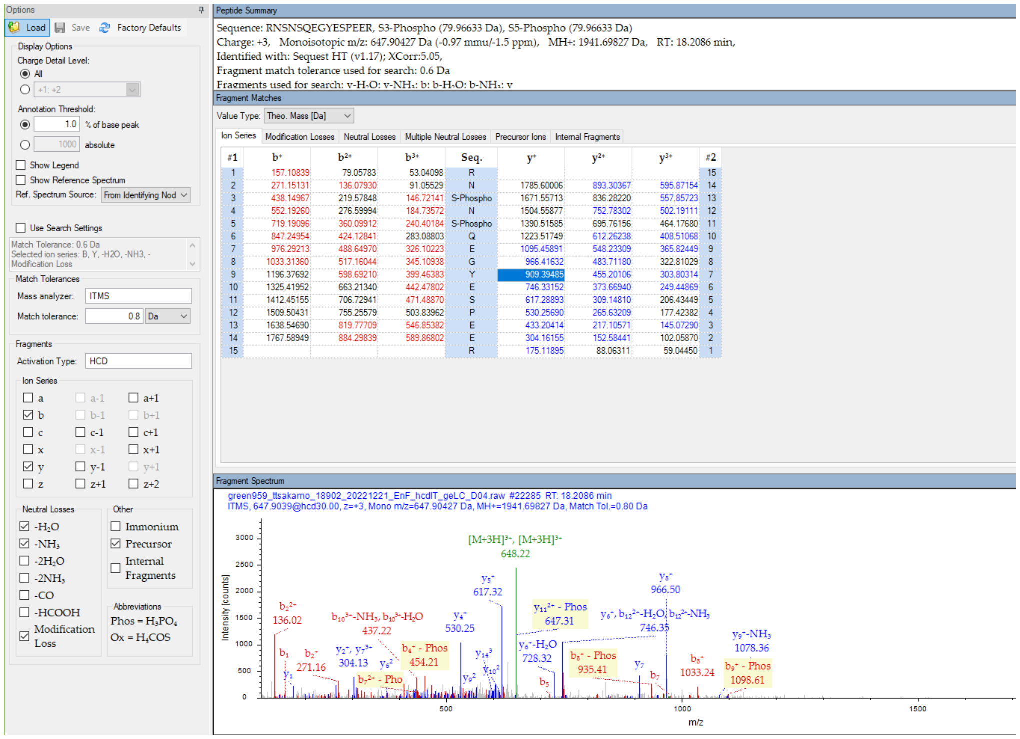


Diagnostic fragment ions detected

### Experiment VI (lab code 18879)
### RNSNSQEGYEsPEER (S35)
### <u>High confidence level</u>

SPE-11

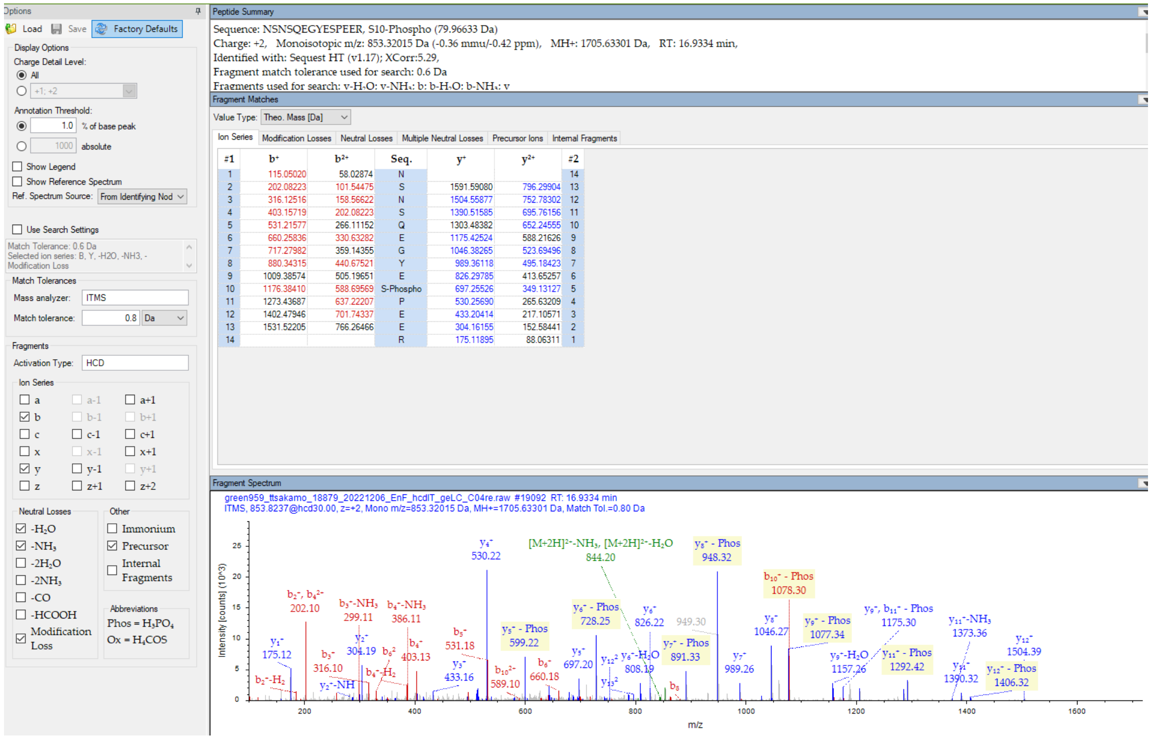


| #1 | $b^+$ | $b^{2+}$ | Seq. | $y^+$ | $y^{2+}$ | #2 |
|---|---|---|---|---|---|---|
| 1 | 115.05020 | 58.02874 | N | | | 14 |
| 2 | 202.08223 | 101.54475 | S | 1591.59080 | 796.29904 | 13 |
| 3 | 316.12516 | 158.56622 | N | 1504.55877 | 752.78302 | 12 |
| 4 | 403.15719 | 202.08223 | S | 1390.51585 | 695.76156 | 11 |
| 5 | 531.21577 | 266.11152 | Q | 1303.48382 | 652.24555 | 10 |
| 6 | 660.25836 | 330.63282 | E | 1175.42524 | 588.21626 | 9 |
| 7 | 717.27982 | 359.14355 | G | 1046.38265 | 523.69496 | 8 |
| 8 | 880.34315 | 440.67521 | Y | 989.36118 | 495.18423 | 7 |
| 9 | 1009.38574 | 505.19651 | E | 826.29785 | 413.65257 | 6 |
| 10 | 1176.38410 | 588.69569 | S-Phospho | 697.25526 | 349.13127 | 5 |
| 11 | 1273.43687 | 637.22207 | P | 530.25690 | 265.63209 | 4 |
| 12 | 1402.47946 | 701.74337 | E | 433.20414 | 217.10571 | 3 |
| 13 | 1531.52205 | 766.26466 | E | 304.16155 | 152.58441 | 2 |
| 14 | | | R | 175.11895 | 88.06311 | 1 |

Diagnostic fragment ions detected

### Experiment V (lab code 18850)
### RNSNsQEGYEsPEER (S29, S35)
### <u>High confidence level</u>

SPE-11

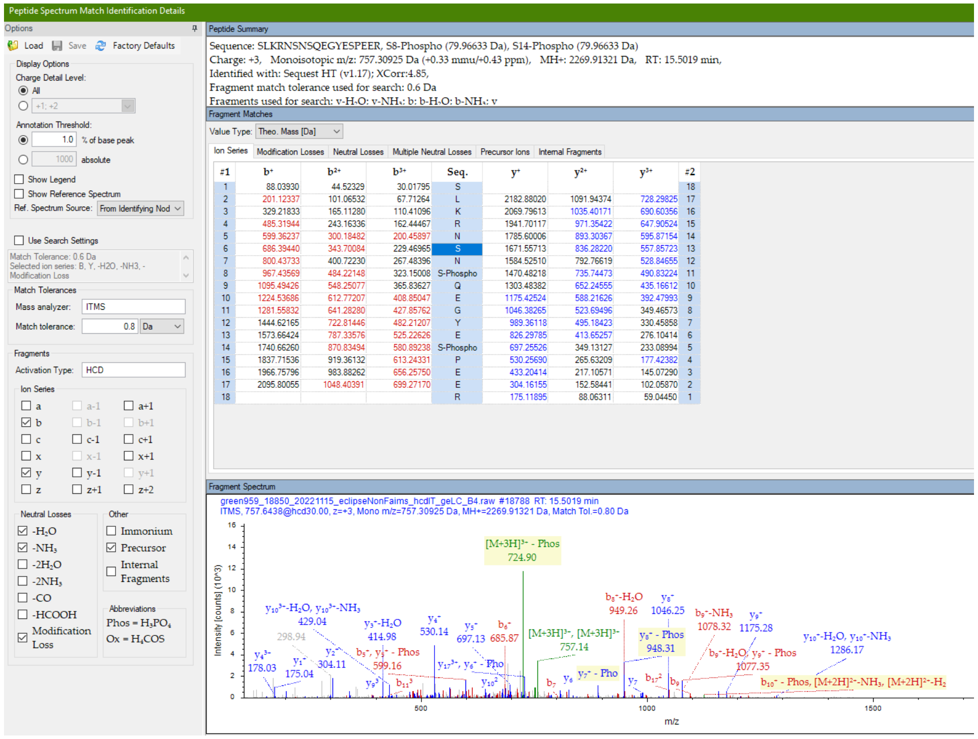


| #1 | $b^+$ | $b^{2+}$ | $b^{3+}$ | Seq. | $y^+$ | $y^{2+}$ | $y^{3+}$ | #2 |
|---|---|---|---|---|---|---|---|---|
| 1 | 88.03930 | 44.52329 | 30.01795 | S | | | | 18 |
| 2 | 201.12337 | 101.06532 | 67.71264 | L | 2182.88020 | 1091.94374 | 728.29825 | 17 |
| 3 | 329.21833 | 165.11280 | 110.41096 | K | 2069.79613 | 1035.40171 | 690.60356 | 16 |
| 4 | 485.31944 | 243.16336 | 162.44467 | R | 1941.70117 | 971.35422 | 647.90524 | 15 |
| 5 | 599.36237 | 300.18482 | 200.45897 | N | 1785.60006 | 893.30367 | 595.87154 | 14 |
| 6 | 686.39440 | 343.70084 | 229.46965 | S | 1671.55713 | 836.28220 | 557.85723 | 13 |
| 7 | 800.43733 | 400.72230 | 267.48396 | N | 1584.52510 | 792.76619 | 528.84655 | 12 |
| 8 | 967.43569 | 484.22148 | 323.15008 | S-Phospho | 1470.48218 | 735.74473 | 490.83224 | 11 |
| 9 | 1095.49426 | 548.25077 | 365.83627 | Q | 1303.48382 | 652.24555 | 435.16612 | 10 |
| 10 | 1224.53686 | 612.77207 | 408.85047 | E | 1175.42524 | 588.21626 | 392.47993 | 9 |
| 11 | 1281.55832 | 641.28280 | 427.85762 | G | 1046.38265 | 523.69496 | 349.46573 | 8 |
| 12 | 1444.62165 | 722.81446 | 482.21207 | Y | 989.36118 | 495.18423 | 330.45858 | 7 |
| 13 | 1573.66424 | 787.33576 | 525.22626 | E | 826.29785 | 413.65257 | 276.10414 | 6 |
| 14 | 1740.66260 | 870.83494 | 580.89238 | S-Phospho | 697.25526 | 349.13127 | 233.08994 | 5 |
| 15 | 1837.71536 | 919.36132 | 613.24331 | P | 530.25690 | 265.63209 | 177.42382 | 4 |
| 16 | 1966.75796 | 983.88262 | 656.25750 | E | 433.20414 | 217.10571 | 145.07290 | 3 |
| 17 | 2095.80055 | 1048.40391 | 699.27170 | E | 304.16155 | 152.58441 | 102.05870 | 2 |
| 18 | | | | R | 175.11895 | 88.06311 | 59.04450 | 1 |

Diagnostic fragment ions detected

### Experiment V (lab code 18850)
### RNsNsQEGYEsPEER (S27, S29, S35)
### <u>High confidence level</u>

SPE-11

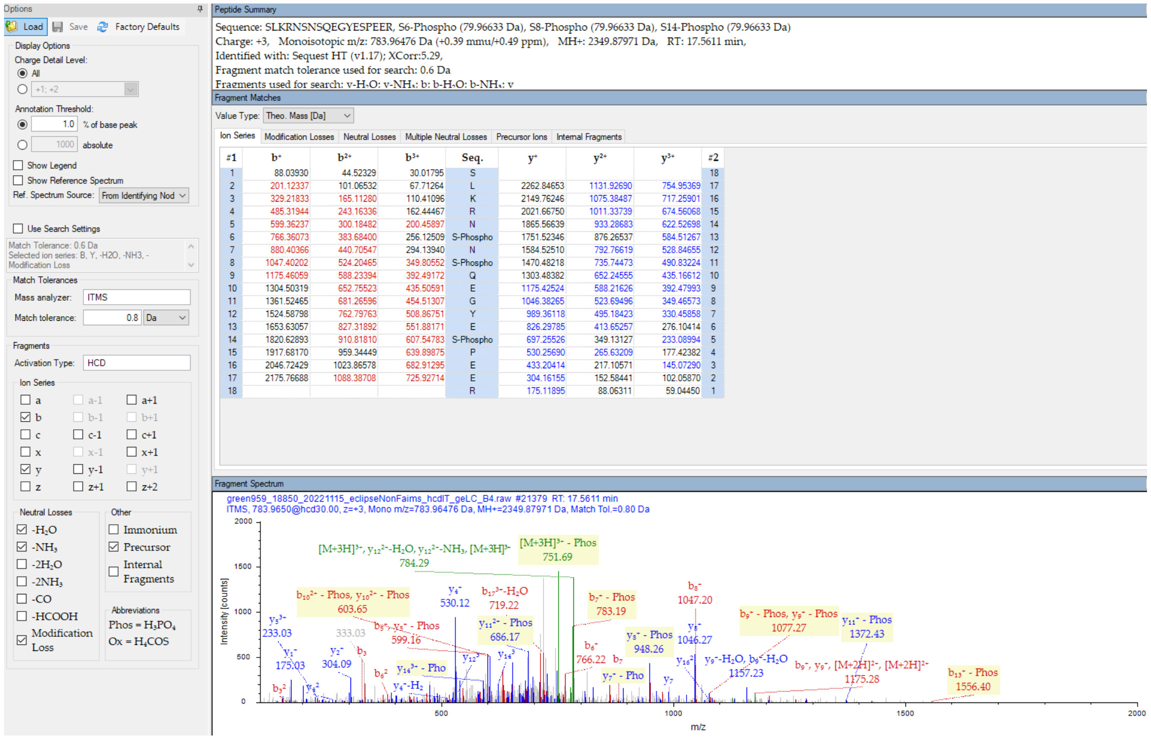


| #1 | $b^+$ | $b^{2+}$ | $b^{3+}$ | Seq. | $y^+$ | $y^{2+}$ | $y^{3+}$ | #2 |
|---|---|---|---|---|---|---|---|---|
| 1 | 88.03930 | 44.52329 | 30.01795 | S | | | | 18 |
| 2 | 201.12337 | 101.06532 | 67.71264 | L | 2262.84653 | 1131.92690 | 754.95369 | 17 |
| 3 | 329.21833 | 165.11280 | 110.41096 | K | 2149.76246 | 1075.38487 | 717.25901 | 16 |
| 4 | 485.31944 | 243.16336 | 162.44467 | R | 2021.66750 | 1011.33739 | 674.56068 | 15 |
| 5 | 599.36237 | 300.18482 | 200.45897 | N | 1865.56639 | 933.28683 | 622.52698 | 14 |
| 6 | 766.36073 | 383.68400 | 256.12509 | S-Phospho | 1751.52346 | 876.26537 | 584.51267 | 13 |
| 7 | 880.40366 | 440.70547 | 294.13940 | N | 1584.52510 | 792.76619 | 528.84655 | 12 |
| 8 | 1047.40202 | 524.20465 | 349.80552 | S-Phospho | 1470.48218 | 735.74473 | 490.83224 | 11 |
| 9 | 1175.46059 | 588.23394 | 392.49172 | Q | 1303.48382 | 652.24555 | 435.16612 | 10 |
| 10 | 1304.50319 | 652.75523 | 435.50591 | E | 1175.42524 | 588.21626 | 392.47993 | 9 |
| 11 | 1361.52465 | 681.26596 | 454.51307 | G | 1046.38265 | 523.69496 | 349.46573 | 8 |
| 12 | 1524.58798 | 762.79763 | 508.86751 | Y | 989.36118 | 495.18423 | 330.45858 | 7 |
| 13 | 1653.63057 | 827.31892 | 551.88171 | E | 826.29785 | 413.65257 | 276.10414 | 6 |
| 14 | 1820.62893 | 910.81810 | 607.54783 | S-Phospho | 697.25526 | 349.13127 | 233.08994 | 5 |
| 15 | 1917.68170 | 959.34449 | 639.89875 | P | 530.25690 | 265.63209 | 177.42382 | 4 |
| 16 | 2046.72429 | 1023.86578 | 682.91295 | E | 433.20414 | 217.10571 | 145.07290 | 3 |
| 17 | 2175.76688 | 1088.38708 | 725.92714 | E | 304.16155 | 152.58441 | 102.05870 | 2 |
| 18 | | | | R | 175.11895 | 88.06311 | 59.04450 | 1 |

Diagnostic fragment ions detected

### Experiment VII (lab code 18902)
### AGNPtMEQMMTDDLDEDEARAEAEWER (T171)
### High confidence level

SPE-11

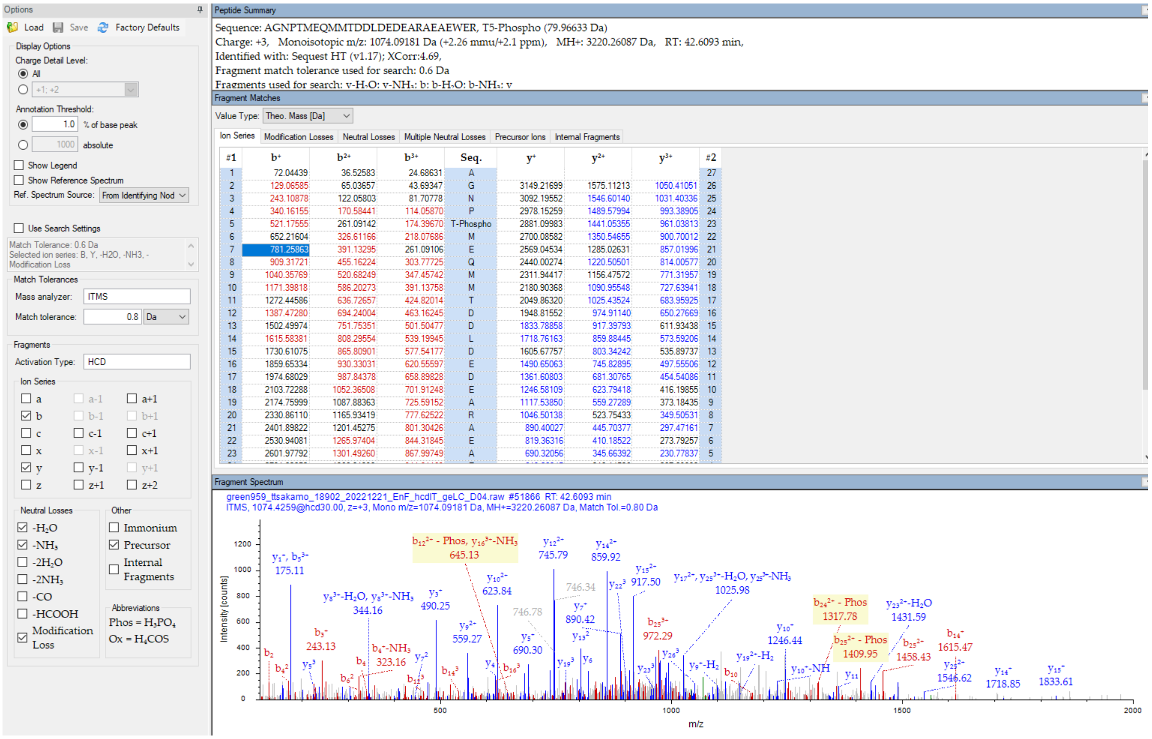


Diagnostic fragment ions detected

### Experiment VI (lab code 18879)

### NSNSQEGyESPEEREIVYPSVFGAIGTPMAK (Y33)

### <u>Medium confidence level</u>

SPE-11

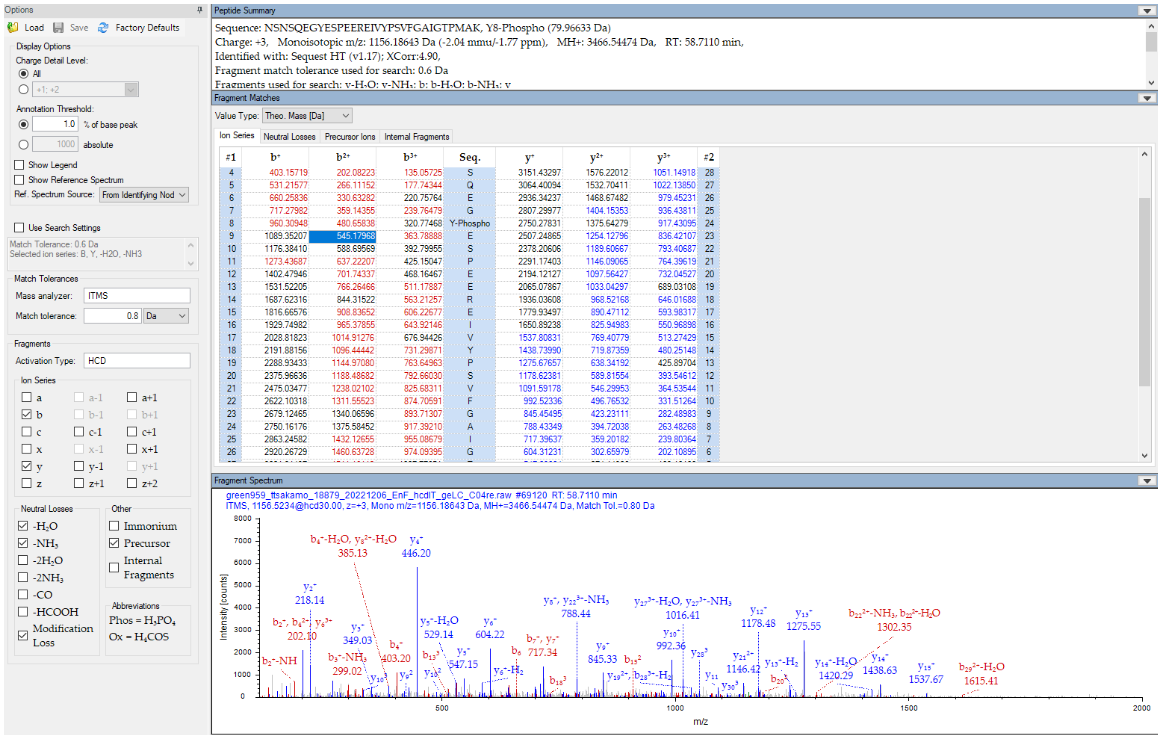


| #1 | b⁺ | b²⁺ | b³⁺ | Seq. | y⁺ | y²⁺ | y³⁺ | #2 |
|---|---|---|---|---|---|---|---|---|
| 4 | 403.15719 | 202.08223 | 135.05725 | S | 3151.43297 | 1576.22012 | 1051.14918 | 28 |
| 5 | 531.21577 | 266.11152 | 177.74344 | Q | 3064.40094 | 1532.70411 | 1022.13850 | 27 |
| 6 | 660.25836 | 330.63282 | 220.75764 | E | 2936.34237 | 1468.67482 | 979.45231 | 26 |
| 7 | 717.27982 | 359.14355 | 239.76479 | G | 2807.29977 | 1404.15353 | 936.43811 | 25 |
| 8 | 960.30948 | 480.65838 | 320.77468 | Y-Phospho | 2750.27831 | 1375.64279 | 917.43095 | 24 |
| 9 | 1089.35207 | 545.17968 | 363.78888 | E | 2507.24865 | 1254.12796 | 836.42107 | 23 |
| 10 | 1176.38410 | 588.69569 | 392.79955 | S | 2378.20606 | 1189.60667 | 793.40687 | 22 |
| 11 | 1273.43687 | 637.22207 | 425.15047 | P | 2291.17403 | 1146.09065 | 764.39619 | 21 |
| 12 | 1402.47946 | 701.74337 | 468.16467 | E | 2194.12127 | 1097.56427 | 732.04527 | 20 |
| 13 | 1531.52205 | 766.26466 | 511.17887 | E | 2065.07867 | 1033.04297 | 689.03108 | 19 |
| 14 | 1687.62316 | 844.31522 | 563.21257 | R | 1936.03608 | 968.52168 | 646.01688 | 18 |
| 15 | 1816.66576 | 908.83652 | 606.22677 | E | 1779.93497 | 890.47112 | 593.98317 | 17 |
| 16 | 1929.74982 | 965.37855 | 643.92146 | I | 1650.89238 | 825.94983 | 550.96898 | 16 |
| 17 | 2028.81823 | 1014.91276 | 676.94426 | V | 1537.80831 | 769.40779 | 513.27429 | 15 |
| 18 | 2191.88156 | 1096.44442 | 731.29871 | Y | 1438.73990 | 719.87359 | 480.25148 | 14 |
| 19 | 2288.93433 | 1144.97080 | 763.64963 | P | 1275.67657 | 638.34192 | 425.89704 | 13 |
| 20 | 2375.96636 | 1188.48682 | 792.66030 | S | 1178.62381 | 589.81554 | 393.54612 | 12 |
| 21 | 2475.03477 | 1238.02102 | 825.68311 | V | 1091.59178 | 546.29953 | 364.53544 | 11 |
| 22 | 2622.10318 | 1311.55523 | 874.70591 | F | 992.52336 | 496.76532 | 331.51264 | 10 |
| 23 | 2679.12465 | 1340.06596 | 893.71307 | G | 845.45495 | 423.23111 | 282.48983 | 9 |
| 24 | 2750.16176 | 1375.58452 | 917.39210 | A | 788.43349 | 394.72038 | 263.48268 | 8 |
| 25 | 2863.24582 | 1432.12655 | 955.08679 | I | 717.39637 | 359.20182 | 239.80364 | 7 |
| 26 | 2920.26729 | 1460.63728 | 974.09395 | G | 604.31231 | 302.65979 | 202.10895 | 6 |

Diagnostic fragment ions detected near the noise; site localization not clearly defined

### Experiment V (lab code 18850)

### sLKRNSNSQEGYESPEER (S22)

### <u>Low confidence level</u>

SPE-11

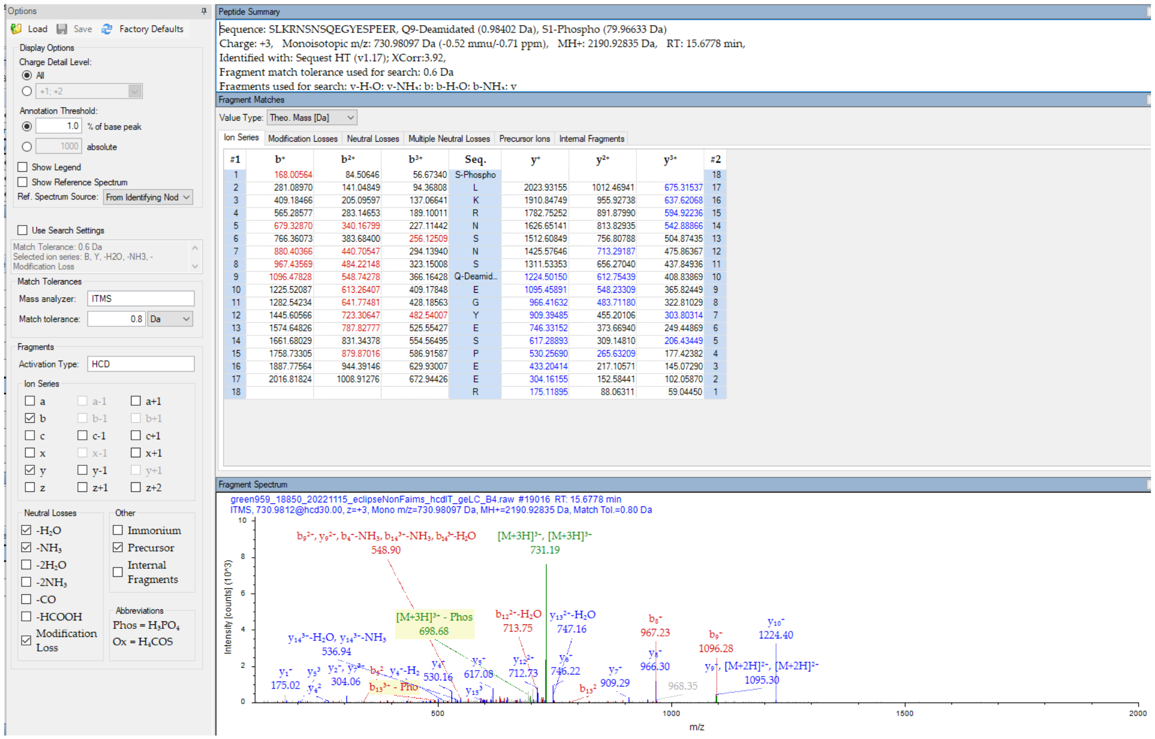


Diagnostic fragment ions in the noise

### Experiment VII (lab code 18902)
### RNSNSQEGyEsPEER (Y33, S35)
### <u>Low confidence level</u>

SPE-11

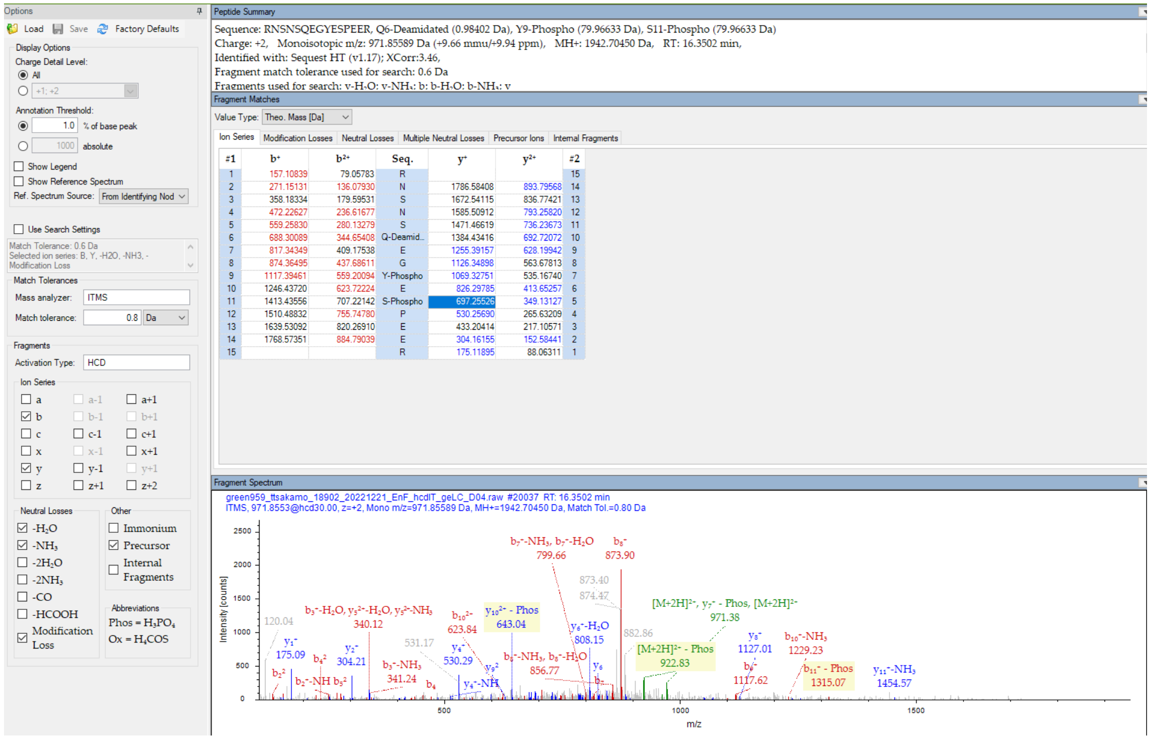


| #1 | b+ | b2+ | Seq. | y+ | y2+ | #2 |
|---|---|---|---|---|---|---|
| 1 | 157.10839 | 79.05783 | R | | | 15 |
| 2 | 271.15131 | 136.07930 | N | 1786.58408 | 893.79568 | 14 |
| 3 | 358.18334 | 179.59531 | S | 1672.54115 | 836.77421 | 13 |
| 4 | 472.22627 | 236.61677 | N | 1585.50912 | 793.25820 | 12 |
| 5 | 559.25830 | 280.13279 | S | 1471.46619 | 736.23673 | 11 |
| 6 | 688.30089 | 344.65408 | Q-Deamid... | 1384.43416 | 692.72072 | 10 |
| 7 | 817.34349 | 409.17538 | E | 1255.39157 | 628.19942 | 9 |
| 8 | 874.36495 | 437.68611 | G | 1126.34898 | 563.67813 | 8 |
| 9 | 1117.39461 | 559.20094 | Y-Phospho | 1069.32751 | 535.16740 | 7 |
| 10 | 1246.43720 | 623.72224 | E | 826.29785 | 413.65257 | 6 |
| 11 | 1413.43556 | 707.22142 | S-Phospho | 697.25526 | 349.13127 | 5 |
| 12 | 1510.48832 | 755.74780 | P | 530.25690 | 265.63209 | 4 |
| 13 | 1639.53092 | 820.26910 | E | 433.20414 | 217.10571 | 3 |
| 14 | 1768.57351 | 884.79039 | E | 304.16155 | 152.58441 | 2 |
| 15 | | | R | 175.11895 | 88.06311 | 1 |

Diagnostic fragment ions in the noise
