## Supplemental File 5 for "A sperm–oocyte protein partnership required for egg activation in *Caenorhabditis* elegans"

### File S5. *C. elegans* strains used in this study

| Strain | Genotype | Comments |
|---|---|---|
| N2 | Wild type, Bristol isolate | |
| AD200 | *unc-119(ed3)* III*; asIs1[pie-1p::gfp::egg-3 + unc-119(+)]* | |
| AD226 | *egg-3(tm1191)/mIn1[mIs14 dpy-10(e128)]* II | |
| AG214 | *unc-119(ed3) III; avIs143[cbd-1p::cbd-1::mCherry + unc-119(+)]* | |
| AJL30 | *oops-1(tn1898)/tmC25* IV*; ltIs37[pie-1p::mCherry::his-58 + unc-119(+)]* IV*; ruIs57[pie-1::gfp::tba-2 + unc-119(+)]* | |
| AJL32 | *spe-11(hc90)/tmC18* I*; ltIs37[pie-1p::mCherry::his-58 + unc-119(+)]* IV*; ruIs57[pie-1::gfp::tba-2 + unc-119(+)]* | |
| AJL56 | *spe-11(tn2059)/tmC18(dpy-5[tmIs1236]) I; him-8 (el489)* IV | |
| AJL66 | *chs-1(ude35[egfp::chs-1])* I | |
| AJL72 | *spe-11(tn2059)/tmC18(dpy-5[tmIs1236])* I*; avIs143[cbd-1p::cbd-1::mCherry + unc-119(+)]* | |
| AJL79 | *chs-1(ude35[egfp::chs-1])* I*; oops-1(tn1898)/tmC25(unc-5[tnIs1241]) IV; him-5(e1490)* V | |
| AJL120 | *unc-119(ed3) III; spe-11(tn2059)/tmC18(dpy-5[tmIs1236])* I*; oops-1(tn1898) ltIs37[unc-119(+) pie-1p::mCherry::H2B])/tmC25[unc-5(tmIs1241)]* IV*;ruIs57[pie-1p::β-tubulin::GFP; unc-119(+)]* | |
| BA717 | *spe-11(hc90); sDp2(I,f)* | Reduction-of-function mutant allele of *spe-11* (D145E and W191Stop) |
| BS553 [a] | *fog-2(oz40)* V | |
| CB3855 | *plg-1(e2001)* III*; him-5(e1490)* V | |
| CGC43 | *unc-4(e120)/mnC1[dpy-10(e128) unc-52(e444) umnIs32]* II | Balancer chromosome marked with *myo-2::GFP* |
| DG4746 | *oops-1(tm6141)/tmC25[unc-5(tmIs1241)]* IV | Deletion allele of *oops-1,* removes 622 bp (see Fig. S1) |
| DG4753 | *oops-1(tm6141)/tmC25[unc-5(tmIs1241)]* IV*; him-5(e1490)* V | |
| DG4763 [a] | *oops-1(tm6141)/tmC25[unc-5(tmIs1241)]* IV*; fog-2(oz40)* V | |
| DG4778 | *oops-1(tn1898)/tmC25[unc-5(tmIs1241)]* IV | Entire open reading frame deletion allele of *oops-1* |
| DG4800 | *oops-1(tn1908[gfp::tev::3xflag::oops-1a])* IV | N-terminal GFP fusion of *oops-1a/c* |
| DG4806 | *oops-1(tn1914[oops-1::gfp::tev::3xflag])* IV | C-terminal GFP fusion of *oops-1* |
| DG4826 | *lin-41(n2914)* I*/hT2[qIs48]* (I; III)*; oops-1(tn1908[gfp::tev::3xflag::oops-1a])* IV | |
| DG4833 | *oops-1(tn1898)/tmC25[unc-5(tmIs1241)]* IV*; asIs3[egg-1::gfp + unc-119(+)]* | |
| DG4835 | *oops-1(tn1898)/tmC25[unc-5(tmIs1241)]* IV*; asIs4[egg-2::gfp + unc-119(+)]* | |
| DG4837 | *oops-1(tn1898)/tmC25[unc-5(tmIs1241)]* IV*; asIs1[pie-1p::gfp::egg-3 + unc-119(+)]* | |
| DG4839 | *oops-1(tn1898)/tmC25(unc-5[tnIs1241])* IV*; axIs1140[pie-1p::gfp::mbk-2 + unc-119(+)]* | |
| DG4873 | *pmnSi5[perm-4p::perm-4::mCherry + unc-119(+)]* II | |
| DG4875 | *oops-1(tn1898)/tmC25(unc-5[tnIs1241])* IV*; avIs143 [cbd-1p::cbd-1::mCherry + unc-119(+)]* | |
| DG4877 | *pmnSi1[perm-2p::perm-2::mCherry + unc-119(+)]* II*; oops-1(tn1898)/tmC25[unc-5(tmIs1241)]* IV | |

| | | |
|---|---|---|
| DG4879 | *pmnSi5[perm-4p::perm-4::mCherry + unc-119(+)]* II*; oops-1(tn1898)/tmC25[unc-5(tmIs1241)]* IV | |
| DG4883 | *his-72(uge30[gfp::his-72])* III*; oops-1(tn1898)/tmC25[unc-5(tmIs1241)]* IV | |
| DG4890 | *oops-1(tn1927[gfp::tev::3xflag::oops-1a]) oma-1(zu405te33)* IV | |
| DG4904 | *oops-1(tn1927[gfp::tev::3xflag::oops-1a]) oma-1(zu405te33)/nT1[qIs51]* IV*; oma-2(te51)/nT1[qIs51]* V | |
| DG4912 [a] | *oops-1(tn1908[gfp::tev::3xflag::oops-1a])* IV*; fog-2(oz40)* V | |
| DG4915 [a] | *his-72(uge30[gfp::his-72])* III*; fog-2(oz40)* V | |
| DG4917 | *lin-41(n2914)/tmC18[dpy-5(tmIs1236)]* I*; oops-1(tn1927[gfp::tev::3xflag::oops-1a]) oma-1(zu405te33)/nT1[qIs51]* IV*; oma-2(te51)/nT1[qIs51]* V | |
| DG4954 | *oops-1(tn1898)/tmC25[unc-5(tmIs1241)]* IV*; him-5(e1490)* V | |
| DG4956 [a] | *his-72(uge30[gfp::his-72])* III*; oops-1(tn1898)/tmC25[unc-5(tmIs1241)]* IV*; fog-2(oz40)* V | |
| DG4984 | *spe-11(ok2143)/tmC18[dpy-5[tmIs1236)]* I | Strong loss-of-function *spe-11* allele, 1x backcrossed |
| DG4985 | *spe-11(hc90)/tmC18[dpy-5[tmIs1236)]* I | |
| DG4993 | *oops-1(tn1908[gfp::tev::3xflag::oops-1a] tn1935)* IV | Deletion A (GFP::OOPS-1 Δ1–102 bp *3'UTR*) in Fig. 6A |
| DG4997 | *oops-1(tn1908[gfp::tev::3xflag::oops-1a] tn1938)* IV | Deletion B (GFP::OOPS-1 Δ103–203 bp *3'UTR*) in Fig. 6A |
| DG4999 | *oops-1(tn1908[gfp::tev::3xflag::oops-1a] tn1940)* IV | Deletion C (GFP::OOPS-1 Δ204–358 bp *3'UTR*) in Fig. 6A |
| DG5005 | *oops-1(tn1908[gfp::tev::3xflag::oops-1a] tn1944)* IV | Deletion D (GFP::OOPS-1 Δ1–203 bp *3'UTR*) in Fig. 6A |
| DG5009 | *oops-1(tn1908[gfp::tev::3xflag::oops-1a] tn1947)* IV | Deletion E (GFP::OOPS-1 Δ103–358 bp *3'UTR*) in Fig. 6A |
| DG5013 | *oops-1(tn1908[gfp::tev::3xflag::oops-1a])* IV*; him-5(e1490)* V | |
| DG5017 | *oops-1(tn1908[gfp::tev::3xflag::oops-1a] tn1949*ts*)* IV | Deletion F (GFP::OOPS-1 Δ1–358 bp *3'UTR*) in Fig. 6A |
| DG5033 | *oops-1(tn1908[gfp::tev::3xflag::oops-1a] tn1954)* IV | Deletion G (GFP::OOPS-1 ΔM1–V28) in Fig. 6A |
| DG5037 | *oops-1(tn1908[gfp::tev::3xflag::oops-1a] tn1957)* IV | Deletion H (GFP::OOPS-1 ΔK29–N65) in Fig. 6A |
| DG5045 | *oops-1(tn1908[gfp::tev::3xflag::oops-1a] tn1961)* IV | Deletion I (GFP::OOPS-1 ΔG66–S109) in Fig. 6A |
| DG5047 | *oops-1(tn1908[gfp::tev::3xflag::oops-1a] tn1963)* IV | Deletion J (GFP::OOPS-1 ΔG110–I138) in Fig. 6A |
| DG5049 | *oops-1(tn1908[gfp::tev::3xflag::oops-1a] tn1965)/tmC25[unc-5(tmIs1241)]* IV | Deletion K (GFP::OOPS-1 ΔT139–M201) in Fig. 6A |
| DG5051 | *oops-1(tn1908[gfp::tev::3xflag::oops-1a] tn1967)/tmC25[unc-5(tmIs1241)]* IV | Deletion L (GFP::OOPS-1 ΔI202–S209) in Fig. 6A |
| DG5053 | *oops-1(tn1908[gfp::tev::3xflag::oops-1a] tn1969)/tmC25[unc-5(tmIs1241)]* IV | Deletion M (GFP::OOPS-1 ΔR210–F238) in Fig. 6A |
| DG5055 | *oops-1(tn1908[gfp::tev::3xflag::oops-1a] tn1971)/tmC25[unc-5(tmIs1241)]* IV | Deletion N (GFP::OOPS-1 ΔK239–Y284) in Fig. 6A |
| DG5065 | *oops-1(tn1908[gfp::tev::3xflag::oops-1a] tn1975)/tmC25[unc-5(tmIs1241)]* IV | Deletion O (GFP::OOPS-1 ΔV285–V328) in Fig. 6A |
| DG5067 | *oops-1(tn1908[gfp::tev::3xflag::oops-1a] tn1977)* IV | Deletion P (GFP::OOPS-1 ΔT329–P374) in Fig. 6A |
| DG5097 | *oops-1(tn1908[gfp::tev::3xflag::oops-1a] tn1983)* IV | Deletion Q (GFP::OOPS-1 ΔL375–P416) in Fig. 6A |
| DG5099 | *oops-1(tn1908[gfp::tev::3xflag::oops-1a] tn1985)* IV | Deletion R (GFP::OOPS-1 ΔL417–S474) in Fig. 6A |
| DG5101 | *oops-1(tn1908[gfp::tev::3xflag::oops-1a] tn1987)* IV | Deletion S (GFP::OOPS-1 ΔL417–D443) in Fig. 6A |
| DG5103 | *oops-1(tn1908[gfp::tev::3xflag::oops-1a] tn1989)* IV | Deletion T (GFP::OOPS-1 ΔG444–S474) in Fig. 6A |

| | | |
|---|---|---|
| DG5105 | *oops-1(tn1908[gfp::tev::3xflag::oops-1a] tn1991)/tmC25[unc-5(tmIs1241)]* IV | Deletion V (GFP::OOPS-1 ΔT139–G156) in Fig. 6A |
| DG5107 | *oops-1(tn1908[gfp::tev::3xflag::oops-1a] tn1993)/tmC25[unc-5(tmIs1241)]* IV | Deletion W (GFP::OOPS-1 ΔP157–M201) in Fig. 6A |
| DG5137 | *oops-1(tn1908[gfp::tev::3xflag::oops-1a] tn1995)* IV | Deletion X (GFP::OOPS-1 ΔV475–A498) in Fig. 6A |
| DG5139 | *oops-1(tn1908[gfp::tev::3xflag::oops-1a] tn1997)* IV | Deletion Y (GFP::OOPS-1 ΔA499–L534) in Fig. 6A |
| DG5142 | *oops-1(tn1908[gfp::tev::3xflag::oops-1a] tn2000)* IV | Deletion U (GFP::OOPS-1 ΔM1–R103) in Fig. 6A |
| DG5147 | *oops-1(tn1908[gfp::tev::3xflag::oops-1a] tn2003)/tmC25[unc-5(tmIs1241)]* IV | Deletion Z (GFP::OOPS-1 ΔM1–L534) in Fig. 6A |
| DG5172 | *oops-1(tn1908[gfp::tev::3xflag::oops-1a] tn2013)* IV | Deletion GJ (GFP::OOPS-1 ΔM1–I138) in Fig. 6A |
| DG5177 | *oops-1(tn1908[gfp::tev::3xflag::oops-1a] tn2017)/tmC25[unc-5(tmIs1241)]* IV | Deletion P1Y (GFP::OOPS-1 ΔT329–L534) in Fig. 6A |
| DG5187 | *oops-1(tn1908[gfp::tev::3xflag::oops-1a] tn2023)* IV | Deletion QY (GFP::OOPS-1 ΔL375–L534) in Fig. 6A |
| DG5189 | *oops-1(tn1908[gfp::tev::3xflag::oops-1a] tn2025)/tmC25[unc-5(tmIs1241)]* IV | Deletion VO (GFP::OOPS-1 ΔT139–V328) in Fig. 6A |
| DG5195 | *oops-1(tn1908[gfp::tev::3xflag::oops-1a] tn2013 tn2027)* IV | Deletion GJ+QY (GFP::OOPS-1 ΔM1–I138 and ΔL375–L534) in Fig. 6A |
| DG5217 | *oops-1(tn1908[gfp::tev::3xflag::oops-1a] tn2013 tn2037)/tmC25[unc-5(tmIs1241)]* IV | Deletion GJ+P1Y (GFP::OOPS-1 ΔM1–I138 and ΔT329–L534) in Fig. 6A, GFP::OOPS-1 Mini expression shown in Fig. S5 |
| DG5219 | *oops-1(tn1908[gfp::tev::3xflag::oops-1a] tn2039)* IV | Deletion P2Y (GFP::OOPS-1 ΔL350–L534) in Fig. 6A |
| DG5221 | *oops-1(tn1908[gfp::tev::3xflag::oops-1a] tn2013 tn2041)* IV | Deletion GJ+P2Y (GFP::OOPS-1 ΔM1–I138 and ΔL350–L534) in Fig. 6A. Expresses GFP::OOPS-1 Mini. Expression shown in Fig. S5 |
| DG5223 | *oops-1(tn1908[gfp::tev::3xflag::oops-1a] tn2043)* IV | Deletion P1 (GFP::OOPS-1 ΔT329–C349) in Fig. 6A |
| DG5225 | *oops-1(tn1908[gfp::tev::3xflag::oops-1a] tn2045)* IV | Deletion P2 (GFP::OOPS-1 ΔL350–P374) in Fig. 6A |
| DG5227 | *oops-1(tn1908[gfp::tev::3xflag::oops-1a] tn2047)/tmC25[unc-5(tmIs1241)]* IV | Deletion OP1 (GFP::OOPS-1 ΔV285–C349) in Fig. 6A |
| DG5237 | *oops-1(tn1908[gfp::tev::3xflag::oops-1a] tn2051)/tmC25[unc-5(tmIs1241)]* IV | Deletion VP1 (GFP::OOPS-1 ΔT139–C349) in Fig. 6A |
| DG5266 | *spe-11(tn2059)/tmC18[dpy-5[tmIs1236)]* I | Entire open reading frame deletion allele of *spe-11* |
| DG5275 | *spe-11(ok2213)/tmC18[dpy-5[tmIs1236)]* I | Strong loss-of-function *spe-11* allele, 1x backcrossed |
| DG5295 | *spe-11(tn2068[mScarlet::tev::3xflag::spe-11])* I | N-terminal mScarlet fusion of *spe-11* |
| DG5305 | *spe-11(tn2068[mScarlet::tev::3xflag::spe-11])* I*; oops-1(tn1908[gfp::tev::3xflag::oops-1a])* IV | Complementary expression strain of mScarlet::SPE-11 and GFP::OOPS-1A in Fig. 4 |
| DG5307 | *spe-11(tn2068[mScarlet::tev::3xflag::spe-11])* I*; oops-1(tn1914[oops-1::gfp::tev::3xflag])* IV | Expresses OOPS-1::GFP (tagged at the C-terminus) and mScarlet::SPE-11. Expression shown in Fig. S5 |
| DG5321 | *spe-11(tn2059)/tmC18[dpy-5(tmIs1236)]* I*; him-5(e1490)* V | |
| DG5327 | *tnSi5(Cbr-unc-119(+) + mex-5p::mScarlet::3xflag::spe-11::spe-11 3'utr (65 bp) + 318-bp downstream)* II | Ectopic expression of mScarlet:: SPE-11 in oocytes using the *mex-5* promoter and the *spe-11 3'utr* |
| DG5328 | *tnSi6(Cbr-unc-119(+) + mex-5p::mScarlet::3xflag::spe-11::spe-11 3'utr (65 bp) + 318-bp downstream)* II | Ectopic expression of mScarlet:: SPE-11 in oocytes using the *mex-5* promoter and the *spe-11 3'utr* |
| DG5358 [a] | *oops-1(tn1898)/tmC25[unc-5(tmIs1241)]* IV*; fog-2(oz40)* V | |
| DG5368 | *spe-11(tn2059)* I*; tnSi5 (Cbr-unc-119(+) + mex-5p::mScarlet::3xflag::spe-11::spe-11 3'utr)* II | |
| DG5370 | *tnSi5(Cbr-unc-119(+) + mex-5p::mScarlet::3xflag::spe-11::spe-11 3'utr)* II*; oops-1(tn1898)/tmC25[unc-5(tmIs1241)]* IV | |
| DG5372 | *tnSi5(Cbr-unc-119(+) + mex-5p::mScarlet::3xflag::spe-11::spe-11 3'utr)* II*; oops-1(tn1908[gfp::tev::3xflag::oops-1a])* IV | |

| | | |
|---|---|---|
| DG5376 [a] | *tnSi5(Cbr-unc-119(+) + mex-5p::mScarlet::3xflag::spe-11::spe-11 3'utr)* II*; fog-2(oz40)* V | |
| DG5382 | *spe-11(tn2059)* I*; tnSi6 (Cbr-unc-119(+) + mex-5p::mScarlet::3xflag::spe-11::spe-11 3'utr)* II | |
| DG5390 [a] | *tnSi6(Cbr-unc-119(+) + mex-5p::mScarlet::3xflag::spe-11::spe-11 3'utr)* II*; fog-2(oz40)* V | |
| DG5430 | *spe-11(tn2094[gfp::tev::3xflag::spe-11])* I | N-terminal GFP fusion of *spe-11* |
| DG5462 | *spe-11(tn2094[gfp::tev::3xflag::spe-11])* I*; him-5(e1490)* V | |
| DG5479 | *tnSi10(Cbr-unc-119p::unc-119(+) + spe-11p::gfp::3xflag::oops-1a::spe-11 3'utr + 318-bp downstream)* II | Ectopic expression of GFP::OOPS-1A in sperm using the *spe-11* promoter and *3'utr* |
| DG5495 | *tnSi10(Cbr-unc-119p::unc-119(+) + spe-11p::gfp::3xflag::oops-1a::spe-11 3'utr + 318-bp downstream)* II*; oops-1(tn1898)/tmC25[unc-5(tmIs1241)]* IV | |
| DG5518 | *tnSi10(Cbr-unc-119p::unc-119(+) + spe-11p::gfp::3xflag::oops-1a::spe-11 3'utr + 318-bp downstream)* II*; him-5(e1490)* V | |
| DG5544 | *tnSi12(Cbr-unc-119p::unc-119(+) + peel-1p::gfp::3xflag::oops-1a::peel-1 3'utr + 682-bp downstream)* II | Ectopic expression of GFP::OOPS-1A in sperm using the *peel-1* promoter and *3'utr* |
| DG5546 | *tnSi12(Cbr-unc-119p::unc-119(+) + peel-1p::gfp::3xflag::oops-1a::peel-1 3'utr + 682-bp downstream)* II*; oops-1(tn1898)/tmC25[unc-5(tmIs1241)]* IV | |
| DG5550 | *tnSi12(Cbr-unc-119p::unc-119(+) + peel-1p::gfp::3xflag::oops-1a::peel-1 3'utr + 682-bp downstream)* II*; him-5(e1490)* V | |
| DG5554 | *tnSi14(Cbr-unc-119p::unc-119(+) + trp-3p::gfp::3xflag::oops-1a::trp-3 3'utr + 1590-bp downstream)* II | Ectopic expression of GFP::OOPS-1A in sperm using the *trp-3* promoter and *3'utr* |
| DG5556 | *tnSi14(Cbr-unc-119p::unc-119(+) + trp-3p::gfp::3xflag::oops-1a::trp-3 3'utr + 1590-bp downstream)* II*; oops-1(tn1898)/tmC25[unc-5(tmIs1241)]* IV | |
| DG5594 | *spe-11(tn2068[mScarlet::tev::3xflag::spe-11])* I*; oops-1 (tn1908 tn2000)* IV | Expresses GFP::OOPS-1B and mScarlet::SPE-11. Expression shown in Fig. S5 |
| DG5560 | *tnSi14(Cbr-unc-119p::unc-119(+) + trp-3p::gfp::3xflag::oops-1a::trp-3 3'utr + 1590-bp downstream)* II*; him-5(e1490)* V | |
| DG5610 | *spe-11(tn2059)/tmC18* I*; ltIs37[pie-1p::mCherry::his-58 + unc-119(+)]* IV*; ruIs57[pie-1::gfp::tba-2 + unc-119(+)]* | |
| DG5611 | *tnSi16(Cbr-unc-119p::unc-119(+) + spe-11p::gfp::3xflag::oops-1b::spe-11 3'utr + 318-bp downstream)* II | Ectopic expression of GFP::OOPS-1B in sperm using the *spe-11* promoter and *3'utr* |
| DG5613 | *tnSi16(Cbr-unc-119p::unc-119(+) + spe-11p::gfp::3xflag::oops-1b::spe-11 3'utr + 318-bp downstream)* II*; oops-1(tn1898)/tmC25[unc-5(tmIs1241)]* IV | |
| DG5617 | *tnSi16(Cbr-unc-119p::unc-119(+) + spe-11p::gfp::3xflag::oops-1b::spe-11 3'utr + 318-bp downstream)* II*; him-5(e1490)* V | |
| DG5627 | *spe-11(hc90)/tmC18[dpy-5(tmIs1236)]* I*; oops-1(tn1898)/tmC25[unc-5(tmIs1241)]* IV*; ruIs57; ltIs37* | |
| DG5643 | *spe-11(tn2094[gfp::tev::3xflag::spe-11] tn2139)* I | Deletion A (GFP:: SPE-11 ΔM1–N14) in Fig. 6D |
| DG5645 | *spe-11(tn2094[gfp::tev::3xflag::spe-11] tn2141)* I | Deletion B (GFP:: SPE-11 ΔK15–S29) in Fig. 6D |
| DG5647 | *spe-11(tn2094[gfp::tev::3xflag::spe-11] tn2143)* I | Deletion C (GFP:: SPE-11 ΔQ30–Y43) in Fig. 6D |
| DG5649 | *spe-11(tn2094[gfp::tev::3xflag::spe-11] tn2145*ts*)* I | Deletion D (GFP:: SPE-11 ΔP44–S57) in Fig. 6D |
| DG5651 | *spe-11(tn2094[gfp::tev::3xflag::spe-11] tn2146)* I | Deletion E (GFP:: SPE-11 ΔD58–K72) in Fig. 6D |

| | | |
|---|---|---|
| DG5653 | *spe-11(tn2094[gfp::tev::3xflag::spe-11] tn2147)* I | Deletion F (GFP:: SPE-11 ΔK73–D87) in Fig. 6D |
| DG5655 | *spe-11(tn2094[gfp::tev::3xflag::spe-11] tn2148)/tmC18[dpy-5[tmIs1236)]* I | Deletion G (GFP:: SPE-11 ΔM88–F101) in Fig. 6D |
| DG5657 | *spe-11(tn2094[gfp::tev::3xflag::spe-11] tn2150)/tmC18[dpy-5[tmIs1236)]* I | Deletion H (GFP:: SPE-11 ΔL102–A115) in Fig. 6D |
| DG5659 | *spe-11(tn2094[gfp::tev::3xflag::spe-11] tn2152)/tmC18[dpy-5[tmIs1236)]* I | Deletion I (GFP:: SPE-11 ΔR116–G129) in Fig. 6D |
| DG5661 | *spe-11(tn2094[gfp::tev::3xflag::spe-11] tn2154)/tmC18[dpy-5[tmIs1236)]* I | Deletion J (GFP:: SPE-11 ΔF130–N144) in Fig. 6D |
| DG5663 | *spe-11(tn2094[gfp::tev::3xflag::spe-11] tn2156)/tmC18[dpy-5[tmIs1236)]* I | Deletion K (GFP:: SPE-11 ΔD145–A158) in Fig. 6D |
| DG5685 | *spe-11(tn2094[gfp::tev::3xflag::spe-11] tn2160)/tmC18[dpy-5[tmIs1236)]* I | Deletion L (GFP:: SPE-11 ΔT159–M172) in Fig. 6D |
| DG5687 | *spe-11(tn2094[gfp::tev::3xflag::spe-11] tn2161)/tmC18[dpy-5[tmIs1236)]* I | Deletion M (GFP:: SPE-11 ΔE173–A185) in Fig. 6D |
| DG5689 | *spe-11(tn2094[gfp::tev::3xflag::spe-11] tn2163)/tmC18[dpy-5[tmIs1236)]* I | Deletion N (GFP:: SPE-11 ΔR186–L200) in Fig. 6D |
| DG5718 | *spe-11(tn2059)/tmC18[dpy-5[tmIs1236)]* I*; oops-1(tn1898)/tmC25[unc-5(tmIs1241)]* IV | |
| DG5723 | *spe-11(tn2094[gfp::tev::3xflag::spe-11] tn2169)/tmC18[dpy-5[tmIs1236)]* I | Deletion O (GFP:: SPE-11 ΔA201–D215) in Fig. 6D |
| DG5725 | *spe-11(tn2094[gfp::tev::3xflag::spe-11] tn2171)/tmC18[dpy-5[tmIs1236)]* I | Deletion P (GFP:: SPE-11 ΔA216–E230) in Fig. 6D |
| DG5738 | *spe-11(tn2094[gfp::tev::3xflag::spe-11] tn2175)/tmC18[dpy-5[tmIs1236)]* I | Deletion R (GFP:: SPE-11 ΔL246–K258) in Fig. 6D |
| DG5743 | *spe-11(tn2094[gfp::tev::3xflag::spe-11] tn2178)/tmC18[dpy-5[tmIs1236)]* I | Deletion T (GFP:: SPE-11 ΔD272–R285) in Fig. 6D |
| DG5745 | *spe-11(tn2094[gfp::tev::3xflag::spe-11] tn2180)/tmC18[dpy-5[tmIs1236)]* I | Deletion U (GFP:: SPE-11 ΔN286–K299) in Fig. 6D |
| DG5747 | *spe-11(tn2094[gfp::tev::3xflag::spe-11] tn2181)/tmC18[dpy-5[tmIs1236)]* I | Deletion AF (GFP:: SPE-11 ΔM1–D87) in Fig. 6D |
| DG5749 | *spe-11(tn2094[gfp::tev::3xflag::spe-11] tn2183)* I | Deletion W (GFP:: SPE-11 ΔR295–K299) in Fig. 6D |
| DG5759 | *spe-11(tn2094[gfp::tev::3xflag::spe-11] tn2184)/tmC18[dpy-5[tmIs1236)]* I | Deletion AE (GFP:: SPE-11 ΔM1–K72) in Fig. 6D |
| DG5762 | *spe-11(tn2094[gfp::tev::3xflag::spe-11] tn2185)/tmC18[dpy-5[tmIs1236)]* I | Deletion Q (GFP:: SPE-11 ΔL231–W245) in Fig. 6D |
| DG5764 | *spe-11(tn2094[gfp::tev::3xflag::spe-11] tn2187)/tmC18[dpy-5[tmIs1236)]* I | Deletion S (GFP:: SPE-11 ΔF259–E271) in Fig. 6D |
| DG5780 | *chs-1(tn2195) spe-11(tn2094[gfp::tev::3xflag::spe-11] tn2145*ts*)* I | 1x outcrossed using *tmC18/+* males |
| DG5783 | *chs-1(tn2192) spe-11(tn2094[gfp::tev::3xflag::spe-11] tn2145*ts*)* I | 1x outcrossed using *tmC18/+* males |
| DG5829 | *chs-1(tn2201) spe-11(tn2094[gfp::tev::3xflag::spe-11] tn2145*ts*)* I | 7x outcrossed using *tmC18/+* males |
| DG5830 | *chs-1(tn2198) spe-11(tn2094[gfp::tev::3xflag::spe-11] tn2145*ts*)* I | 7x outcrossed using *tmC18/+* males |
| DG5837 | *chs-1(tn2191) spe-11(tn2094[gfp::tev::3xflag::spe-11] tn2145*ts*)* I | 7x outcrossed using *tmC18/+* males |
| DG5842 | *gsp-3(tn2202) spe-11(tn2094[gfp::tev::3xflag::spe-11] tn2145*ts*)* I | 8x outcrossed using *tmC18/+* males |
| DG5843 | *spe-11(tn2094[gfp::tev::3xflag::spe-11] tn2145*ts*)* I; *egg-3(tn2205)* II | 7x outcrossed using *tmC18/+* males |
| DG5845 | *chs-1(tn2189) spe-11(tn2094[gfp::tev::3xflag::spe-11] tn2145*ts*)* I | 7x outcrossed using *tmC18/+* males |
| DG5847 | *chs-1(tn2210) spe-11(tn2094[gfp::tev::3xflag::spe-11] tn2145*ts*)* I | 3x outcrossed using *tmC18/+* males |
| DG5833 | *chs-1(tn2210) spe-11(tn2094[gfp::tev::3xflag::spe-11] tn2145*ts*)* I | 7x outcrossed using *tmC18/+* males |
| DG5855 | *tnEx265[str-1p::gfp]* | |
| DG5866 | *tnSi21(Cbr-unc-119p::unc-119(+) + oma-1p::gfp::3xflag::spe-11::oma-1 3'utr)* II | Ectopic expression of GFP:: SPE-11 in oocytes using the *oma-1* promoter and *3'utr* |
| DG5868 | *tnSi23(Cbr-unc-119p::unc-119(+) + oma-2p::gfp::3xflag::spe-11::oma-2 3'utr)* II | Ectopic expression of GFP:: SPE-11 in oocytes using the *oma-2* promoter and *3'utr* |
| DG5870 | *tnSi25(Cbr-unc-119p::unc-119(+) + rme-2p::gfp::3xflag::spe-11::rme-2 3'utr)* II | Ectopic expression of GFP:: SPE-11 in oocytes using the *rme-2* promoter and *3'utr* |
| DG5894 | *tnSi27(Cbr-unc-119p::unc-119(+) + puf-5p::gfp::3xflag::spe-11::puf-5 3'utr)* II | Ectopic expression of GFP:: SPE-11 in oocytes using the *puf-5* promoter and *3'utr* |
| DG5913 | *chs-1(ok1120)/tmC18[dpy-5[tmIs1236)]* I*; oops-1(tn1908[gfp::tev::3xflag::oops-1a])* IV | |

| | | |
|---|---|---|
| DG5915 | egg-3(tm1191)/mnC1[dpy-10(e128) unc-52(e444) umnIs32] II; oops-1(tn1908[gfp::tev::3xflag::oops-1a]) IV | |
| DG5917 | egg-3(ok3651)/mnC1[dpy-10(e128) unc-52(e444) umnIs32] II; oops-1(tn1908[gfp::tev::3xflag::oops-1a]) IV | |
| DG5927 | oops-1(gk503838) IV | 1x backcrossed |
| DG5933 | chs-1(tn2298) spe-11(tn2094[gfp::tev::3xflag::spe-11] tn2145ts) I | 1x backcrossed |
| DG5936 | spe-11(tn2059) I; tnSi21(Cbr-unc-119p::unc-119(+) + oma-1p::gfp::3xflag::spe-11::oma-1 3'utr) II | |
| DG5938 | spe-11(tn2059) I; tnSi23(Cbr-unc-119p::unc-119(+) + oma-2p::gfp::3xflag::spe-11::oma-2 3'utr) II | |
| DG5942 | spe-11(tn2059) I; tnSi25(Cbr-unc-119p::unc-119(+) + rme-2p::gfp::3xflag::spe-11::rme-2 3'utr) II | |
| DG5959 | spe-11(tn2059)/tmC18[dpy-5[tmIs1236)] I; tnSi5/mnC1[dpy-10(e128) unc-52(e444) umnIs32] II | |
| DG5970 | spe-11(tn2059) I; tnSi5(Cbr-unc-119(+) + mex-5p::mScarlet::3xflag::spe-11::spe-11 3'utr) II | |
| DG6013 | chs-1(tn2242) spe-11(tn2094[gfp::tev::3xflag::spe-11] tn2145ts) I | 7x outcrossed using tmC18/+ males |
| DG6032 | spe-11(tn2094[gfp::tev::3xflag::spe-11] tn2145ts) I; egg-3(tn2190) II | 7x outcrossed using tmC18/+ males |
| DG6035 | spe-11(tn2059) I; tnSi27(Cbr-unc-119p::unc-119(+) + puf-5p::gfp::3xflag::spe-11::puf-5 3'utr) II | |
| EG6699 | ttTi5605 II; unc-119(ed3) III; oxEx1578[eft-3p::gfp + Cbr-unc-119(+)] | |
| FAS46 | his-72(uge30[gfp::his-72]) II | |
| FM125 | unc-119(ed3) III; ruIs57[pAZ147:pie-1::β-tubulin::GFP; unc-119(+)]; ltIs37 [unc-119(+) pie-1::mCherry::H2B]) IV | |
| FX01031 [b] | rog-1(tm1031)/+ II | |
| FX01600 [b] | Y54F10AR.1(tm1600)/+ III | |
| FX02902 [b] | Y60A9.3(tm2902)/+ X | |
| FX02920 [b] | F54D5.5(tm2920)/+ II | |
| FX03459 [b] | E02H1.5(tm3459)/+ II | |
| FX03823 [b] | tmed-4(tm3823)/+ V | |
| FX04390 [b] | R148.3(tm4390)/+ III | |
| FX05077 [b] | fbxl-2(tm5077)/+ III | |
| FX05958 [b] | tmem-184(tm5958)/+ X | |
| FX06117 [b] | tmem-131(tm6117)/+ III | |
| FX06123 [b] | C55A6.11(tm6123)/+ V | |
| FX06315 [b] | H18N23.2(tm6315)/+ X | |
| FX06697 [b] | B0416.5(tm6697)/+ X | |
| FX06735 [b] | B0395.3(tm6735)/+ X | |
| FX07138 [b] | garr-1(tm7138)/+ IV | |
| FX14551 [b] | atx-2(tm3562) III/hT2[bli-4(e937) let-?(q782) qIs48] (I; III) | |
| FX14556 [b] | atx-2(tm4373) III/hT2[bli-4(e937) let-?(q782) qIs48] (I; III) | |
| FX14680 [b] | ept-1(tm3093) I/hT2[bli-4(e937) let-?(q782) qIs48] (I; III) | |
| FX14731 [b] | pygl-1(tm5211) V/nT1[qIs51] (IV; V) | |

| | | |
|---|---|---|
| FX14783 [b] | *F26E4.7(tm3127)* I/*hT2[bli-4(e937) let-?(q782) qIs48]* (I; III) | |
| FX14825 [b] | *E02D9.1(tm5862)* I/*hT2[bli-4(e937) let-?(q782) qIs48]* (I; III) | |
| FX14836 [b] | *oops-1(tm6141)* IV/*nT1[qIs51]* (IV; V) | Deletion allele of *oops-1,* removes 622 bp (see Fig. S1) |
| FX14861 [b] | *pigm-1(tm3400)/mIn1[mIs14 dpy-10(e128)]* II | |
| FX14877 [b] | *T05H10.1(tm5233)/mIn1[mIs14 dpy-10(e128)]* II | |
| FX14889 [b] | *R186.3(tm5639)* V/*nT1[qIs51]* (IV; V) | |
| FX15076 [b] | *pigg-1(tm2377)/dpy-10(e128)* II | |
| FX16511 [b] | *W04D2.4(tm586)* V/*nT1[qIs51]* (IV; V) | |
| FX16640 [b] | *C06A5.3(tm5259)* I/*hT2[bli-4(e937) let-?(q782) qIs48]* (I; III) | |
| FX16751 [b] | *rbmx-2(tm6741)/mIn1[mIs14 dpy-10(e128)]* II | |
| FX16752 [b] | *metl-13(tm6756)* IV/*nT1[qIs51]* (IV; V) | |
| FX16798 [b] | *algn-9(tm3297)/+* II | |
| FX16946 [b] | *efr-3(tm862)/mIn1[mIs14 dpy-10(e128)]* II | |
| FX17004 [b] | *T26C5.3(tm1317)/unc-4(e120)* II | |
| FX17310 [b] | *B0025.4(tm6909)* I/*hT2[bli-4(e937) let-?(q782) qIs48]* (I; III) | |
| FX17625 [b] | *znf-706(tm4820)* IV/*nT1[qIs51]* (IV; V) | |
| FX17944 [b] | *E02D9.1(tm6625)* I/*hT2[bli-4(e937) let-?(q782) qIs48]* (I; III) | |
| FX17951 [b] | *Y110A7A.7(tm6680)* I/*hT2[bli-4(e937) let-?(q782) qIs48]* (I; III) | |
| FX17983 [b] | *rfk-1(tm7119)* IV/*nT1[qIs51]* (IV; V) | |
| FX18183 [b] | *nfs-1(tm3516)* I/*hT2[bli-4(e937) let-?(q782) qIs48]* (I; III) | |
| FX18190 [b] | *mjl-1(tm1651)* I/*hT2[bli-4(e937) let-?(q782) qIs48]* (I; III). | |
| FX18202 [b] | *R186.3(tm5542)* V/*nT1[qIs51]* (IV; V) | |
| FX18296 [b] | *F19B6.1(tm2376)* IV/*nT1[qIs51]* (IV; V) | |
| FX18303 [b] | *C50C3.1(tm4185)* III/*hT2[bli-4(e937) let-?(q782) qIs48]* (I; III) | |
| FX18488 [b] | *Y57A10A.31(tm2349)/mnC1[dpy-10(e128) unc52(e444) nIs190 let-?]* II | |
| FX18582 [b] | *C09D4.4(tm4773)* I/*hT2[bli-4(e937) let-?(q782) qIs48]* (I; III) | |
| FX19059 [b] | *sec-61.B(tm1986)/tmIn3* IV | |
| FX19152 [b] | *F56C11.3(tm6013)* I/*hT2[bli-4(e937) let-?(q782) qIs48]* (I; III) | |
| FX19197 [b] | *F56C11.3(tm5845)* I/*hT2[bli-4(e937) let-?(q782) qIs48]* (I; III) | |
| FX19220 [b] | *B0334.5(tm4576)/mIn1[mIs14 dpy-10(e128)]* II | |
| FX19232 [b] | *B0334.5(tm4759)/mIn1[mIs14 dpy-10(e128)]* II | |
| FX19256 [b] | *C32D5.8(tm5559)/mIn1[mIs14 dpy-10(e128)]* II | |
| FX19279 [b] | *C18E9.2(tm2236)/mIn1[mIs14 dpy-10(e128)]* II | |
| FX19298 [b] | *Y53C12B.1(tm2353)/mIn1[mIs14 dpy-10(e128)]* II | |
| FX19347 [b] | *ZK546.5(tm769)/mIn1[mIs14 dpy-10(e128)]* II | |
| FX19350 [b] | *F56D1.1(tm814)/mIn1[mIs14 dpy-10(e128)]* II | |
| FX19441 [b] | *F39H2.3(tm2016)* I/*hT2[bli-4(e937) let-?(q782) qIs48]* (I; III) | |
| FX20937 [b] | *tm10937*(I)*; wrn-1(tm764)* II*; polq-1(tm9609) exo-1(tm1842)* III*; C42C1.8 C42C1.12 (tm10938)/+* IV | |
| FX21247 [b] | *wrn-1(tm764)* II*; polq-1(tm9609) F25B5.6 F25B5.3 F25B5.7 F25B5.2 F25B5.9 etc (tm11247)/+ exo-1(tm1842)* III*; tm11249/tmC5[F36H1.3(tmIs1220) tm7167]* IV | |

| | | |
|---|---|---|
| FX30168 | *tmC18[dpy-5(tmIs1236)]* I | Balancer chromosome marked with *myo-2p::mCherry* |
| FX30203 | *tmC25[unc-5(tmIs1241)]* IV | Balancer chromosome marked with *myo-2p::Venus* |
| FX31218 [b] | *D2013.8 D2013.9 F42A8.1 F42A8.2 F42A8.3 C06C3.1(tm8373)/mIn1[mIs14 dpy-10(e128)]* II | |
| FX31257 [b] | *C55B6.4 C55B6.7 C55B6.2 C55B6.1 C55B6.5 ZK867.2 ZK867.3 ZK867.1 F46H5.3 (tm10686)/tmC30[ubc-17(tmIs1247)]* X | |
| FX31302 [b] | *ZK418.4 ZK418.5 ZK418.6 ZK418.7 ZK418.8 ZK418.9 ZK418.2 ZK418.1 B0280.5 B0280.6 B0280.4 (tm11024)* III | |
| FX31303 [b] | *K11D2.2 K11D2.3 K11D2.5 K11D2.4 K11D2.6 (tm11283)* I/*hT2[bli-4(e937) let-?(q782) qIs48]* (I; III) | |
| FX31304 [b] | *F35E12.8 F20G2.1 F20G2.9 F20G2.10 F20G2.2 F20G2.7 F20G2.6 F20G2.3 F20G2.4 F20G2.5 (tm11108)* V/*nT1[qIs51]* (IV; V) | |
| FX31307 [b] | *F32H2.9 F32H2.6 F36F2.7 F36F2.11 F36F2.4 F36F2.6 F36F2.3 F36F2.8 F36F2.2 F36F2.1 (tm11208)* I/*hT2[bli-4(e937) let-?(q782) qIs48]* (I; III). | |
| FX31308 [b] | *K08D10.9 K08D10.8 K08D10.7 K08D10.18 K08D10.5 K08D10.13 K08D10.4 K08D10.3 K08D10.2 K08D10.12 K08D10.1 K08D10.11 K06B9.3 K06B9.4 (tm11365)* IV/*nT1[qIs51]* (IV; V) | |
| FX31314 [b] | *C25A1.18 sup-46 C25A1.19 C25A1.17 C25A1.5 C25A1.16 C25A1.6 C25A1.7 C25A1.8 C25A1.9 C25A1.10 C25A1.12 C25A1.13 C25A1.11 C25A1.15 Y106G6E.1 (tm11402)* I/*hT2[bli-4(e937) let-?(q782) qIs48]* (I; III) | |
| FX31316 [b] | *B0350.2 B0350.74 B0350.83 C46G7.5 C46G7.111 C46G7.110 C46G7.2 C46G7.109 C46G7.1 (tm11129)* IV/*nT1[qIs51]* (IV; V) | |
| FX31331 [b] | *F12F6.1 F40F11.4 F40F11.3 F40F11.6 F40F11.2 F40F11.5 F40F11.1 Y24F12A.2 Y24F12A.1 tm11220*/+ *tm11221/tmC5 [F36H1.3(tmIs1220)]* IV | |
| FX31350 [b] | *tm10671*/+ *K10G6.3 Y14H12B.1 Y14H12B.2 (tm10672)*/+ *wrn-1(tm764)*/+ II | |
| FX31351 [b] | *T23B7.3 F11G11.7 F11G11.5 F11G11.8 F11G11.4 F11G11.9 F11G11.14 F11G11.10 F11G11.11 F11G11.12 F11G11.13 F11G11.3 F11G11.2 F11G11.1 R05F9.7 (tm10689)*/+ *wrn-1(tm764)*/+ II | |
| FX31352 [b] | *W08F4.16 W08F4.18 W08F4.17 W08F4.7 W08F4.8 W08F4.14 W08F4.3 W08F4.15 W08F4.9 W08F4.12 W08F4.2 W08F4.10 W08F4.11 W08F4.1 K07E8.6 K07E8.7 K07E8.5 K07E8.8 (tm11164)*/+ II | |
| FX31354 [b] | *T02H6.1 Y39F10B.1 C08G5.1 C08G5.2 C08G5.3 C08G5.4 (tm10816)*/+ *wrn-1(tm764)*/+ II | |
| FX31355 [b] | *tm10935*/+ *F57G8.6 W08G11.5 W08G11.1 W08G11.4 W08G11.3 W08G11.6 (tm10936)*/+ V | |
| FX31359 [b] | *wrn-1(tm764)*/+ *K09E4.3 algn-3 (tm10700)*/+ II | |
| FX31367 [b] | *Y54G2A.16(tm11276)*/+ IV | |
| FX31368 [b] | *Y41D4B.24 Y41D4B.28 K08D12.5 K08D12.4 K08D12.1 K08D12.3 K08D12.2 (tm11185)*/+ IV | |
| FX31369 [b] | *F42H10.5 F42H10.6 F42H10.14 F42H10.3 F42H10.7 F42H10.2 F42H10.16 F42H10.15 C04D8.1 F42H10.11 (tm8850)*/+ III | |
| FX31375 [b] | *B0212.1 Y37E11B.2 Y37E11B.1 Y37E11B.3 Y37E11B.4 Y37E11B.t1 Y37E11B.t2 (tm10723)*/+ IV | |

| | |
|---|---|
| FX31376 [b] | *Y71A12C.1 Y71A12C.4 Y71A12C.3 Y71A12C.2 F47G4.7 F47G4.8 F47G4.1 F47G4.9 F47G4.6 F47G4.5 F47G4.4 (tm11071)/+* I |
| FX31382 [b] | *W08G11.1 W08G11.4 W08G11.3 W08G11.6 (tm11313)/+* V |
| JH1576 | *unc-119(ed3)* III; *axIs1140 [pie1p::GFP::mbk-2 + unc-119(+)]* |
| POM1 | *pmnSi1[perm-2p::perm-2::mCherry + unc-119(+)]* II*; unc-119(ed3)* III |
| RB1189 | *chs-1(ok1120)* I/*hT2[bli-4(e937) let-?(q782) qIs48]* (I; III) |
| RT495 | *unc-119(ed3)* III*; asIs4[egg-2::gfp + unc-119(+)]* |
| RT497 | *unc-119(ed3)* III*; asIs3[egg-1::gfp + unc-119(+)]* |
| VC1128 [b] | *mis-12 Y47G6A.25(ok1536)/szT1[lon-2(e678)] I; +/szT1* X |
| VC1135 [b] | *R166.3(gk541)/mIn1[mIs14 dpy-10(e128)]* II |
| VC1259 [b] | *K05C4.2 K05C4.11(ok1713)/hIn1[unc-101(sy241)]* I |
| VC1825 [b] | *F44E2.8 F44E2.9(ok2134)* III/*hT2[bli-4(e937) let-?(q782) qIs48]* (I; III) |
| VC2135 [b] | *lelo-2(ok2740)/sC1[dpy-1(s2170)]* III |
| VC2238 [b] | *nfs-1(ok2890)* I/*hT2 [bli-4(e937) let-?(q782) qIs48]* (I; III) |
| VC2735 [b] | *+/mT1* II*; M142.5(ok3554)/mT1[dpy-10(e128)]* III |
| VC2840 [b] | *C24D10.4(ok3613)* IV/*nT1[qIs51]* (IV; V) |
| VC2876 | *egg-3(ok3651)/mIn1[mIs14 dpy-10(e128)]* II |
| VC2972 [b] | *R148.3(ok3525)/qC1[dpy-19(e1259) glp-1(q339)]* III |
| VC40188 | *oops-1(gk503838)* IV |

[a] Male-female strain.

[b] Deletion mutant strains annotated as sterile or lethal, which affect uncharacterized genes whose mRNA transcripts are associated with OMA-1 and/or LIN-41 (Tsukamoto et al., 2017). In strains containing multiple deletions, the genes that were identified as encoding OMA-1 and/or LIN-41-associated transcripts are underlined
